# ASCL1 primes medullary thymic epithelial cell programs required for central tolerance

**DOI:** 10.1101/2025.08.06.668799

**Authors:** Nobuko Akiyama, Kenta Horie, Kano Namiki, Jen-Chien Chang, Maki Miyauchi, Takahisa Miyao, Nanako Uchida, Naho Hagiwara, Masaki Yoshida, Mizuki Morota, Tsukasa Kouno, Miki Kojima, Jonathan Moody, Yoshinari Ando, Hirotake Ichise, Hiroto Ishii, Rin Endo, Wataru Muramatsu, Manli Yang, Nobuaki Yoshida, Jun-ichiro Inoue, Chung Chau Hon, Jay W. Shin, Piero Carninci, Georg A. Hollander, Aki Minoda, Jun Nakajima, Taishin Akiyama

## Abstract

Immunological self-tolerance depends on medullary thymic epithelial cells (mTECs), which express a broad repertoire of self-antigens to support negative selection of autoreactive T cells and the development of regulatory T cells. Although the autoimmune regulator AIRE is essential for this process, additional factors are required to establish the full mTEC gene expression program. Because thymoma is frequently associated with autoimmunity, implicating defective thymic tolerance, we performed single-cell RNA sequencing of TECs from thymoma patients and identified the transcription factor ASCL1 as selectively downregulated in tumor mTECs. Deletion of Ascl1 in mouse TECs resulted in spontaneous autoimmunity without inducing thymic tumorigenesis. Transcriptomic and chromatin accessibility analyses revealed that ASCL1 influences mTEC gene expression programs and is associated with corresponding changes in chromatin accessibility. Genetic interaction analyses further suggested that ASCL1 activity in immature mTECs modulates the AIRE dependency of gene expression in mature mTECs. Together, these findings identify ASCL1 as a key regulator of mTEC function and central tolerance, providing insight into mechanisms that safeguard immune homeostasis and whose disruption may contribute to autoimmune disease.

## Introduction

Thymic epithelial cells (TECs) are essential for establishing self-tolerant T cells. They comprise cortical (cTECs) and medullary (mTECs) subsets, which are distinct in both location and function: cTECs support early T cell development, while mTECs express thousands of tissue-specific self-antigens (TSAs) to mediate negative selection and promote regulatory T cell induction ^1^. Consequently, dysfunctional TECs cause immune deficiency and autoimmunity ^2^.

TSA expression in mTECs is transcriptionally regulated, with the autoimmune regulator (AIRE) playing a pivotal role ^3, 4^. However, many TSAs are AIRE-independent ^5^. Furthermore, single-cell analyses have revealed remarkable TEC heterogeneity in both humans and mice ^6, 7, 8, 9, 10, 11, 12, 13^, beyond traditional flow cytometric classifications based on expression of some marker molecules like MHC class II (MHC II), AIRE, and CCL21. Notably, subsets of mTECs activate “mimetic” gene programs resembling peripheral tissues, contributing to TSA diversity through AIRE-independent mechanisms ^12, 13^. However, the molecular mechanisms underlying AIRE-independent TSA expression within AIRE⁺ cells and the regulatory logic distinguishing AIRE-dependent from AIRE-independent control of specific TSAs remain poorly understood.

Further complexity arises from the existence of AIRE⁺ mTECs that include both “classical” TSA-expressing cells and transit-amplifying populations with low Aire-dependent gene expression ^11, 14^, suggesting that AIRE expression alone is insufficient to drive AIRE-dependent TSA expression and that TSA repertoire formation is tightly coupled to the mTEC differentiation program.

Thymomas, epithelial tumors of the thymus, commonly arise in adults and often retain the capacity to support thymopoiesis, generating immature thymocytes that subsequently differentiation into CD4⁺ and CD8⁺ T cell lineages ^15, 16^. Thymomas are strongly associated with autoimmune diseases, including myasthenia gravis (MG) and other forms of thymoma-associated autoimmunity ^17, 18, 19, 20^. Histologically, thymomas are classified into types A, AB, B1, B2, B3, and carcinoma ^21, 22^. Types AB, B1, and B2 typically contain abundant thymocytes, with type B2 showing the strongest association with autoimmunity ^20^. Although thymomas generally exhibit a low mutational burden^21^, *GTF2I* mutations are frequently found in type A and AB tumors. While these mutations can induce thymoma-like phenotypes in mice, they do not recapitulate the autoimmune manifestations ^23^.

Several non-mutually exclusive mechanisms have been proposed to explain thymoma-associated autoimmunity ^24^. Reduced AIRE expression in thymomas may compromise central tolerance, leading to the escape of autoreactive T cells and impaired regulatory T cell development ^20, 25, 26^. However, the autoimmune manifestations observed in thymoma-associated autoimmunity differ from those in APECED ^27, 28^, implying that additional alterations in thymoma TECs contribute to disease pathogenesis. Comparative studies of thymomas from MG and non-MG patients have revealed increased expression of muscle-specific antigens in MG-associated thymomas ^22, 29^, but, beyond this, expression of genes with a role in immunity and tolerance induction does not differ substantially between MG and non-MG thymomas ^22^. Thus, the molecular basis of thymoma-associated autoimmunity remains unclear.

ASCL1 acts as a pioneer factor in neuronal and lung neuroendocrine differentiation ^30^ and is expressed in TECs in both mice and humans ^10, 31, 32, 33^. It has been proposed to promote a neuroendocrine-like mTEC (Endo mTEC) subset ^13, 33^ and to mark human mTEC progenitors ^32^, although mouse fate-mapping studies challenge this ^10^. Thus, the precise roles of ASCL1 in mTEC gene regulation, immune tolerance, and human disease remain to be elucidated.

Given the limited transcriptional differences between MG and non-MG thymomas ^22^, we hypothesized that, relative to normal TECs, tumor-derived TECs may acquire immunogenic properties that compromise self-tolerance. Comparing gene expression profiles of tumor TECs and normal TECs from thymoma patients, we identified ASCL1 as significantly downregulated in tumor mTECs. In mice, TEC-specific *Ascl1* deletion led to spontaneous autoimmunity.

Mechanistically, ASCL1 shapes mTEC gene expression and influences the AIRE dependency of tissue-specific antigen (TSA) expression in association with changes in chromatin accessibility. Together, these findings establish ASCL1 as a critical regulator of mTEC function and central immune tolerance, acting both independently and cooperatively with AIRE to prevent autoimmunity.

## Results

### Paired single-cell analysis reveals tumor-specific reprogramming of thymic epithelial cells in thymoma patients

We hypothesize that thymoma-associated autoimmunity could arise from primary alterations in tumor TECs that disrupt immune tolerance, followed by secondary changes promoting the expression of self-antigens including muscle antigens, thereby contributing to disease progression. Comparing tumor and normal TECs provides a unique opportunity to identify fundamental mechanisms that safeguard against autoimmunity.

To investigate this, we performed single-cell RNA sequencing (scRNA-seq) of TECs isolated from three spatially separated tumor and three non-tumor regions of whole thymus specimens from thymoma patients (Fig. S1). The three thymoma samples analyzed were classified histologically as WHO type B1, likely B1, and B2, with corresponding Masaoka-Koga stages IIb, III, and III, respectively (Data S1). One patient (type B2) exhibited the MG symptoms (Data S1, Fig. S1). Flow cytometric analysis confirmed the presence of immature CD4⁺CD8⁺ thymocytes in both regions, validating the samples as thymic tissue and indicating that tumor TECs retain T cell-generating capacity (Fig. S1A). scRNA-seq data were integrated using the Harmony package ^34, 35^. After excluding non-TEC clusters (e.g., stromal cells and T cells; Fig. S2), TECs were annotated based on canonical marker gene expression (Fig. S3). Integrative analysis of non-tumorous TECs (normal TECs) with published healthy donor datasets ^7^ ^10^ showed substantial overlap (Fig. S4), suggesting that these cells represent *bona fide* normal TEC populations. Comparison of tumor versus normal TECs revealed subset shifts, including reduced mTEC clusters and a relative increase in cTEC clusters (Fig. 1A, B). Consistent with previous reports ^22^, MG and non-MG TECs were similar in UMAP dimension, and our analysis showed that tumor and normal TECs largely overlapped (Fig. S5A), suggesting that, despite these shifts, tumor TECs appear to retain the capacity to support T cell development.

**Figure 1.**
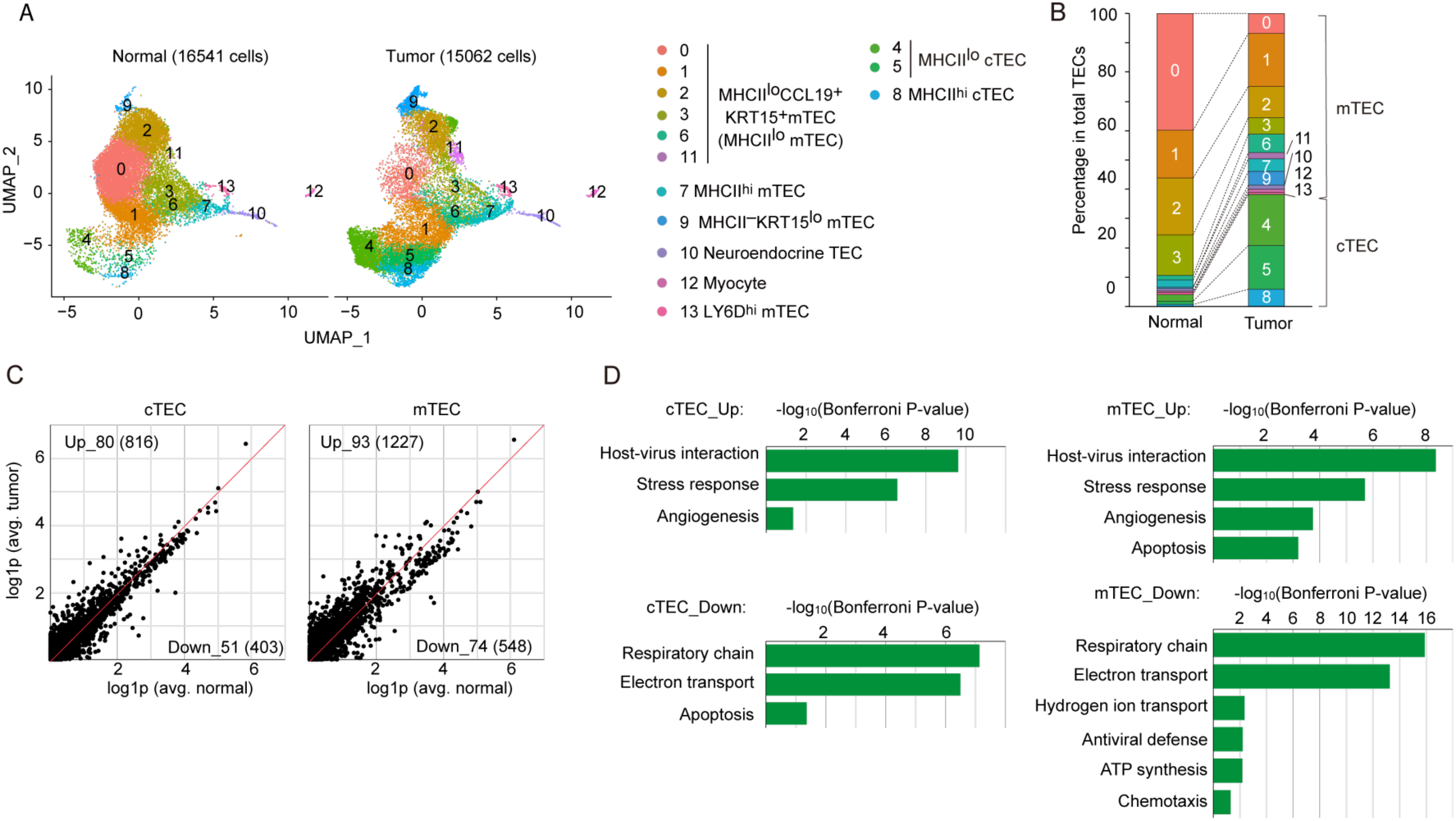
Tumor and normal TECs from thymoma patients exhibit modest differences in gene expression. A. UMAP projection for scRNA-seq data of TECs prepared from 3 non-tumoral (normal) and 3 tumor regions of thymi from 4 human thymoma patients (Data S1). After the data integration, each cluster was assigned to indicated subsets according to expression levels of marker genes (Fig. S3). B. The proportions of individual clusters in total TECs for each sample. Colors and numbers match the left cell population. C. Scatterplot comparing gene expression levels between normal and tumor cTECs and mTECs. Expression values are shown as log1p-transformed counts [logₑ(1 + x)]. The numbers of up-regulated genes (log2FC > 0.25 and P < 0.05) and down-regulated genes (log2FC < -0.25 and P < 0.05) are indicated in the panel (Data S2). D. Gene ontology analysis of up-regulated genes and down-regulated genes in cTECs and mTECs. GO terms of P < 0.05 (Bonferroni) are shown.

We next performed differential gene expression analysis between tumor and normal TECs. Using a permissive threshold (|log₂FC| > 0.25, FDR < 0.05), we identified extensive transcriptional alterations in both mTEC and cTEC populations (Fig. 1C, Data S2). Gene ontology analysis revealed shared upregulation of pathways related to “host–virus interaction”, stress response, and angiogenesis, together with downregulation of genes involved in the “respiratory chain” and “electron transport” in both tumor mTECs and cTECs (Fig. 1D, Data S2I). Tumor mTECs further exhibited reduced expression of genes associated with “mitochondrial ATP production”, consistent with impaired oxidative phosphorylation and a shift toward glycolysis (Fig. 1D, Data S2I). Collectively, these findings indicate that tumor-associated TECs acquire stress-adapted, pro-angiogenic, and metabolically reprogrammed features characteristic of tumor-like phenotypes compared with normal TECs. Subset-specific alterations were also observed: “antiviral defense” and “apoptosis-related genes” were decreased in tumor cTECs, whereas “chemotaxis-related genes” were selectively reduced in tumor mTECs, suggesting impaired immune-supportive functions.

### Downregulation of ASCL1 in human thymoma mTECs and increased autoimmunity susceptibility in mouse model

Given the central role of mTECs in establishing self-tolerance, we next sought key transcriptional regulators underlying the altered gene expression program in tumor mTECs. Analysis of differentially expressed transcription factors (Data S2G) identified ASCL1 as the most significantly and consistently downregulated factor (Fig. 2A, B; Fig. S5B).

**Figure 2.**
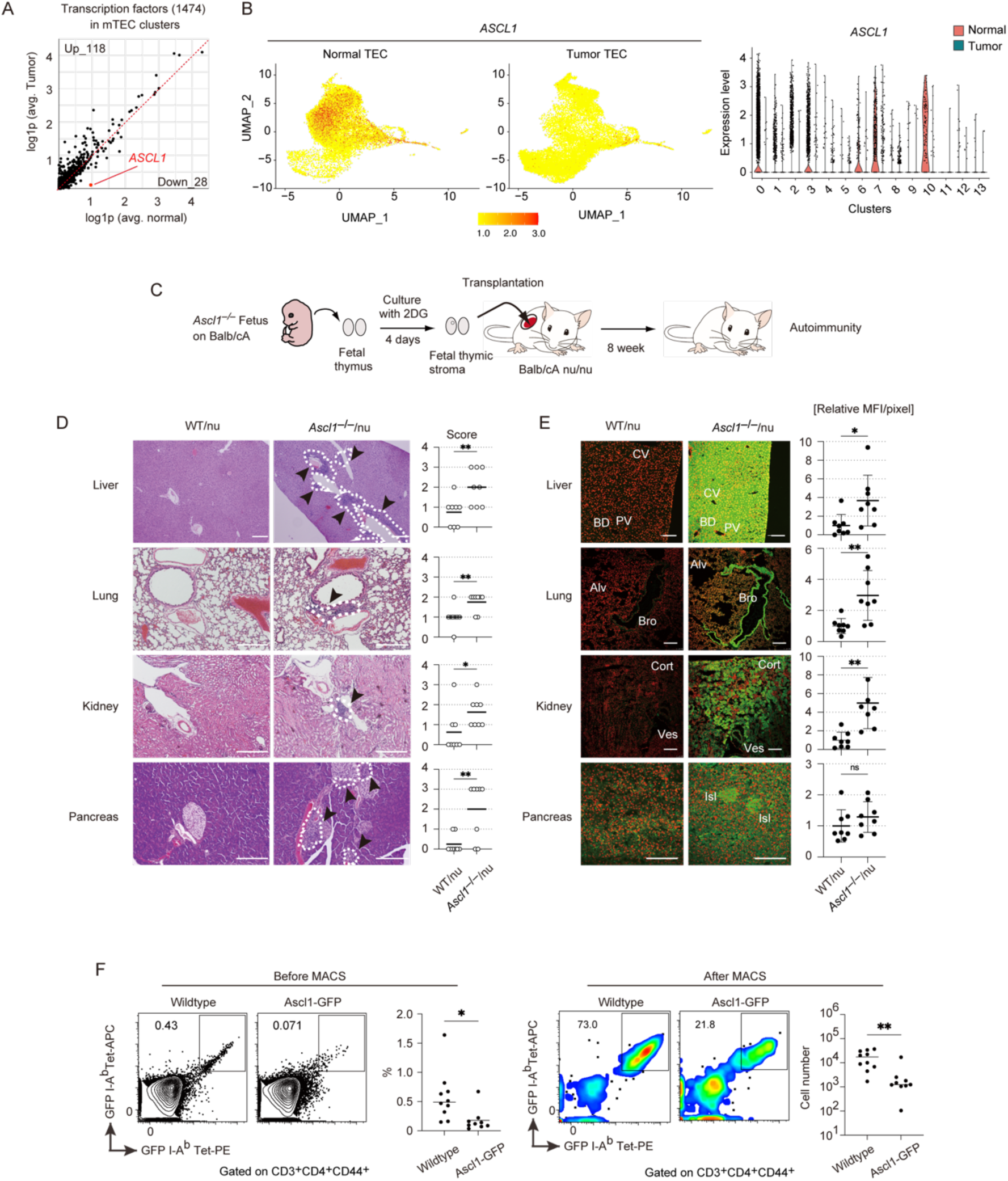
ASCL1 is downregulated in tumor TECs in human thymoma and contributes to thymic stroma–dependent self-tolerance. A. Scatterplot showing transcription factor expression in normal versus tumor human mTECs from thymoma patients. Values are logₑ(1 + x)–transformed. Numbers of upregulated (log₂FC > 0.25, P < 0.05) and downregulated (log₂FC < −0.25, P < 0.05) genes are indicated (Data S2D, E). B. UMAP visualization and violin plot of *ASCL1* expression in normal and tumor human TECs from thymoma patients. C. Experimental scheme of thymic stroma transplantation. D. Hematoxylin and eosin–stained sections of thymus-grafted nude mice (WT/nu and KO/nu), showing inflammatory cell infiltration (arrowheads; N = 8). Scale bars, 200 µm. Scatterplots show infiltration scores; bars indicate means and circles individual animals. P < 0.05, P < 0.01; Wilcoxon rank-sum test. E. Detection of autoantibodies in sera from thymus-grafted nude mice. Organs from *Rag1*⁻^/^⁻ mice were stained with serum (green) and propidium iodide (red). Representative images show staining of hepatocytes (liver), subsets of bronchial epithelial cells (lung), renal tubular cells (kidney), and Langerhans cells (pancreas). Anatomical landmarks: CV, central vein; BD, bile duct; PV, portal vein; Alv, alveoli; Bro, bronchi; Cort, kidney cortex; Ves, kidney vessels; Isl, pancreatic islets. Scale bars, 200 µm. Right panels show relative mean fluorescence intensity (MFI) per pixel (N = 8), normalized to the WT/nu average. Center line, median; box, interquartile range; whiskers, minimum to maximum; points, individual animals. P < 0.05, P < 0.01; two-tailed Student’s t test. F. Flow cytometric analysis of GFP peptide–specific CD4 T cells (CD3⁺CD4⁺CD44⁺Lin⁻) in the spleen. Wild-type and Ascl1-GFP mice were immunized with GFP peptide; enriched GFP-specific CD4 T cells were analyzed. Wild type, N = 10; Ascl1-GFP, N = 9. P < 0.05; two-tailed Student’s t test.

We investigated the role of ASCL1 in thymic self-tolerance using a mouse model. To this end, we transplanted thymic stroma from *Ascl1*-deficient (*Ascl1*^−/−^) mice under the kidney capsule of athymic nude recipients (Fig. 2C). Eight weeks after transplantation, histological analysis revealed lymphocytic infiltration in multiple organs of recipients receiving *Ascl1*^−/−^ thymic stroma (Fig. 2D). Moreover, sera from these recipients contained autoantibodies against various tissues (Fig. 2E and Fig. S6), indicating a breakdown of self-tolerance in the absence of ASCL1 in thymic stroma.

To evaluate the contribution of ASCL1-expressing cells to shaping the T cell repertoire, we adapted a system to quantify T cells specific for GFP, used here as a neo–self-antigen, under polyclonal conditions. *Ascl1*-GFP reporter mice, in which GFP is expressed under the control of the *Ascl1* gene promoter, were immunized with GFP peptides and compared to wild-type controls. Flow cytometric analysis of splenocytes using GFP peptide–MHC tetramers revealed a markedly reduced expansion of GFP-specific T cells in *Ascl1*-GFP mice (Fig. 2F). These results suggest that antigens expressed in ASCL1-expressing cells is recognized as self, supporting a functional role for these cells in establishing T cell tolerance.

Collectively, our findings establish ASCL1 as a critical regulator of thymic stroma-dependent T cell tolerance, with ASCL1-expressing cells contributing to T cell selection. The reduction of ASCL1 in human thymoma mTECs may represent an important mechanism linking thymic dysfunction to autoimmunity.

### ASCL1 regulates mTEC differentiation and cooperates with AIRE to maintain thymic epithelial homeostasis

Previous studies suggest that ASCL1 is expressed in subsets of mTECs, including a fraction of post-Aire mTECs^10, 31, 32, 33^; however, its precise distribution across maturation stages remains unclear. Using Ascl1-GFP reporter mice, flow cytometric analysis revealed that GFP expression was highest in immature mTECs (ITGB4⁺MHCII^lo^) (Fig. 3A, Fig. S7A). In contrast, MHCII^hi^ mature mTECs, including Aire⁺ cells, exhibited lower GFP levels, whereas ITGB4⁻MHCII^lo^ mTECs corresponding to post-Aire mTECs showed minimal GFP, except for a small intermediate-expressing subset. GFP was virtually absent in cTECs. Immunohistochemistry confirmed that ASCL1 protein was predominantly detected in mTECs, particularly within AIRE⁻ TECs (Fig. S7B). Re-analysis of TEC scRNA-seq data ^11^ further demonstrated high *Ascl1* expression in Ccl21⁺ mTEC and TA-TEC clusters, with progressive downregulation along the maturation trajectory (Fig. S7C). *Ascl1* remained detectable in a subset of post-Aire mTECs (Fig. S7C), likely corresponding to endo-TECs ^13, 36^. Together, these findings suggest that *Ascl1* expression is dynamically regulated during mTEC maturation: it is high in immature mTECs and TA-TECs, reduced in mature Aire⁺ mTECs, and further downregulated in post-Aire mTECs, with the exception of a subset consistent with endo-TECs.

**Figure 3.**
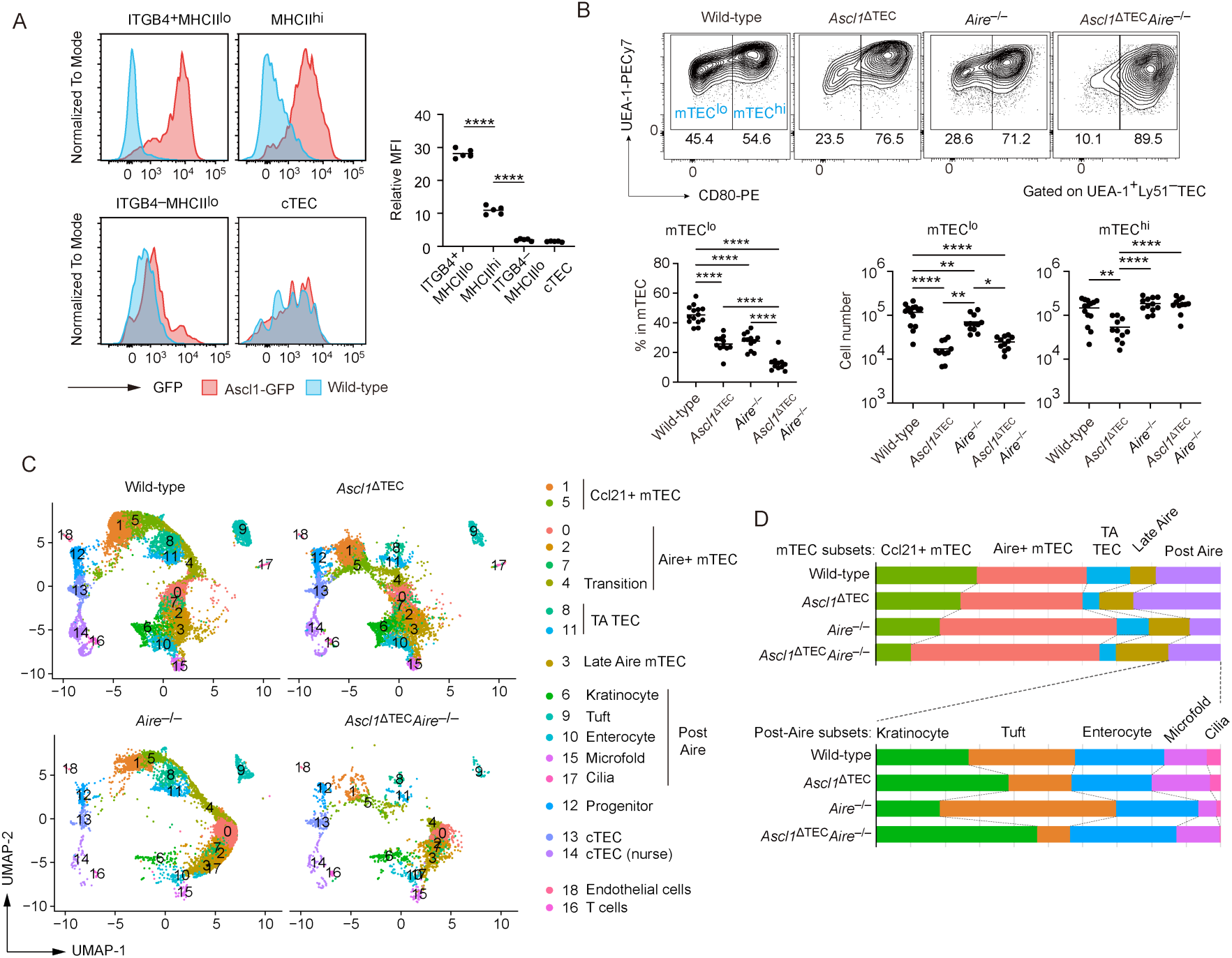
ASCL1 deficiency reshapes the distribution of mTEC subsets. A. Flow cytometric analysis of TECs in 4-week-old Ascl1 GFP mice. Relative MFIs of GFP to control in each TEC subsets are summarized in the right figure (N=5). **** P < 0.0001. Tukey-Kramer test. B. Flow-cytometric analysis of mTECs from 4-week-old control, *Ascl1*^ΔTEC^, *Aire*^−/−^ and *Ascl1*^ΔTEC^*Aire*^−/−^mice. Plots indicate the ratio of mTEC^lo^ in mTECs, cell number of mTEC^lo^, and mTEC^hi^; control (N=14), *Ascl1*^ΔTEC^ (N=11), *Aire*^−/−^ (N=12) and *Ascl1*^ΔTEC^*Aire*^−/−^ (N=11). Data are means (bars) and individual animals (circles). *P<0.05, **P<0.01, ***P<0.001; Tukey-Kramer test. C. UMAP visualization of scRNA-seq data of TECs (EpCAM^+^ CD45^−^ TER119^−^) from 4-week-old wild-type, *Ascl1*^ΔTEC^, *Aire*^−/−^, and *Ascl1*^ΔTEC^*Aire*^−/−^ mice. D. The top graph shows the frequency of each mTEC cluster among total mTECs for each sample in the scRNA-seq dataset. The bottom graph shows the relative frequency of each post-Aire cluster among post-Aire mTECs.

To investigate the role of ASCL1 in mTEC development and function, we generated TEC-specific *Ascl1*-deficient (*Ascl1*^ΔTEC^) mice (Fig. S8A), as germline *Ascl1*^−/−^ mice are neonatally lethal. Because ASCL1 expression precedes AIRE during mTEC maturation, we hypothesized that ASCL1 may prime the transcriptional or chromatin landscape required for subsequent AIRE-dependent gene expression. To determine whether ASCL1 functionally interacts with AIRE, we generated *Ascl1*^ΔTEC^ mice on an Aire-deficient background (*Ascl1*^ΔTEC^*Aire*^−/−^).

Notably, in human TECs, an integrated analysis of published and our own datasets revealed an age-associated decline of *AIRE* expression in both normal thymus and thymoma (Fig. S9).

Accordingly, tumor-associated TECs in thymoma exhibit markedly reduced expression of both ASCL1 and AIRE, suggesting that this compound mutant model may partially recapitulate the molecular state of human tumor-associated TECs.

Body weight, thymic weight, total thymic cellularity, and cortico-medullary structure were unaffected in *Ascl1*^ΔTEC^ and *Ascl1*^ΔTEC^*Aire*^−/−^ mice (Fig. S8B, C, D). Notably, no thymomas were observed by 1 year of age, suggesting that reduced ASCL1 expression in thymoma is more likely a consequence rather than a driver of tumorigenesis.

Thymocyte analysis revealed an increase in CD4 single-positive (CD4SP) M2-stage cells, the final phase of thymocyte maturation^37^, in the thymus of *Ascl1*^ΔTEC^ mice (Fig. S10A, B). In addition, the proportion of Helios⁺ cells within the Wave2B stage of the CD4SP population ^9^ was decreased (Fig. S11), suggesting that ASCL1 contributes to negative selection. By contrast, CD4SP Foxp3⁺ regulatory T cells (Tregs) were increased in *Ascl1*^ΔTEC^ mice (Fig. S10), which might be attributable to an increment of CCR7⁻ Tregs (Fig. S12) that recirculate from the periphery ^38^. These observations suggest that ASCL1 may preferentially contribute to negative selection rather than Treg conversion. Interestingly, these ASCL1-dependent alterations were not observed on the *Aire*-deficient background, indicating that ASCL1 influences thymocyte selection through AIRE-dependent mechanisms.

TEC profiling by flow cytometry demonstrated that that the *Ascl1* deletion led to a reduction in both CD80^low^ mTECs (mTEC^lo^) and CD80^high^ mTECs (mTEC^hi^) populations (Fig. 3B, Fig. S8E, F), whereas cTEC numbers remained unchanged (Fig. S13A). A similar reduction in mTEC^lo^ was observed in thymic grafts reconstituted with *Ascl1*^−/−^ embryonic stroma (Fig. S13B). Based on established classification of mTECs into four subsets (mTEC I-IV) ^6^, *Ascl1* deletion reduced all mTEC subsets, with mTEC I (immature CCL21⁺ mTECs) being most affected, whereas mTEC III (post-Aire) showed relatively modest reduction (Fig. S13C,D). Additionally, AIRE expression was not affected by *Ascl1* deletion (Fig. 13E, F). Importantly, comparison of *Ascl1*^ΔTEC^ and *Ascl1*^ΔTEC^*Aire*^−/−^ mice revealed that the reduction in mTEC^hi^ cells (Fig. 3B) and mTEC II (Fig. S13D) was partially rescued by *Aire* deletion, suggesting that ASCL1 promotes Aire⁺ mTEC development through modulation of AIRE-dependent pathways.

Given the marked heterogeneity of TECs, we performed single-cell RNA-seq on mutant TECs to obtain higher-resolution insight into subset composition. Mutant datasets were integrated with previously reported age-matched wild-type controls ^11^ using canonical correlation analysis implemented in Seurat ^35^ (Fig. S14). Clusters were annotated based on canonical marker expression, and the proportion of each mTEC subset was determined (Fig. 3C). Consistent with prior reports^36, 39^, *Aire* deficiency increased the proportion of Aire⁺ mTECs while reducing post-Aire subsets. In contrast, ASCL1-deficient mice (*Ascl1*^ΔTEC^ and *Ascl1*^ΔTEC^*Aire*^−/−^) exhibited a distinct pattern. Ccl21⁺ mTECs were reduced in both mutants, consistent with flow cytometric analysis, and TA-TECs were also decreased. In the post-Aire compartment, tuft-like mTECs were reduced, whereas keratinocyte subsets might display an modest increase (Fig. 3D). In addition, *Chgb⁺* endo-TEC–like cells were decreased (Fig. S14C), consistent with a previous report ^13^. Overall, *Ascl1* deficiency preferentially impairs Ccl21⁺ mTECs and TA-TECs and selectively alters post-Aire lineages, supporting a role for ASCL1 in early mTEC differentiation and subsequent lineage specification.

### Stage-specific genetic interplay between ASCL1 and AIRE regulates mTEC transcriptional programs

In the UMAP projection of integrated scRNA-seq data, *Ascl1* deletion induced a pronounced shift in Ccl21⁺ mTEC clusters, indicating broad transcriptional reprogramming (Fig. 4A), whereas other TEC subsets, including post-Aire mTECs and cTECs, were comparatively stable across genotypes (Fig. S15F). In contrast, *Aire* deletion exerted only modest effects on Ccl21⁺ mTECs, and the combined deletion of *Aire* and *Ascl1* largely phenocopied *Ascl1* deletion alone, establishing ASCL1 as the dominant regulator of Ccl21⁺ mTEC identity. The pattern was reversed in Aire⁺ mTECs: Aire deficiency markedly altered their distribution, whereas *Ascl1* loss alone had minimal impact (Fig. 4A). Notably, *Aire^−/−^* and *Ascl1*^ΔTEC^*Aire^−/−^* mice displayed substantially divergent Aire⁺ mTEC distribution. Consistently, differential expression analysis demonstrated largely distinct sets of downregulated genes in *Aire^−/−^* and *Ascl1*^ΔTEC^*Aire^−/−^* mTECs (Fig. S15A,B). Together, these findings indicate that ASCL1 is the primary regulator of Ccl21⁺ mTECs, whereas AIRE predominates in Aire⁺ mTECs. Of note, the impact of *Ascl1* deletion on Aire⁺ mTECs becomes evident only in the absence of AIRE, implying a genetic interaction between *Ascl1* and *Aire*.

**Figure 4.**
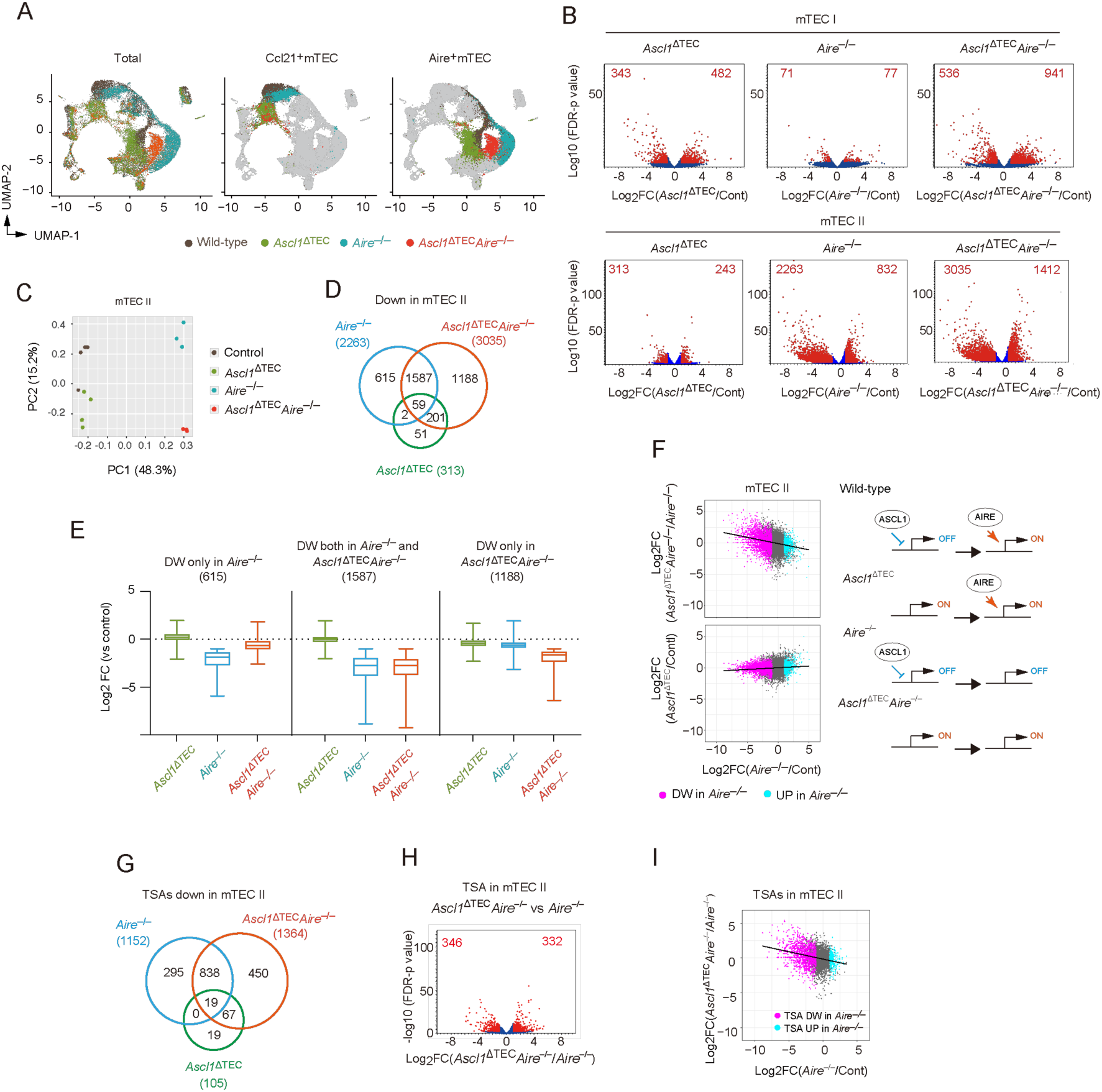
ASCL1 modulates mTEC phenotypes and AIRE function. A. UMAP visualization of scRNA-seq data from total TECs, Ccl21⁺ mTECs, and Aire⁺ mTECs isolated from 4-week-old control, *Ascl1*^ΔTEC^, *Aire*^−/−^, and *Ascl1*^ΔTEC^*Aire*^−/−^ mice. B. Volcano plots showing differential gene expression from bulk RNA-seq of mTEC I (top) and mTEC II (bottom) in *Ascl1*^ΔTEC^, *Aire*^−/−^, and *Ascl1*^ΔTEC^*Aire*^−/−^ mice compared with controls. Red dots indicate genes with FDR < 0.05 and fold change > 2 or < −2. Sample sizes: mTEC I, N = 3 per group; mTEC II, N = 4 (control, *Ascl1*^ΔTEC^) and N = 3 (*Aire*^−/−^, *Ascl1*^ΔTEC^*Aire*^−/−^). C. Principal component analysis (PCA) of bulk RNA-seq data from mTEC II populations (control, N = 4; *Ascl1*^ΔTEC^, N = 4; *Aire*^−/−^, N = 3; *Ascl1*^ΔTEC^*Aire*^−/−^, N = 3). D. Venn diagram showing overlap of genes significantly downregulated in mTEC II (FDR < 0.05, fold change < −2). E. Box plots showing log2-transformed fold changes of normalized expression values in *Ascl1*^ΔTEC^, *Aire*^−/−^, and *Ascl1*^ΔTEC^*Aire*^−/−^ mice compared with controls in mTEC II. Genes of “DW only in *Aire*^−/−^ (615)” in the left, “DW both in *Aire*^−/−^ and *Ascl1*^ΔTEC^*Aire*^−/−^ (1587)” in the center, and “DW only in *Ascl1*^ΔTEC^*Aire*^−/−^ (1188)” in the right were selected. F. Scatter plots comparing log₂ fold changes in mTEC II between *Ascl1*^ΔTEC^*Aire*^−/−^ versus *Aire*^−/−^and *Aire*^−/−^ versus control (left), and *Aire*^−/−^ versus control and *Ascl1*^ΔTEC^ versus control (right). Pink dots indicate genes downregulated in *Aire*^−/−^ mTECs; light blue dots indicate upregulated genes in *Aire*^−/−^ mTECs. Schematics summarize proposed regulatory mechanisms. G. Venn diagram of tissue-specific antigen (TSA) genes downregulated in mTEC II of *Ascl1*^ΔTEC^ (N = 4), *Aire*^−/−^ (N = 3), and *Ascl1*^ΔTEC^*Aire*^−/−^ (N = 3) mice relative to controls (N = 4). H. Volcano plot showing differential TSA expression in mTEC II of *Ascl1*^ΔTEC^*Aire*^−/−^ versus *Aire*^−/−^ mice. I. Scatter plot comparing log₂ fold changes in TSA expression between *Ascl1*^ΔTEC^*Aire*^−/−^ and *Aire*^−/−^ mTEC II populations.

To mitigate the sparsity of scRNA-seq, we performed bulk RNA-seq on sorted mTEC I and II subsets, which correspond to predominantly Ccl21⁺ mTECs and Aire⁺ mTECs, respectively. As in scRNA-seq, the *Ascl1* loss induced widespread changes in mTEC I, while the *Aire* loss had a less effect (Fig. 4B and Data S3A, B). Conversely, transcriptional alterations in mTEC II were more pronounced following *Aire* deletion than *Ascl1* loss (Fig. 4B and Data S3A, B). Notably, whereas *Aire* deletion primarily resulted in gene downregulation, *Ascl1* deletion led to both up-and downregulation across both subsets, suggesting a dual regulatory role for ASCL1 in mTEC gene expression (Fig. 4B).

Principal component analysis (Fig. 4C) and volcano plot comparison (Fig. S15C) further underscored the distinct mTEC II transcriptional profiles between *Aire^−/−^* and *Ascl1*^ΔTEC^*Aire^−/−^*mice. In downregulated genes, while 1,587 were shared between the two mutants, 1,188 were uniquely downregulated in mTEC II from *Ascl1*^ΔTEC^*Aire^−/−^* mice (Fig. 4D, E, Data S3E), indicating functional redundancy and cooperativity between ASCL1 and AIRE.

Conversely, 615 genes were selectively downregulated in *Aire^−/−^* mTEC II (Fig. 4D, E, Data S3E), suggesting antagonistic effect of ASCL1 against AIRE-dependent gene program. Supporting this, genes downregulated in *Aire^−/−^* mTEC II were preferentially upregulated when *Ascl1* was additionally deleted in the *Aire*-deficient background (Fig. 4F), a reverse correlation not observed in the *Aire*-sufficient context. These findings suggest that AIRE tends to activate genes that are pre-suppressed by ASCL1 (Fig. 4F, right).

Given the essential role of TSAs in tolerance, it is notable that TSAs comprised approximately 50% of downregulated genes in mTEC II from both *Aire^−/−^*and *Ascl1*^ΔTEC^*Aire^−/−^* mice (Fig. 4G, Fig. S15D, Data S4), although the affected TSA repertoires differed substantially (Fig. 4G, H and Fig. S15E). Importantly, ASCL1 antagonized AIRE-driven TSA expression in a pre-emptive manner (Fig. 4I), highlighting its unique role also in shaping the TSA repertoire.

Collectively, these results demonstrate that ASCL1 exerts a dual role in AIRE-driven gene regulation, including modulation of TSA expression, through both cooperative and antagonistic interactions.

### ASCL1 shapes chromatin accessibility programs upstream of AIRE in mTECs

ASCL1 has been reported to regulate chromatin accessibility in several cell types in addition to controlling transcription ^40^; however, its role in mTECs remains undefined. To determine whether ASCL1 contributes to chromatin accessibility in mTECs, we performed single-cell ATAC-seq (scATAC-seq) on TECs isolated from mutant and control mice. Following data integration, UMAP embedding, clustering, and annotation were conducted using established analytical pipelines ^11, 41^ (Fig. S16). DNA motif analysis revealed that, in control mice, genomic regions containing ASCL1 motifs were preferentially accessible in Ccl21⁺ mTECs compared with other TEC populations (Fig. S16D, E).

Deletion of Ascl1 altered the distribution of Ccl21⁺ mTEC and TA-TEC clusters in UMAP space (Fig. 5A , Fig. S16D and Fig. S17A) and reduced ASCL1 motif activity in these populations (Fig. S16E). In contrast, *Aire* deletion primarily affected Aire⁺ mTEC clusters relative to other TEC subsets (Fig. 5A and Fig. S17A). Notably, the distribution of Aire⁺ mTEC clusters differed between *Aire^−/−^* and *Ascl1*^ΔTEC^*Aire^−/−^*mice (Fig. 5A), indicating a functional interaction between ASCL1 and AIRE in regulating chromatin accessibility, consistent with our RNA-seq data. By comparison, post-Aire mTECs, tuft-like mTECs, and cTECs exhibited largely similar distributions across genotypes (Fig. S17A) , suggesting more limited effects in these populations.

**Figure 5.**
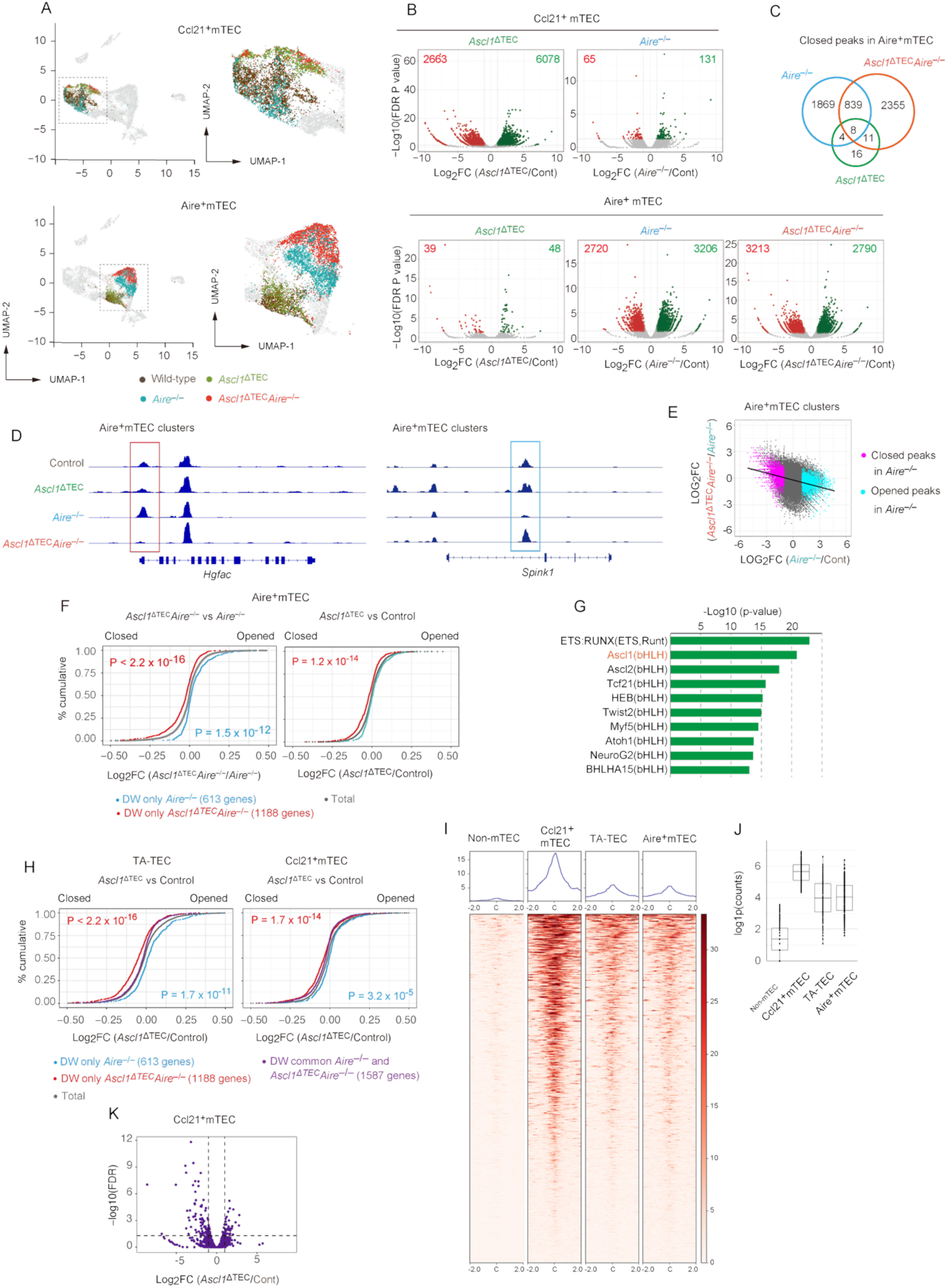
ASCL1 modifies the chromatin landscape to define gene expression programs in mTECs. A. UMAP visualization of Ccl21⁺ and Aire⁺ mTECs from control, *Ascl1*^ΔTEC^, *Aire*^−/−^, and *Ascl1*^ΔTEC^*Aire*^−/−^ mice. Enlarged plots were in the right. B. Volcano plots of differentially open (green) and closed (red) chromatin regions in Ccl21⁺ and Aire⁺ mTECs across genotypes compared with controls. C Venn diagram of ATAC-seq peaks significantly closed (FDR P < 0.05, >2-fold) in Aire⁺ mTECs of each mutant genotype relative to controls. D. Representative genomic loci showing regions selectively closed in Aire⁺ mTECs of *Ascl1*^ΔTEC^*Aire*^−/−^ (left) or *Aire*^−/−^ (right) mice compared with other genotypes. E. Scatter plot comparing log₂ fold changes of ATAC peaks in Aire⁺ mTECs between *Aire*^−/−^versus control and *Ascl1*^ΔTEC^*Aire*^−/−^ versus control mice. Pink and blue dots indicate significantly closed regions. F. Cumulative distribution function (CDF) showing changes in ATAC-derived gene activity in Aire⁺ mTECs, relating scATAC-seq to bulk RNA-seq data. Left, *Ascl1*^ΔTEC^*Aire*^−/−^ versus *Aire*^−/−^; right, *Ascl1*^ΔTE^ versus control. Blue and red dots denote genes uniquely downregulated in *Aire*^−/−^or *Ascl1*^ΔTEC^*Aire*^−/−^ mTEC II, respectively; gray indicates all genes (Mann–Whitney U test). G. Top 10 enriched transcription factor motifs in distal regulatory regions associated with genes specifically downregulated in *Ascl1*^ΔTEC^*Aire*^−/−^ mTEC II. H. CDF of ATAC-derived gene activity changes in TA-TECs and Ccl21⁺ mTECs from *Ascl1*^ΔTEC^ versus control mice, highlighting *Aire*^−/−^-specific (blue), *Ascl1*^ΔTEC^*Aire*^−/−^-specific (red), and shared (purple) downregulated mTEC II gene sets (Mann–Whitney U test). I. Heatmap and metaprofile of ATAC-seq signal centered on ASCL1 motifs (±2 kb) identified from the JASPAR database within distal regulatory regions associated with the “DW only Ascl1ΔTEC Aire–/–” gene set. J. Quantification of distal regulatory ATAC peaks containing ASCL1 motifs linked to genes specifically downregulated in *Ascl1*^ΔTEC^*Aire*^−/−^ mTEC II (“DW only *Ascl1*^ΔTEC^*Aire^−/−^*”). K. Volcano plot showing differential abundance of ASCL1 motif–containing distal regulatory regions in Ccl21⁺ mTECs from *Ascl1*^ΔTEC^ versus control mice (FDR < 0.05, >2-fold).

Differential accessibility analysis revealed widespread chromatin changes in Ccl21⁺ mTECs and TA-TECs of *Ascl1*^ΔTEC^ mice, and in Aire⁺ mTECs of *Aire^−/−^* mice, relative to controls (Fig. 5B and Fig. S17B), indicating that ASCL1 and AIRE influence distinct mTEC subsets. In Aire⁺ mTECs from *Ascl1*^ΔTEC^*Aire^−/−^*mice, chromatin accessibility was also extensively altered; however, many affected regions were distinct from those altered in *Aire^−/−^* mice (Fig. 5C and Fig. S17C, D). Notably, 2,355 peaks were uniquely altered in the double mutant but not in either single mutant (Fig. 5C, D), indicating that ASCL1 and AIRE redundantly maintain chromatin accessibility at a subset of loci in Aire⁺ mTECs. Conversely, 1,869 regions were altered exclusively in *Aire^−/−^* mice but not in the double mutant (Fig. 5C). Moreover, regions that became less accessible in *Aire^−/−^* mTECs showed increased accessibility when *Ascl1* was deleted in the AIRE-deficient background (Fig. 5E). This pattern suggests that AIRE-dependent chromatin opening at these loci requires prior antagonistic activity of ASCL1. Together, these findings indicate that ASCL1 exerts a dual influence on chromatin accessibility in Aire⁺ mTECs: a redundant role with AIRE at certain loci, and a pre-antagonistic function that licenses AIRE-dependent chromatin opening at others. A similar pattern was observed at TSA-associated loci (Fig. S18), supporting a role for this regulatory interplay in the establishment of central tolerance.

Cumulative distribution analysis of log₂ fold changes in chromatin accessibility (scATAC-seq; gene body and promoter regions) and gene expression (bulk RNA-seq; Fig. 4) demonstrated coordinated alterations in Ccl21⁺ and Aire⁺ mTECs following deletion of *Ascl1*, *Aire*, or both (Fig. S19A, B), supporting a link between chromatin accessibility and transcriptional output. Among the 615 genes uniquely downregulated in mTEC II from *Aire^−/−^* mice (“DW only *Aire^−/−^*”), chromatin accessibility in Aire⁺ mTECs increased upon Ascl1 deletion in the AIRE-deficient background (*Ascl1*^ΔTEC^*Aire^−/−^*vs *Aire^−/−^*; Fig. 5F left, blue; Fig. S19C). This increase was not observed under AIRE-sufficient conditions (*Ascl1*^ΔTEC^ vs control; Fig. 5F right, blue; Fig. S19C). In contrast, for the 1,188 genes uniquely downregulated in mTEC II from *Ascl1*^ΔTEC^*Aire^−/−^* mice (“DW only *Ascl1*^ΔTEC^*Aire^−/−^*”), chromatin accessibility in Aire⁺ mTECs was reduced irrespective of AIRE expression (Fig. 5F, red; Fig. S19C). ASCL1 motifs were enriched in distal regulatory regions associated with these genes (Fig. 5G). Collectively, these data support a model in which ASCL1-dependent chromatin remodeling contributes directly to gene expression programs in Aire⁺ mTECs and functions upstream of AIRE in the transcriptional hierarchy.

*Ascl1* is highly expressed in Ccl21⁺ mTECs and TA-TECs (Fig. 3A and Fig. S7C), which serve as precursors of Aire⁺ mTECs ^11, 42^. Cumulative distribution analysis demonstrated that genomic regions linked to the “DW only *Ascl1*^ΔTEC^*Aire^−/−^*” gene set were already less accessible in TA-TECs and Ccl21⁺ mTECs following *Ascl1* deletion (Fig. 5H, Fig. S19D). Motif-centered heatmap analysis and quantification of motif-containing ATAC peaks further revealed that ASCL1 motif–containing distal peaks associated with the “DW only *Ascl1*^ΔTEC^*Aire^−/−^*” gene set were preferentially accessible in Ccl21⁺ mTECs compared with non-mTEC populations, including cTECs and progenitors, and remained accessible in TA-TECs and Aire⁺ mTECs, although accessibility was reduced relative to Ccl21⁺ mTECs. (Fig. 5I, J), consistent with chromatin accessibility established in immature mTECs being maintained during differentiation. Accessibility at these sites in Ccl21⁺ mTECs also tended to be reduced in the absence of ASCL1 (Fig. 5K). Together, these findings support a model in which ASCL1 establishes chromatin accessibility patterns in immature Ccl21+ mTECs that persist into the Aire⁺ mTEC II stage.

In contrast, genomic regions associated with the “DW only *Aire^−/−^*” gene set displayed increased accessibility in precursor populations upon *Ascl1* deletion (Fig. 5H, blue; Fig. S19D), consistent with ASCL1-dependent repression of these regions, although these effects are likely indirect. Finally, chromatin accessibility at loci corresponding to genes commonly downregulated in both *Ascl1*^ΔTEC^*Aire^−/−^*and *Aire^−/−^* mTECs remained relatively stable in TA-TECs and Ccl21⁺ mTECs (Fig. 5H, purple; Fig. S19D), indicating independence from ASCL1-dependent chromatin remodeling. Collectively, these results demonstrate that ASCL1 contributes to chromatin accessibility at a subset of AIRE-regulated genes in precursor mTECs, thereby shaping the transcriptional programs of Aire⁺ mTECs (Fig. S20).

### ASCL1 potentiates AIRE-dependent transcriptional programs

If ASCL1 primes the AIRE-dependent program, it may also modulate AIRE-driven gene expression outside the thymic epithelial context. To test this, we used HEK293 cells, which lack endogenous ASCL1 and AIRE and therefore provide a tractable system for controlled co-expression ^43, 44, 45^. Cells were transfected with ASCL1, AIRE, or both, sorted based on IRES-mediated GFP and RFP expression (Fig. S21A, B), and subjected to RNA-seq.

ASCL1 overexpression significantly upregulated 9 genes (Fig. 6A and Fig. S21C), including *ATOH8* and *ID1*, the murine orthologs of which were downregulated in mTEC I of *Ascl1*^ΔTEC^ mice (Data S3C, S5B). Consistent with previously reported ^43, 44, 45^, AIRE overexpression modestly induced genes such as *KRT14*, *S100A8*, *KRT5*, and *CHRNA1*, though not reaching statistical significance (Fig. 6B). Notably, co-expression of ASCL1 and AIRE led to further upregulation of these genes, suggesting a synergistic effect. In addition, TSAs including *AGXT*, *PRSS1*, and *PLA2G4D* were significantly induced upon co-expression of ASCL1 and AIRE (Fig. 6B).

**Figure 6.**
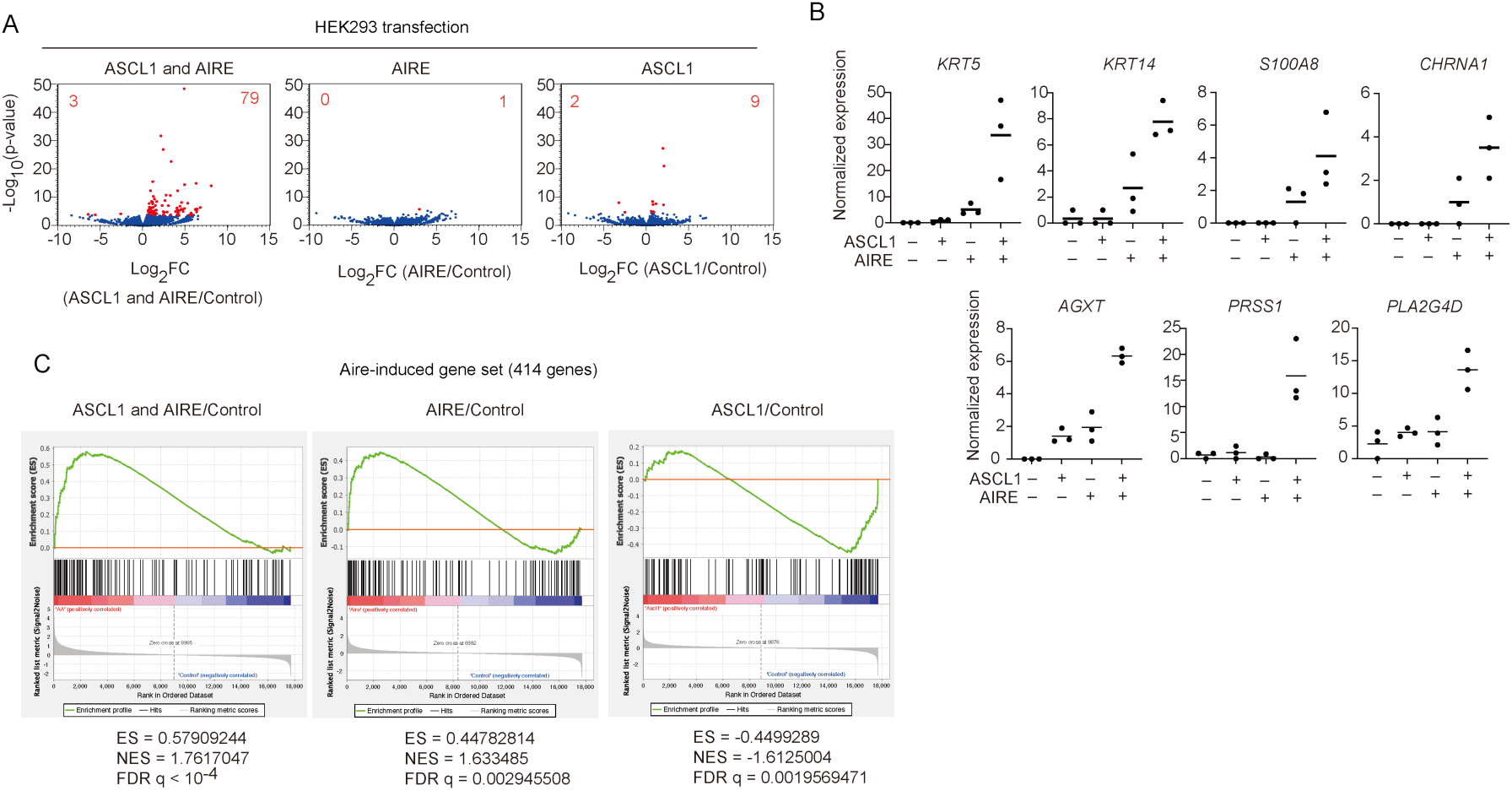
ASCL1 promotes AIRE-dependent gene expression in non-TECs. A. Volcano plots showing differential gene expression in HEK293 cells transfected with ASCL1, AIRE, or both, compared to control. Red dots indicate significantly differentially expressed genes (FDR P < 0.05). The number of differentially expressed genes is indicated in each panel. B. Normalized expression levels of representative genes in HEK293 cells transfected with ASCL1, AIRE, or both (N=3 each). C. Gene set enrichment analysis (GSEA) of human orthologs of mouse Aire-induced genes (n = 414) in ASCL1+AIRE-transfected cells vs. control, AIRE-transfected cells vs. control and, ASCL1-transfected cells vs. control,.

Among the 79 genes significantly upregulated by co-expression of ASCL1 and AIRE (Data S5), 34 (43.0%) were classified as TSA genes, and 24 (30.4%) corresponded to human orthologs of previously defined Aire-dependent genes ^5^ (Fig. S21D). Gene set enrichment analysis (GSEA) using a curated Aire-induced gene set, defined as transcripts absent in Aire-deficient Aire-positive mTECs ^5^ (Data S5D), confirmed enrichment following AIRE transfection (Fig. 6C). In contrast, the Aire-induced gene set was enriched among genes downregulated by ASCL1 alone (Fig. 6C), suggesting a repressive effect of ASCL1 in the absence of AIRE. Notably, co-expression of ASCL1 with AIRE further increased the enrichment of Aire-induced genes among upregulated transcripts compared with AIRE alone (Fig. 6C). Together, these findings support a model in which ASCL1 primes gene loci for subsequent activation by AIRE.

### ASCL1 and AIRE synergistically restrain peripheral autoimmunity

To investigate the cooperative impact of *Aire* and *Ascl1* deletions on autoimmune phenotypes, we performed flow cytometric analysis of peripheral T cells. Effector memory T cells (CD44⁺CD62L⁻) in the lymph nodes were markedly increased in *Ascl1*^ΔTEC^*Aire^−/−^*mice, with milder increases observed in *Ascl1*^ΔTEC^ or *Aire^−/−^*mice individually (Fig. 7A and Fig. S22), suggesting a synergistic role for AIRE and ASCL1 in restraining peripheral T cell self-activation. Consistent with findings from thymic stroma transplantation studies, *Ascl1*^ΔTEC^ mice exhibited immune cell infiltration in several organs (Fig. 7B). Some organs, such as eyes, salivary glands, and stomach, were affected only in *Aire^−/−^* mice but not in *Ascl1*^ΔTEC^ mice, indicating distinct autoimmune manifestations between the single mutants (Fig. 7B and Fig. S23). Consistent with the flow cytometric data, *Ascl1*^ΔTEC^*Aire^−/−^* mice exhibited a broader spectrum of affected tissues compared with either single mutant, suggesting more severe autoimmune phenotypes due to the combined or synergistic effect of both Ascl1 and Aire deletions (Fig. 7B and Fig. S23).

**Figure 7.**
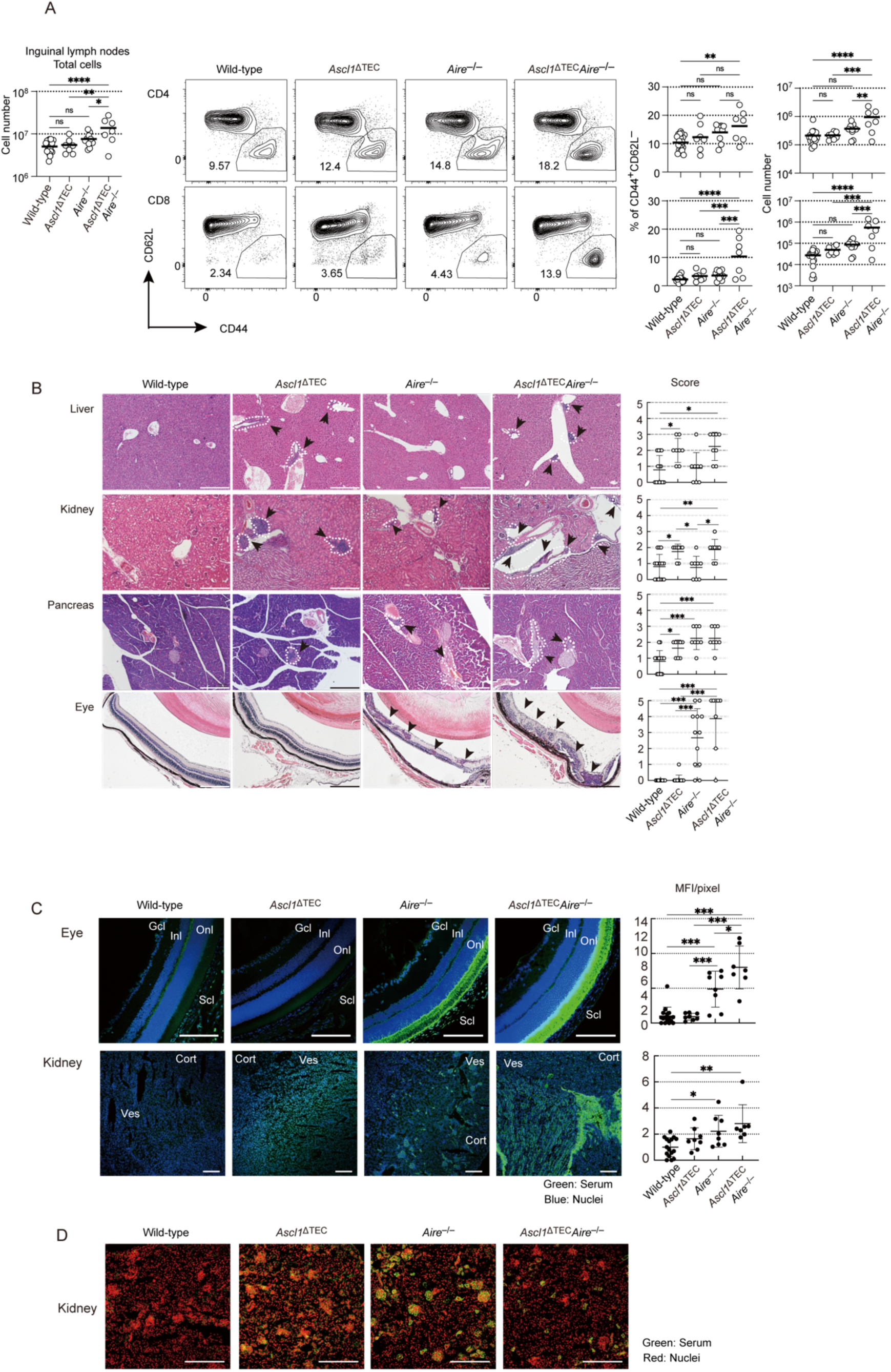
ASCL1 and AIRE cooperatively regulates autoimmunity. A. The panel on the left shows total cell number of inguinal lymph nodes from 20-week-old control, *Ascl1*^ΔTEC^, *Aire*^−/−^, and *Ascl1*^ΔTEC^*Aire*^−/−^ mice. Center panels show representative data of flow cytometric analysis of inguinal lymph nodes from 20-week-old control, *Ascl1*^ΔTEC^, *Aire*^−/−^, and *Ascl1*^ΔTEC^*Aire*^−/−^ mice in the center. Distributions of CD4- and CD8-positive cells in lymphocytes are shown on the left. Control (n = 19), *Ascl1*^ΔTEC^ (N=7), *Aire*^−/−^ (N=10), and *Ascl1*^ΔTEC^*Aire*^−/−^ (N=7). Percentages and cell numbers of CD44^hi^CD62L^lo^ cells in CD4- and CD8-positive T cells in inguinal lymph nodes are shown on the right. Data are means (bars) and individual animals (circles). *P<0.05, **P<0.01, ***P<0.001; Tukey-Kramer test. B. Histological analysis of some organs from control (N=15), *Ascl1*^ΔTEC^ (N=8), *Aire*^−/−^ (N=8, N=12 for eye), and *Ascl1*^ΔTEC^*Aire*^−/−^ (N=8) mice at 20 weeks, H&E staining for infiltrating lymphocytes. White dotted lines and arrows: inflammatory cell infiltration. Left: representative sample data. Bars, 200 µm. Right: infiltration scores. *P<0.05, ***P<0.001; Steel-Dwass test. C. Detection of autoantibodies. Organs from *Rag1^−/−^* mice were stained with sera (green) from control (n = 16), *Ascl1*^ΔTEC^ (N=8), *Aire*^−/−^ (N=8), and *Ascl1*^ΔTEC^*Aire*^−/−^ (N=7) mice at 20 weeks and DAPI (blue); Scale bars, 200 µm. Eye, Gcl: ganglion cell layer, Inl: inner nuclear layer, Onl: outer nuclear layer, Ch: choroid, Scl: sclera. Kidney, Cort: cortex, Ves: vessels. Right plots, relative MFI/pixel of autoantibodies. *P<0.05, **P<0.01, ***P<0.001; Tukey-Kramer test. D. Typical panels of detection of autoantibodies against kidney cortex. Kidney from *Rag1^−/−^*mice were stained with sera (green) from control, *Ascl1*^ΔTEC^, *Aire*^−/−^, and *Ascl1*^ΔTEC^*Aire*^−/−^ mice at 20 weeks and proprium iodide (red); Scale bars, 200 µm.

Immunofluorescence analysis to detect serum autoantibodies revealed that the combined deletion of *Aire* and *Ascl1* resulted in elevated antibody titers against multiple tissues, including the eye, salivary glands, and stomach, compared with single-mutant mice (Fig. 7C and Fig. S23). In the kidney, antigenic targets varied by genotype: while *Aire^−/−^* mice produced autoantibodies against glomerulus and perivascular cells (Fig. 7D), sera from *Ascl1*^ΔTEC^*Aire^−/−^*mice did not react glomerulus and instead showed strong reactivity toward renal medullary cells (Fig. 7C, D). These differences may reflect distinct alterations in the TSA repertoire presented by mTECs in *Aire^−/−^* versus *Ascl1*^ΔTEC^*Aire^−/−^*mice.

Because reduced TSA expression could theoretically predispose to organ-specific autoimmunity, we next examined whether loss of ASCL1 and AIRE promotes features of myasthenia gravis (MG), an autoimmune disease strongly associated with thymic abnormalities and characterized by anti–acetylcholine receptor (AChR) antibodies. Our scRNA-seq data indicated reduced expression of CHRNA1, which encodes AchR, in tumor mTECs compared with normal mTECs (Fig. S24A). Similarly, expression levels of mouse ortholog *Chrna1* and a subset of other potential MG antigen genes ^29^ were reduced in *Aire*^−/−^ and *Ascl1*^ΔTEC^*Aire^−/−^*mice (Fig. S24B).

However, we did not observe any MG-like autoimmune symptoms and anti-AchR antibody generation in these mice. These findings suggest that the disease onset of MG requires additional events beyond the reduction of ASCL1 and AIRE.

### TEC abnormalities in *Ascl1*^ΔTEC^*Aire^−/−^* mice partially recapitulate transcriptional features of human thymoma

Given that ASCL1 downregulation likely reflects a downstream consequence of thymoma development, along with the age-associated decline in AIRE expression, we examined whether TEC abnormalities in *Ascl1*^ΔTEC^*Aire^−/−^* mice recapitulate features of tumor TECs observed in human thymoma, while acknowledging that this model may not capture all aspects of the disease.

Differential gene expression analysis revealed partial overlap: approximately 29% (21/73) of downregulated and 23% (21/92) of upregulated genes in human thymoma mTECs (P < 0.05, ≥2-fold change) corresponded to the human orthologs of genes dysregulated in mTECs of *Ascl1*^ΔTEC^*Aire^−/−^*mice (Data S3C, D). Among the commonly upregulated genes, we noticed on the aberrant induction of cTEC-associated genes such as *PRSS16* (Fig. 8A) and *CCL25* (Fig. S25A) in tumor mTECs. GSEA revealed a significant enrichment of a human cTEC-preferential gene signature (Data S6) among the upregulated genes in tumor mTECs (Fig. 8B), suggesting that tumorigenesis induces aberrant cTEC-like transcriptional features in mTECs. In mice, *Prss16* and *Ccl25* were upregulated in mTECs from both *Ascl1*^ΔTEC^ and *Ascl1*^ΔTEC^*Aire^−/−^*mice (Fig. 8C and Fig. S25B), indicating that their dysregulation is driven primarily by ASCL1 deficiency. GSEA confirmed the enrichment of a murine cTEC-preferential gene set (Data S7) in both mTEC I and mTEC II populations of *Ascl1*^ΔTEC^ mice (Fig. 8D), mirroring the transcriptional shifts observed in tumor mTEC of human thymoma. These findings establish ASCL1 as a conserved repressor of cTEC-like gene expression in mTECs, preserving the fidelity of the mTEC transcriptional program.

**Figure 8.**
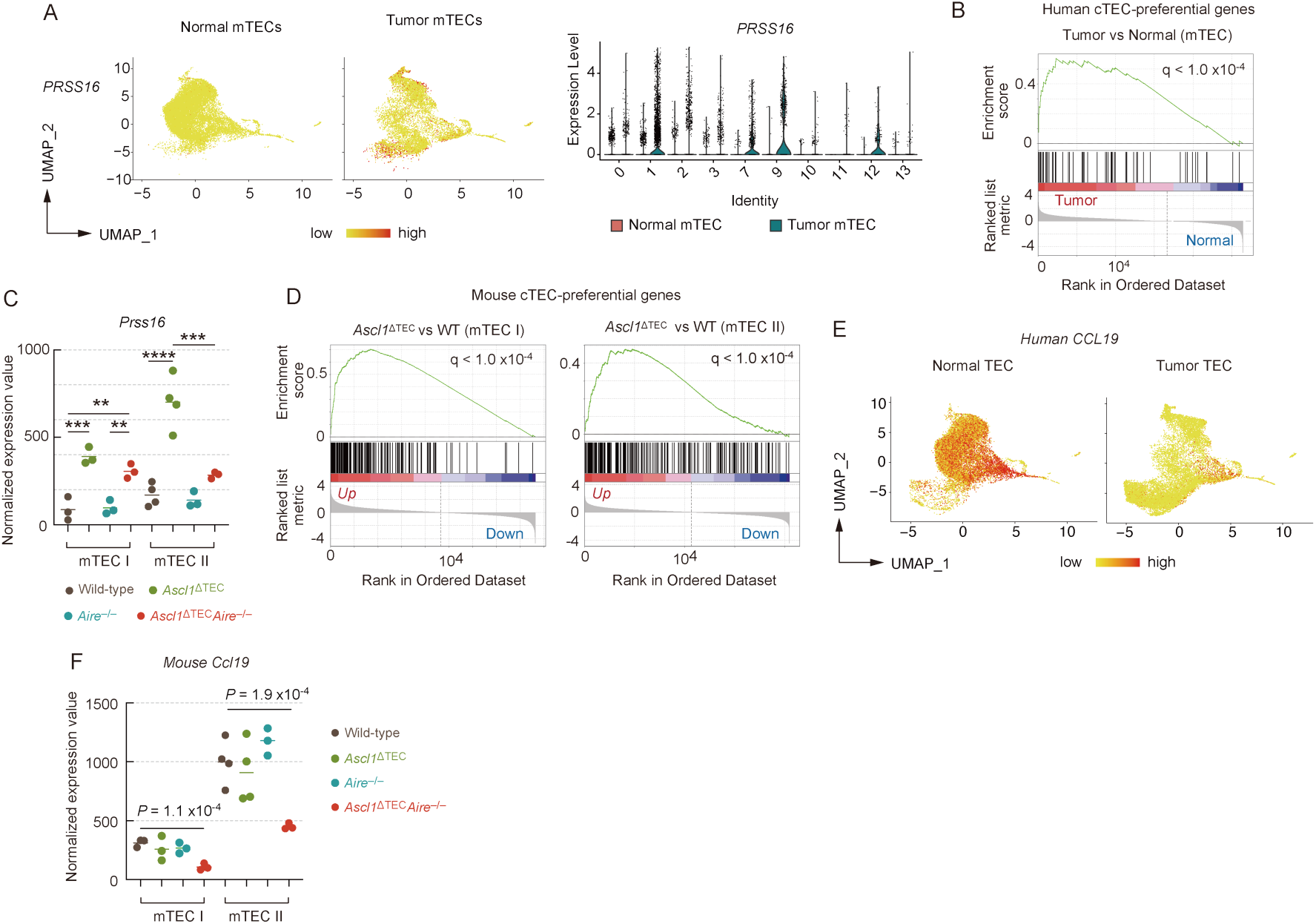
Dual deletion of ASCL1 and AIRE in TECs reproduces certain thymoma-associated characteristics. A. UMAP showing PRSS16⁺ mTECs in normal and tumor samples (left). Violin plots compare PRSS16 expression between normal and tumor mTEC clusters (right). B. GSEA comparing tumor vs. normal mTECs using human cTEC-preferential genes (defined from normal TEC scRNA-seq: log₂FC > 2, P < 0.05; Data S6). C. Normalized expression values of mouse *Prss16* in mTEC I and mTEC II of control, *Ascl1*^ΔTEC^, *Aire*^−/−^, and *Ascl1*^ΔTEC^*Aire*^−/−^ mice in bulk RNA-seq analysis. D. GSEA of mouse cTEC-preferential genes in mTEC I and mTEC II from control, *Ascl1*^ΔTEC^, *Aire*^−/−^, and *Ascl1*^ΔTEC^ *Aire*^−/−^mice. cTEC-preferential genes (Data S7) were defined as those expressed >5-fold higher in cTECs than in both mTEC^lo^ and mTEC^hi^ in bulk RNA-seq. E. UMAP projection highlighting mTECs expressing *CCL19* in normal and tumor mTECs from scRNA-seq. F. Normalized expression values of *Ccl19* in mTEC I and mTEC II of control, *Ascl1*^ΔTEC^, *Aire*^−/−^, and *Ascl1*^ΔTEC^*Aire*^−/−^ mice in bulk RNA-seq.

Among the commonly downregulated genes, CCL19 was markedly reduced in tumor-derived mTECs from thymoma patients (Fig. 8E) and in mTEC I and II subsets from *Ascl1*^ΔTEC^*Aire^−/−^*mice (Fig. 8F). Although CCL21 also showed a reduction trend (Fig. S25C), it did not reach statistical significance, likely due to its inherently low expression in normal human TECs. Since CCL19 and CCL21 attract positively selected T cells via the CCR7 receptor ^46, 47^, diminished CCR7 signaling may be a common feature of tumor mTECs in thymoma. In mice, *Ccl21a* was highly expressed in mTEC I and reduced by the deletion of both *Ascl1* and *Aire* (Fig. S25D), similarly suggesting impaired CCR7 signaling. Thus, disruption of the CCL19/CCL21–CCR7 axis appears to be a shared hallmark of human thymoma and the *Ascl1*^ΔTEC^*Aire^−/−^* mouse thymus. Overall, the combined upregulation of cTEC-associated genes and downregulation of CCL19 in tumor mTECs may result from ASCL1 loss alongside the aging-associated decline in AIRE.

## Discussion

Our study isolates and profiles EpCAM⁺ TECs from both tumor regions and spatially distinct non-tumorous areas within the thymus from thymoma patients. This TEC-specific strategy reveals ASCL1 downregulation as a tumor-associated alteration that was not detectable in bulk transcriptomic datasets or in scRNA-seq studies of total thymoma cells ^22, 29, 48^, where epithelial signals are diluted by abundant lymphocytes and “normal” thymic tissue is often derived from age-unmatched controls.

scRNA-seq analysis showed that tumor TECs and spatially separated normal TECs within the thymus from thymoma patients largely shared similar transcriptional profiles. Although we cannot fully exclude the possibility that tumor TECs exert distal influences on nearby TECs, integration with healthy donor datasets indicates that these non-tumorous regions retain a normal TEC subtype composition. Nevertheless, differential expression analysis revealed that tumor TECs displayed several tumor-associated alterations, including reduced expression of mitochondrial respiratory chain genes, consistent with a shift toward glycolytic metabolism, and decreased type I interferon–induced genes, which may support tumor cell survival. While these alterations reflect tumor biology, their direct relevance to central tolerance remains unclear. One notable exception was the marked reduction of CCL19 and CCL21 expression in tumor TECs. Because these chemokines are essential for guiding thymocytes into the medulla and enabling effective negative selection, their reduction represents a plausible mechanism that could contribute to tolerance breakdown in thymoma. Aside from this chemokine axis, our analysis did not detect additional tumor-specific changes in TECs that clearly account for the autoimmune manifestations associated with thymoma, highlighting the need for future studies with spatially resolved or functional assays.

ASCL1 is expressed at relatively high levels in immature mTECs in mice. Consistent with this, bulk RNA-seq analysis showed that deletion of *Ascl1* resulted in both up- and down-regulation of genes in immature mTECs. Single-cell ATAC-seq analysis revealed accompanying changes in chromatin accessibility, suggesting that ASCL1 expression correlates with differences in accessible chromatin regions. However, whether these accessibility changes reflect direct DNA binding by ASCL1 or indirect consequences of altered transcriptional networks remains unclear. In other cellular contexts, ASCL1 has been reported to interact with chromatin-associated complexes, including BAF/SWI-SNF components in neurons^49^ and histone deacetylases in mesodermal tissues ^50^, supporting the possibility that similar interactions may occur in TECs. Defining the molecular mechanisms through which ASCL1 influences gene expression and chromatin accessibility in mTECs will require further investigation.

Our data indicate that ASCL1 and AIRE exert both cooperative and context-dependent antagonistic effects in shaping gene expression in mature mTECs. Because ASCL1 binds specific DNA motifs ^40^, whereas AIRE lacks a canonical DNA-binding domain and instead associates with chromatin through histone interactions ^51^, direct mechanistic redundancy between the two factors is unlikely. Given the high expression of ASCL1 in immature mTECs, we propose a sequential model (Fig. S20). ASCL1 modulates chromatin accessibility at early stages, establishing regulatory states that influence later transcriptional outcomes. Regions opened in an ASCL1-dependent manner can sustain gene expression independently of AIRE in mature mTECs. Conversely, genomic regions that remain closed or are configured in an ASCL1-dependent manner become accessible through AIRE activity at later stages. In the absence of ASCL1, AIRE gains access to regions normally regulated by ASCL1, altering the repertoire of induced genes, including a subset of TSA genes. Thus, ASCL1 helps determine the spectrum of AIRE-responsive genes in mTECs, providing a framework to explain the distinct transcriptional profiles observed in *Aire^−/^* and *Ascl1*^ΔTEC^*Aire^−/−^* mice. The mechanisms governing AIRE recruitment to ASCL1-influenced regions remain to be defined.

A previous study suggested that Ascl1 regulates the development of neuroendocrine (Endo) TECs, a subset of post-Aire mTECs, which may help suppress autoimmunity against the stomach and thyroid ^13^. In contrast, our study focused on ASCL1 function in Ccl21⁺ and Aire⁺ mTECs. We observed that Ascl1 deletion causes autoimmune manifestations in the liver, kidney, and pancreas, suggesting that ASCL1 can regulate tolerance independently of Endo TECs.

Additionally, impaired negative selection in the thymus in *Ascl1*^ΔTEC^ mice provides further evidence that ASCL1 contributes to the elimination of autoreactive T cells. Combined deletion of Ascl1 and Aire expanded the range of affected organs compared with Aire single knockouts, potentially reflecting changes in tissue-specific antigen profiles. Loss of Ascl1 alone or with Aire also reduced expression of CCR7 ligands Ccl21a and Ccl19, which are important for thymocyte migration and localization ^47^, and may contribute to autoimmunity by disrupting thymocyte–TEC interactions. Furthermore, cTEC genes, including Prss16 and Ccl25, were aberrantly upregulated in mTECs from *Ascl1*^ΔTEC^ mice and tumor mTECs from thymoma patients, potentially affecting T cell selection. We note that the precise self-antigens responsible for these autoimmune responses were not determined, representing a limitation of this study. Overall, these findings suggest that ASCL1 shapes mTEC transcriptional programs and thymic tolerance, and future studies identifying ASCL1-dependent self-antigens will be essential to clarify the mechanistic links between mTEC dysfunction and autoimmunity.

Although *Ascl1*^ΔTEC^ and *Ascl1*^ΔTEC^*Aire^−/−^*mice exhibited autoimmune manifestations, they did not develop MG, indicating that loss of ASCL1 and AIRE alone is insufficient to model thymoma-associated MG. These findings suggest that disruption of ASCL1–AIRE–dependent transcriptional programs may prime the thymic environment by permitting the emergence of autoreactive T cells; however, additional factors are likely required to drive MG-associated autoantibody production. We therefore propose a two-hit model: the first hit involves loss of ASCL1 and AIRE, which increases susceptibility to autoimmunity, whereas the second hit may arise from tumor-associated alterations or external environmental triggers, such as infection or inflammatory signals, that promote activation and expansion of autoreactive T and B cells. This framework highlights the limitations of current mouse models and underscores the need to identify additional mechanisms that precipitate MG in the context of thymoma.

## Materials & Methods

### Human thymoma samples

Thymic samples, thymoma, and thymus tissue were obtained from thymoma patients who underwent thymo-thymectomy at the Department of Thoracic Surgery, The University of Tokyo Hospital. All samples were collected and used in accordance with Ethical Guidelines for Life Science, Medical and Health Research Involving Human Subjects and in agreement with the Research Ethics Committee of RIKEN (Approved number: H30-26(6)) and the Research Board of the University of Tokyo Graduate School of Medicine (Approved number: 2018192G-(2)).

All patients signed a written informed consent form. Immediately after surgery, thymomas and normal thymus glands were placed in RPMI 1640 cell medium supplemented with 5% FBS and processed in a biosafety level 2 laboratory in the next five hours for cell preparation.

### Mice

*Asc1*^gfp^ mice were purchased from Jackson Laboratory (Bar Harbor, ME, USA) (Stock No. 012881) and backcrossed with C57BL/6 and Balb/c mice for > 10 generations. Littermates or age-matched wild-type mice from the same colonies as mutant mice were used as controls. *Ascl1* flox mice were established by recombination in JM8.A3 ES cells (Fig. S2) and subsequent injection into blastocysts. Of these, two mouse lines were established from two independent ES cell clones. *Foxn1*-Cre mice were reported previously (*62*). *Aire*^−/−^ mice (CDB0479K, http://www2.brc.riken.jp/lab/animal/detail.php?brc_no=03515) were provided by the RIKEN BRC through the National Bio-Resource Project of the MEXT, Japan. Balb/c A nu/nu mice were purchased from Clea Japan (Tokyo, Japan). *Rag1*^−/−^ (B6.129S7-Rag1tm1Mom/J, stock No. 002216) mice were from Jackson Laboratory. All mice were maintained under specific pathogen-free conditions and handled in accordance with Guidelines for Animal Experiments of the Institute of Medical Science, University of Tokyo (Tokyo, Japan) and the Institutional Animal Care and Use Committee (IACUC) of RIKEN Yokohama Branch. Embryonic day 0.5 (E 0.5) was defined as the first morning a vaginal plug was observed.

### Antibodies and reagents

Antibodies used included FcR Blocking Reagent human (Miltenyi, Cat#130-059-901), APC-anti-human CD45 (BioLegend, clone HI30, Cat#304011), PE-anti-human CD326(EpCAM) (BioLegend, clone 9C4, Cat#324205), FITC-anti-human CD235a (BechmanCoulter, clone 11E4B-7-6 (KC16), Cat#IM2212U), FITC-anti-human CD4 (BioLegend, clone OKT4, Cat#317407), APC-anti-human CD8a (BioLegend, clone HIT8a, Cat#300911), PE-anti-human TCRa/b (BioLegend, clone IP26, Cat#306707), anti-mouse CD16/32 (BioLegend, clone 93, Cat#101302), FITC anti-mouse CD3e (BioLegend, clone 145-2C11, Cat#100306), APCCy7-anti-mouse CD45 (BioLegend, clone 30 F-11, Cat#103116), APCCy7-anti-mouse TER119 (BioLegend, clone TER-119, Cat#557853), FITC-anti-mouse EpCAM (CD326) (BioLegend, clone G8.8, Cat#118208), BV510-anti-mouse EpCAM (CD326) (BioLegend, clone G8.8, Cat#118231), BV421-anti-mouse EpCAM (CD326) (BioLegend, clone G8.8, Cat#118225), PE-anti-mouse CD80 (ThermoFisher eBioscience, clone 16-10A1, Cat#12-0801-81), Alexa647-anti- mouse Ly51 (BioLegend, clone 6C3, Cat# 108312), PerCP/Cyanine5.5-anti-mouse Ly51 (BioLegend, clone 6C3, Cat# 108315), AlexaFluor647-anti-mouse L1CAM (Novus, clone 555, Cat# FAB5674R), PECy7-anti-mouse IA/IE (BioLegend, clone M5/114.15.2, Cat# 107630), PECy7-anti-mouse IA/IE (BioLegend, clone M5/114.15.2, Cat# 107630), AlexaFluor700-anti-mouse IA/IE (BioLegend, clone M5/114.15.2, Cat# 107621), FITC-anti-mouse Ly6d (BioLegend, clone 49-H4, Cat# 138605), PE-anti-mouse Ly6d (BioLegend, clone 49-H4, Cat# 138603), PE-anti-mouse CD104(ITGB4) (BioLegend, clone 346-11A, Cat# 123609), PECy7-anti-mouse CD104(ITGB4) (BioLegend, clone 346-11A, Cat# 123616), BV711-anti-mouse CD104(ITGB4) (BioLegend, clone 346-11A, Cat# 743082), Biotinylated UEA-1 (Vector Laboratories, Cat#B-1065), PECy7-streptavidin (ThermoFisher eBioscience, Cat# 25-4317-82), APC-streptavidin (BioLegend, Cat# 405207), PerCP/Cyanine5.5-streptavidin (BioLegend, Cat# 405214), Alexa546-streptavidin (ThermoFisher Invitrogen, S11225), BV605-streptavidin (BioLegend, Cat# 405229), PECy7-anti-mouse CD4 (BioLegend, clone RAM4-4, Cat#116016), Alexa647-anti-mouse CD4 (BioLegend, clone RAM4-5, Cat#100533), BV510-anti-mouse CD4 (BioLegend, clone RAM4-5, Cat#100553), FITC-anti-mouse CD4 (ThermoFisher eBioscience, clone GK1.5, Cat# 11-0041-85), APC-anti-mouse CD8a (BioLegend, clone 53-6.7, Cat# 100712), APCCy7-anti-mouse CD8a (BioLegend, clone 53-6.7, Cat# 100722), FITC-anti-mouse CD25 (BioLegend, clonePC61, Cat# 102005), PE-anti-mouse CD25 (BioLegend, clonePC61, Cat# 102007), PE-anti-mouse Foxp3 (ThermoFisher eBioscience, clone FJK-16s, Cat# 12-5773-82), PE-anti-mouse CD44 (BioLegend, clone IM7, Cat# 103007), BV785-anti-mouse CD44 (BioLegend, clone IM7, Cat# 103059), PE-anti-mouse TCRgd (BioLegend, clone GL3, Cat# 118108), Alexa647-anti-mouse H2Kb (BioLegend, clone AF6-88.5.5.3, Cat# 116511), FITC-anti-mouse CD62L (BioLegend, clone MEL-14, Cat# 104406), Pacific Blue-anti-mouse CD11b (BioLegend, clone M1/70, Cat# 101224), Pacific Blue-anti-mouse CD11c (BioLegend, clone N418, Cat# 117322), Pacific Blue-anti-mouse F4/80 (BioLegend, clone BM8, Cat# 123124), Pacific Blue-anti-mouse B220 (BioLegend, clone RA3-6B2, Cat# 103227), anti-mouse Ascl1 (R&D, goat polyclonal, Cat# AF2567), purified anti-mouse Keratin-5 (BioLegend, rabbit polyclonal, Cat# 905504), eFluor660-anti-mouse Aire (ThermoFisher eBioscience, clone 5H12, Cat# 50-5934-82), AlexaFluor488-anti-mouse Aire (ThermoFisher eBioscience, clone 5H12, Cat# 53-5934-82), AlexaFlor488-donkey anti-goat IgG (Invitrogen, Cat# A10055), AlexaFluor546-donkey-anti-rabbit IgG (ThermoFisher Invitrogen, Cat# A11040), AlexaFluor 488-goat-anti-mouse IgG (ThermoFisher Invitrogen, Cat# A11001), AlexaFluor647-anti-mouse Keratin 5 (Abcam, Cat#193895), DAPI (ThermoFisher Invitrogen, D1306), Propidium Iodide (Fujifilm-Wako, Cat#169-26281). APC and PE-labeled GFP peptides:A^b^ tetramers , Phycoerythrin (PE)- CD1d tetramers loaded with or without PBS57 were provided by the Tetramer Core Facility of the US National Institutes of Health. BV785-anti-mouse CD4 (BioLegend, clone RM4-5, Cat#100551), BV711-anti-mouse CD8a (BioLegend, clone 53-6.7, Cat#100759), BV605-anti-mouse CD25 (BioLegend, clone PC61, Cat#102036), PerCPCy5.5 anti-mouse CD24 (BioLegend, clone M1/69, Cat#101823), AlexaFluor700 anti-mouse CD44 (BioLegend, clone IM7, Cat#103026), FITC-anti-mouse CD69 (BioLegend, clone H1.2F3, Cat#104506), PECy7-anti-mouse CD197 (Ccr7) (BioLegend, clone 4B12, Cat#120124), BV421-anti-mouse CD279 (PD-1) (BioLegend, clone 29F.1A12, Cat#135221), PE-eFluor610 anti-mouse Foxp3 (ThermoFisher eBioscience Invitrogen, clone FJK-16s, Cat#61-5773-82), PE-anti-mouse/human Helios (BioLegend, clone 22F6, Cat#137216), BV605-anti-mouse NK1.1 (BioLegend, clone PK136, Cat#108739), APCFire750-anti-mouse TCRb (BioLegend, clone H57-597, Cat#109233)

### Preparation of TECs from human thymomas and normal thymus

Thymic epithelial cells (TECs) were obtained from type B thymomas, as described in Tumor Dissociation Kit (Miltenyi Biotec) protocols with modifications. A piece of tumor (about 100 mg) was placed in ice cold PBS. Fat and fibrous and necrotic areas were removed from samples. Tumor samples were cut into small pieces of 1-2 mm, and transferred into gentleMACS C Tubes with 2 mL of digestion Liberase solution (RPMI 1640 medium supplemented with 0.05 U/mL Liberase (Roche) and 0.1 mg/mL DNase I). After incubation at 37°C in a water bath for 5 min and another incubation at 37°C in the shaker for 7 min under gentle agitation, samples were dissociated by running the program h_tumor_03. Incubation at 37°C in the shaker for 12 min with gentle agitation was repeated twice, running the program h_tumor_03. Supernatants were collected, mixed (v/v) with 1X PBS supplemented with 2% FBS, and 1 mM EDTA to stop the enzymatic digestion, and centrifuged at 350 x g for 5 min at 4°C.

Preparation of single-cell suspensions from samples of normal thymus was performed according to a previous procedure(*63*) with modifications. Tissue pieces were transferred into a gentleMACS C tube (Miltenyi) containing 2 mL of digestion Liberase solution. The gentleMACS program m_spleen_02 was run five times. After incubation for 12 min with gentle agitation, the first digested supernatant was separated by centrifugation at 110 x g for 2 min at room temperature. Supernatant was collected into a tube containing cold FACS buffer (2% FBS in PBS) with 1 mM EDTA, and replaced with fresh 2 mL of digestion Liberase solution. The gentleMACS program m_spleen_01 was run once. After incubation for 12 min, tubes were centrifuged at 110 x g for 2 min at room temperature to remove undigested fragments. Cells were pooled with the supernatant from the previous round of digestion, and centrifuged at 350 x g for 5 min at 4°C. All samples were treated with 2 mL of RBC lysis buffer (Biolegend) for 5 min prior to use for FACS analysis. Cells were counted in AO/PI using a Luna automated cell counter (Logos Biosystems).

### Preparation of TECs from mice

Thymi from 4-week mice were dissected and minced with razor blades. Thymic fragments were pipetted up and down 30 times in 1 mL of Hepes-RPMI (Wako) to break up clumps. Supernatants were taken as lymphocytes, and the remaining fragments were digested in RPMI 1640 medium containing Liberase (Roche) at 0.05U/mL plus DNase I (Sigma-Aldrich) at 0.01% w/v by incubation at 37 °C for 12 min three times, with pipetting after each incubation. Reactions were stopped by addition of the same volume of FACS buffer containing 1 mM ethylenediaminetetraacetic acid and washing with FACS buffer. Total cells were calculated as the sum of lymphocytes plus digested cells.

### Flow cytometric analysis

Single-cell suspensions were stained with anti-human, or anti-mouse antibodies as described above for 20 min on ice after blocking. Dead cells were excluded using 7-aminoactinomycin D (Calbiochem), or Sytox Blue (Invitrogen) staining. Cells were sorted using a FACS Aria instrument (BD). Post-sorted cells were determined to have >95% of relevant fractions.

Regulatory T cells were stained using eBioscience™ Foxp3 / Transcription Factor Staining Buffer Set (Invitrogen, Cat# 00-5523-00). Data were analyzed using Flowjo 10.

### Fetal thymus organ culture (FTOC) and transplantation into nude mice

Thymic lobes were isolated from E14.5–15.5 embryos and cultured on Nucleopore filters (Whatman) placed in R10 medium consisting of RPMI 1640 (Wako), 10% fetal bovine serum (Equitech-Bio), 2 mM L-glutamine (Wako), 100 U/mL penicillin (Banyu Pharmaceutical), 100 µg/mL streptomycin (Meiji Seika Pharma Co., Ltd.), and 50 µM 2-mercaptoethanol (Wako) with 1.35 mM 2′-deoxyguanosine (2-DG; Sigma-Aldrich) for 4 days (2DG-FTOCs). For transplantation experiments, 2DG-FTOCs on a Balb/c background were further cultured for one day in R10 medium without 2-DG to eliminate 2-DG prior to transplantation. 2DG-FTOCs on a Balb/c background were transplanted to kidney capsules of congenic 6-week female nude mice.

### Quantification of GFP-specific T cells

Mice were i.p. injected at two sites with 100µg of GFP peptides (HDFFKSAMPEGYVQE)(BEX CO., LTD.) emulsified 1:1 in PBS/complete Freund’s adjuvants (Rockland Immunochemicals, Inc.). Whole spleens were collected and analyzed seven days after i.p. immunization. All splenocytes was doubly stained with 10nM APC and PE-labeled GFP peptides:A^b^ tetramers (NIH Tetramer Core) for 1 hr at room temperature in DMEM plus 2% FCS and 50nM dasatinib (TargetMol). Then, one tenth cells were taken as samples before MACS column, and others were incubated with anti-PE (Ultra pure) and anti-APC microbeads (Milteny) for 15min on ice and passed through MACS LS column. The samples of tetramer enriched (and unenriched) fractions were stained with B220, CD11b, CD11c, F4/80, CD3e, CD4, and CD44 (Biolegend) . The population and number of GFP-specific CD4+ T cells were quantified by flowcytometry (gating strategy: B220^-^ CD11b^-^ CD11c^-^ F4/80^-^ CD3e ^+^ CD4^+^ CD44^+^).

### Histological examination and detection of autoantibodies

Organs from nude mice were stained with hematoxylin and eosin. Degrees of inflammatory cell infiltration were scored as follows: 0, below one focus of perivascular infiltration; 1, several foci of perivascular infiltration; 2, cellular infiltration in less than 50% of the vasculature; 4, cellular infiltration in more than 50% of the vasculature. To detect autoantibodies in serum, tissue samples from *Rag1*^−/−^ mice were embedded in OCT compound, frozen in liquid N_2_, and sectioned into 5-µm slices. Cryosections were fixed with ice-cold acetone for 5 min and washed with phosphate buffered saline (PBS). Samples were blocked with 10% goat serum in PBS (Thermo Fisher Scientific) and incubated with serum (100× dilution) for 1 h at room temperature. Slides were subsequently incubated with secondary antibody (Alexa Fluor 488-goat-anti-mouse IgG) and 1 µg/mL propidium iodide for 40 min at room temperature. Confocal color images were obtained using an FV1000D (Olympus) or LAS X (Leica) confocal microscope, and MFI of the images were counted. Relative intensities of MFI were calculated as the mean of control=1.

### Immunohistochemistry

For immunohistochemical staining, frozen samples in OCT compound were cut into 5-µm slices. Cryosections were fixed for 5 min in acetone or 4% paraformaldehyde in PBS on ice. After washing with PBS three times, sections were blocked with 10% normal goat serum. When using the goat anti-mouse Ascl1 antibody, 1% normal donkey serum, 1% BSA, and 0.1% Triton-X solution were used for blocking. Ascl1 was detected using a polyclonal goat anti-mouse Ascl1 antibody, followed by a secondary AlexaFluor 488-donkey anti-goat IgG. Keratin-5, Aire and Ly51 were detected using a labeled monoclonal antibody (1:300). Biotin UEA-1 and AlexaFluor 546-streptavidin (Invitrogen) were used to detect UEA-1 binding protein. Images were obtained with a BZ-X710 All-in-One Fluorescence Microscope (Keyence) and analyzed using a BZ-X (Keyence). Confocal color images were obtained with a TCS SP8 (Leica) and analyzed with a LAS X (Leica).

### qRT-PCR

Total RNA was extracted using RNeasy Micro kits (Qiagen) or TRIzol (Invitrogen). cDNA was obtained by random-primed reverse transcription using a Primescript II first strand cDNA synthesis kit (TaKaRa Bio Inc.). Real-time qPCR was performed using CFX connect^™^ (BioRad) and SYBR Green Master Mix (TOYOBO). Primers used were *36B4*: 5′- TCC AGG CTT TGG GCA TCA-3′ and 5′- CTT TAT CAG CTG CAC ATC ACT-3′; *Ascl1*: 5′- ACT TTG GAA GCA GGA TGG CAG CA-3′ and 5′- AGG TGC CCC TGT AGG TTG GCT-3′.

### Single-Cell Sequencing Preparation

Human thymic epithelial cells (EpCAM^+^CD45^−^CD235a^−^) or stroma cells (CD45^−^CD235a ^−^) were sorted from cell suspensions from tumor and normal regions. Pooled mouse TECs (EpCAM^+^CD45^−^TER119^−^) from two or three mice were sorted with a cell sorter (BD FACS Aria Illu). Single-cell emulsions were obtained using a 10x Genomics Controller and a v2 Library and Gel Bead kit (10X Genomics). RNA-sequencing libraries were prepared according to the 10x 3′ v2 kit protocol. At least two scRNA-sequencing libraries were obtained from each genotype. Resulting libraries were sequenced on an Illumina HiSeq. FASTQ files were processed using Fastp(*64*). Reads were demultiplexed and mapped to the mm10 reference genome with Cell Ranger (v3.0.0). Digital expression matrices were prepared by counting unique molecule identifiers (UMI). Downstream single-cell analyses, i.e., integration of two datasets, correction of dataset-specific batch effects, UMAP dimension reduction, cell cluster identification, conserved marker identification, and regressing out cell cycle gens) were done using the Seurat package (v3.0) (*65*). Cells that contained >15% mitochondrial transcripts were filtered out. Genes expressed in more than 3 cells and cells expressing at least 200 genes were selected for analysis.All scRNA-seq data were integrated with previously reported age-matched wild-type data(*11*) using canonical correlation analysis (CCA), and clustering and UMAP dimension reduction were performed using standard settings of Seurat package

### RNA-Seq analysis

Total RNA was extracted using TRIzol reagent according to the manufacturer’s protocol (Thermo Fisher Scientific). After depletion of rRNA using the NEBNext rRNA Depletion Kit, the RNA-Seq library was prepared by using the NEBNext Ultra Directional RNA Library Prep Kit (New England Biolabs). For RNA-seq analysis of mTECs I to IV, random displacement amplification sequencing (RamDA-seq) was used (*66*). Briefly, up to 2000 sorted cells were lysed in 2X TCL buffer (Qiagen). Nucleic acids were purified with Agencourt RNA Clean XP (Beckman Coulter), and then treated with DNase I, an RT-RamDA mixture containing 2.5x PrimeScript Buffer (Takara), 0.6 µM oligo(dT)18 (Thermo), 10 µM RT primer mix, 100 µg/mL of T4 gene 32 protein, and 3× PrimeScript enzyme mix (Takara) for reverse transcription.

Second-strands were synthesized by mixing with 2× NEB buffer 2 (NEB), 625 nM dNTP Mixture (NEB), 25 µM second primers, and 375 U/mL of Klenow Fragments (3’-5’ exo-) (NEB). After purification of double-stranded cDNAs by AMPure XP (Beckman Coulter), sequencing library preparation was performed using the Tn5 tagmentation-based method. Paired-end sequencing was carried out with a NextSeq500 (Illumina). Sequenced reads were quantified for annotated genes using the CLC Genomics Workbench (Version 7.5.1; Qiagen). Transcription expression levels were estimated as “Normalization values” determined with CLC Genomics Workbench or CLC Main Workbench. Expression values of genes were cut off at a threshold of 1 Normalization value. Differential expression was analyzed by empirical analysis with the DGE (edgeR test) tool in CLC Main Workbench, in which the Exact Test of Robinson and Smyth was used (*67*). A false discovery rate (FDR)-corrected P-value was used for testing statistics. Principle component analysis (PCA) was performed using the prcomp function in R-project.

### Droplet-based scATAC-seq analysis

Cell suspensions of thymi from 2 or 3 4-week-old mice were prepared and pooled for each individual scATAC-seq experiment(*11*). The EpCAM^+^CD45^−^TER119^−^ fraction was collected using a cell sorter (BD Aria) more than 10^5^ cells. After washing with PBS containing 0.04% BSA, cells were suspended in lysis buffer (10mM Tris-HCl (pH 7.4), 10 mM NaCl, 3 mM MgCl_2_, 0.1% Tween-20, 0.1% NP-40, 0.01% Digitonin, and 1% BSA) on ice for 3 min. After washing by Wash buffer (10mM Tris-HCl (pH 7.4), 10 mM NaCl, 3 mM MgCl_2_, 0.1% Tween-20, and 1% BSA) twice, the concentration of nuclei and their viability were determined by staining with AO/PI (LogosBiosystems). Just 10,000 nuclear suspensions were loaded onto a Chromium instrument (10× Genomics) to generate a single-cell emulsion. scATAC-seq libraries were prepared using Chromium Next GEM Single Cell ATAC Reagent Kits v1.1 and sequenced in multiplex on an Illumina Hiseq X ten platform. Reads were demultiplexed and mapped to the mm10 reference genome with Cell Ranger ATAC. Processing of data with the Cell Ranger pipeline was performed using an NIG supercomputer at ROIS National Institute of Genetics. Heatmap of ATAC-seq signal centered on ASCL1 motifs was prepared using deepTools ^52^.

### Pre-processing for analysis of snapATAC

FASTQ file libraries by paired-end sequences were assigned to each cell and aligned to the mm10 genome reference using Cell Ranger ATAC program to generate fragment file, barcode file and peak file. FASTQ files of each sample were converted to snap files using SnapTools retaining sequence reads with mapping quality score higher than 30 Cells with at least 10^3.5^ unique fragments and at least 20% of their reads in promoters were only retained to remove low quality cells. Most downstream analysis were performed using SnapATAC package in R. Reads were binned into the genome of 5kb windows to make a bins-by-cells binary matrix. Bins overlapping ENCODE-defined blacklist regions and bins with extremely high or low coverage were filtered out. Finally, cells with coverage of over 1000 bins were used for further downstream analysis. A bins-by-cells matrix was converted into a cells-by-cells matrix of the Jaccard similarity Index between two cells. Due to considering sequencing depth, this similarity matrix was normalized using a regression-based method. Dimensionality reduction was performed using the Nyström method and diffusion maps. The top 16 eigenvectors were retained for downstream analysis. Cell clustering was performed with Louvain graph-based algorithm on K Nearest Neighbor (KNN) Graphs (k = 30), followed by 2D UMAP visualization. To assign scATAC-seq clusters based on scRNA-seq datasets, snap files were converted into Seurat objects to use FindTransferAnchors and TransferData functions for cell type prediction. Differential accessible regions were identified using runDAR function and raw read counts, after peak calling by runMACS function in the SnapATAC package with the options ‘—nomodel --shift 100 --ext 200 --qval 5e-2 -B --SPMR’. Heatmap of ATAC-seq signal centered on ASCL1 motifs was prepared using deepTools ^52^.

### Transfection of HEK293 cells

HEK293 cells plated in 6-cm dishes were transfected with 2.5 µg of plasmid DNA (pMXs-mouse Ascl1-IRES-GFP and/or pMXs-mouse Aire-IRES-mCherry) using 5 µL polyethyleneimine (PEI). After 48 hours, GFP⁺, mCherry⁺, and GFP⁺mCherry⁺ cells were sorted by flow cytometry (BD Aria II). Total RNA was extracted from sorted cells and processed for RNA-seq as described above.

### Dimension reduction and cell clustering

For nonlinear dimensionality reduction, a Nyström landmark diffusion maps algorithm was applied for the bins-by-cells matrix. Using the top 20 eigenvectors determined by an ad hoc method, 2D UMAPs were generated for visualization and Louvain clustering was performed on the k-nearest neighbor graph (k = 30).

### Statistical analysis

P-values were calculated using Student’s t-test with two-tailed distribution or the Mann-Whitney U test. For the multiple comparisons, Tukey-Kramer test, Steel-Dwass test, and Brown-Forsythe and Welch ANOVA test were used.

