## Supplementary_Materials for "ASCL1 primes medullary thymic epithelial cell programs required for central tolerance"

### Chromatin organizer ASCL1 governs gene programs in thymic epithelial cells, defining immunological self

Nobuko Akiyama *et al.*

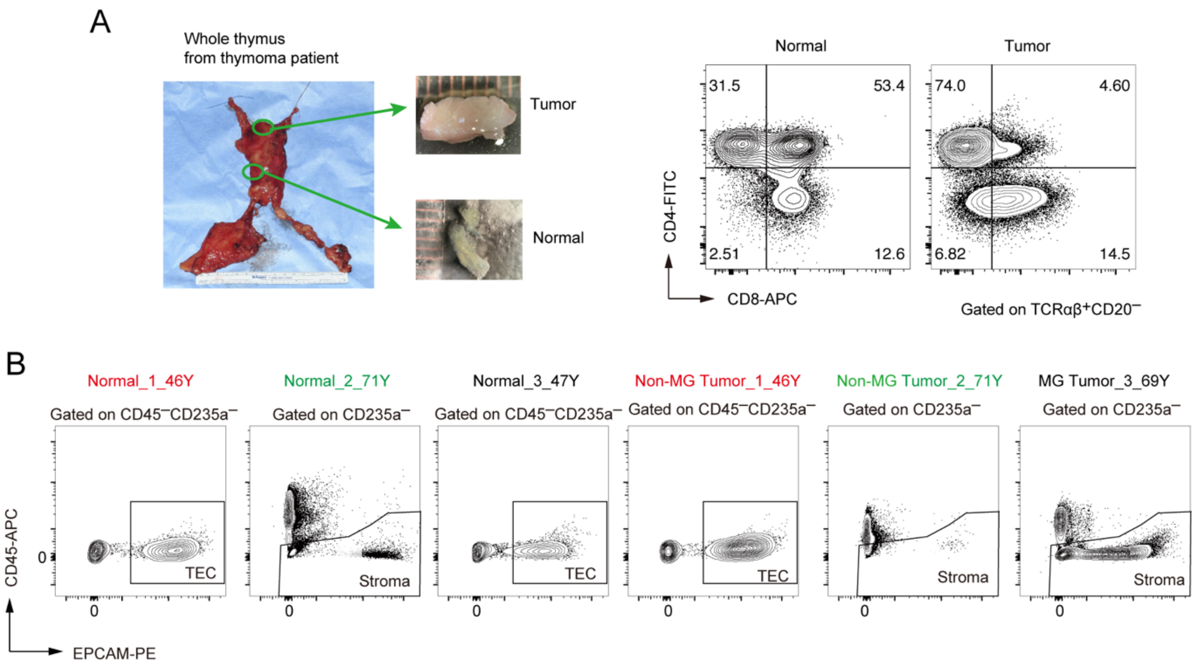


**Fig. S1. Isolation of stroma and thymic epithelial cells (TECs) from tumor and non-tumor regions in human thymoma.**

A. Photos of typical specimens from tumor and non-tumor regions (normal) of thymoma patients are presented. Typical flow cytometric profiles of CD4 and CD8-expressing cells gated on $TCR\alpha\beta^{+}CD20^{-}$ are shown in the right panels. These analyses were performed on cell suspensions prepared from both non-tumor and tumor regions.

B. The gating strategies used for sorting TECs and stromal cells from individual samples are presented. In the case of normal _2_71Y, Tumor_2_71Y, and Tumor_3_69Y_MG samples, stromal cell fractions, including TECs, were sorted and utilized for single-cell analysis.

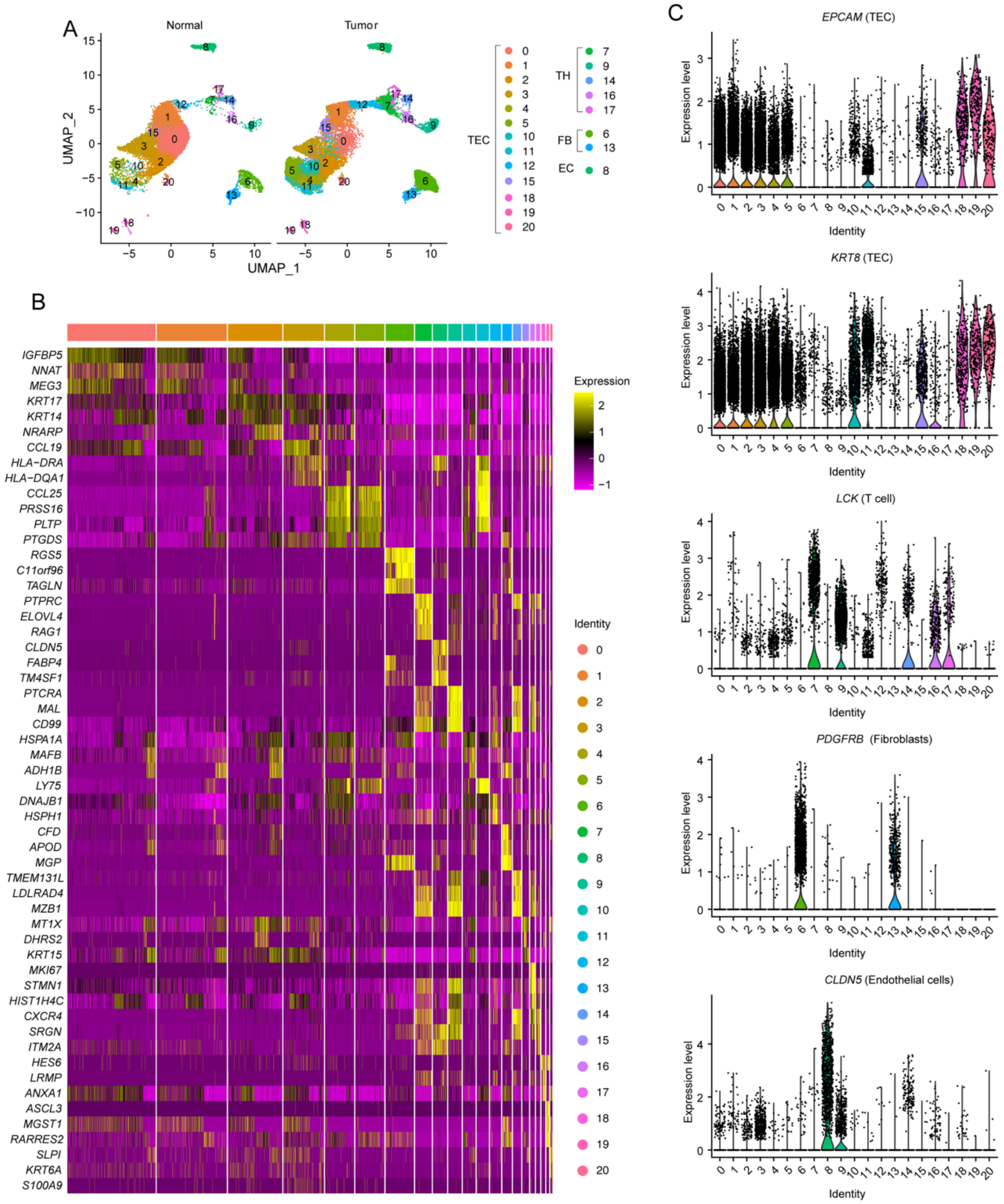


**Fig. S2. Detection of TECs in cell populations sorted from human thymoma**

A. UMAP projection for scRNA-seq data of all sorted cells from non-tumor and tumor regions of thymi from human thymoma patients is shown. TEC, TH, FB, and EC show clusters assigned as TECs, T cells, fibroblasts, and endothelial cells, respectively. These assignments were based on marker gene expressions shown in B and C.

B. A heat map displaying the top three differentially expressed genes in individual clusters is exhibited.

C. Violin plots for expression of typical marker genes.

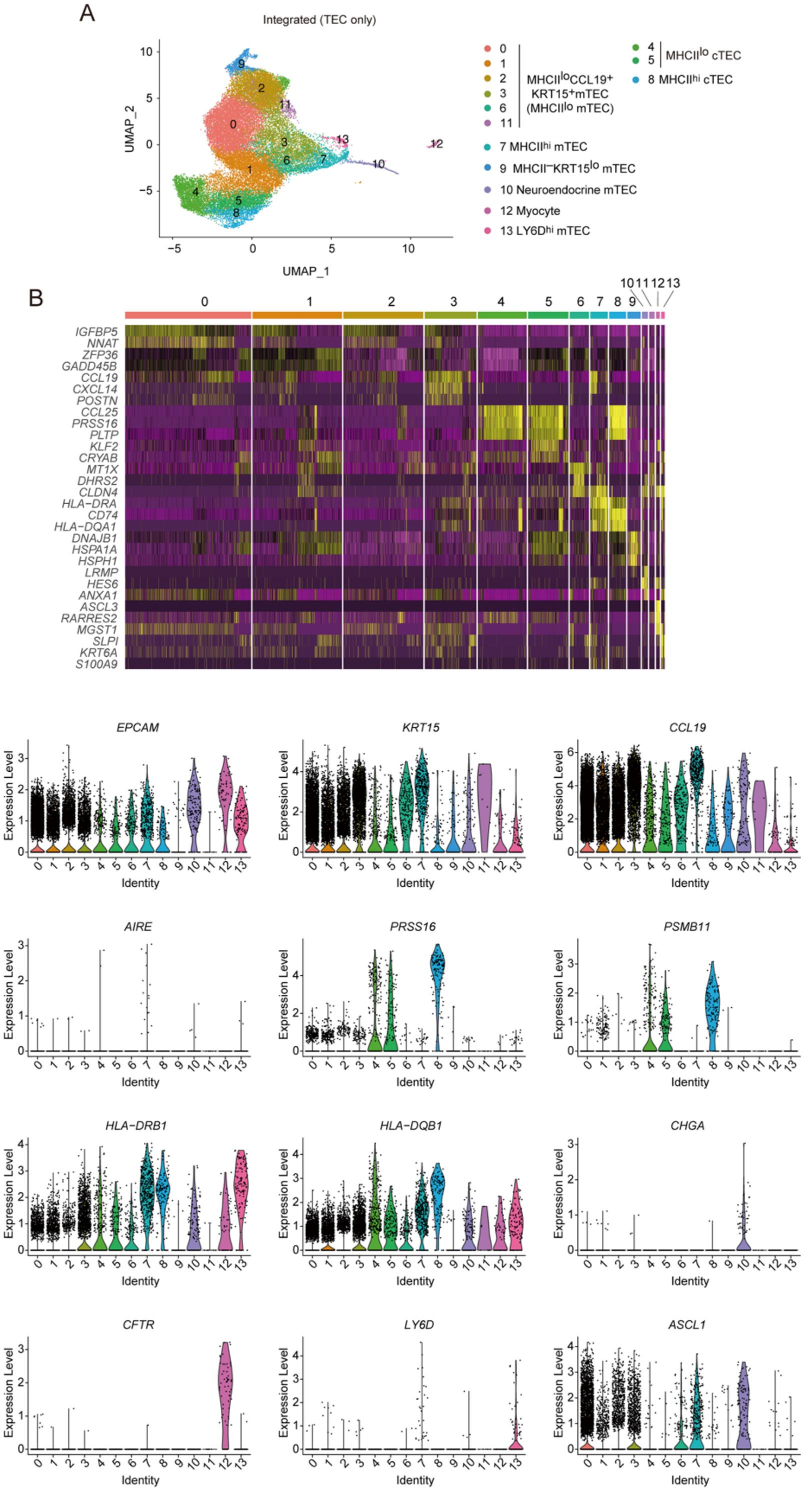


**Fig. S3. Assignment of TEC subsets from human thymoma.**

A. Integrated UMAP projection for scRNA-seq data of TECs from non-tumor and tumor regions of human thymoma.

B. Heatmap of top 3 differentially expressed genes of each cluster in Fig S3A.

C. Violin plots for expression of genes used for assignment of TEC subsets.

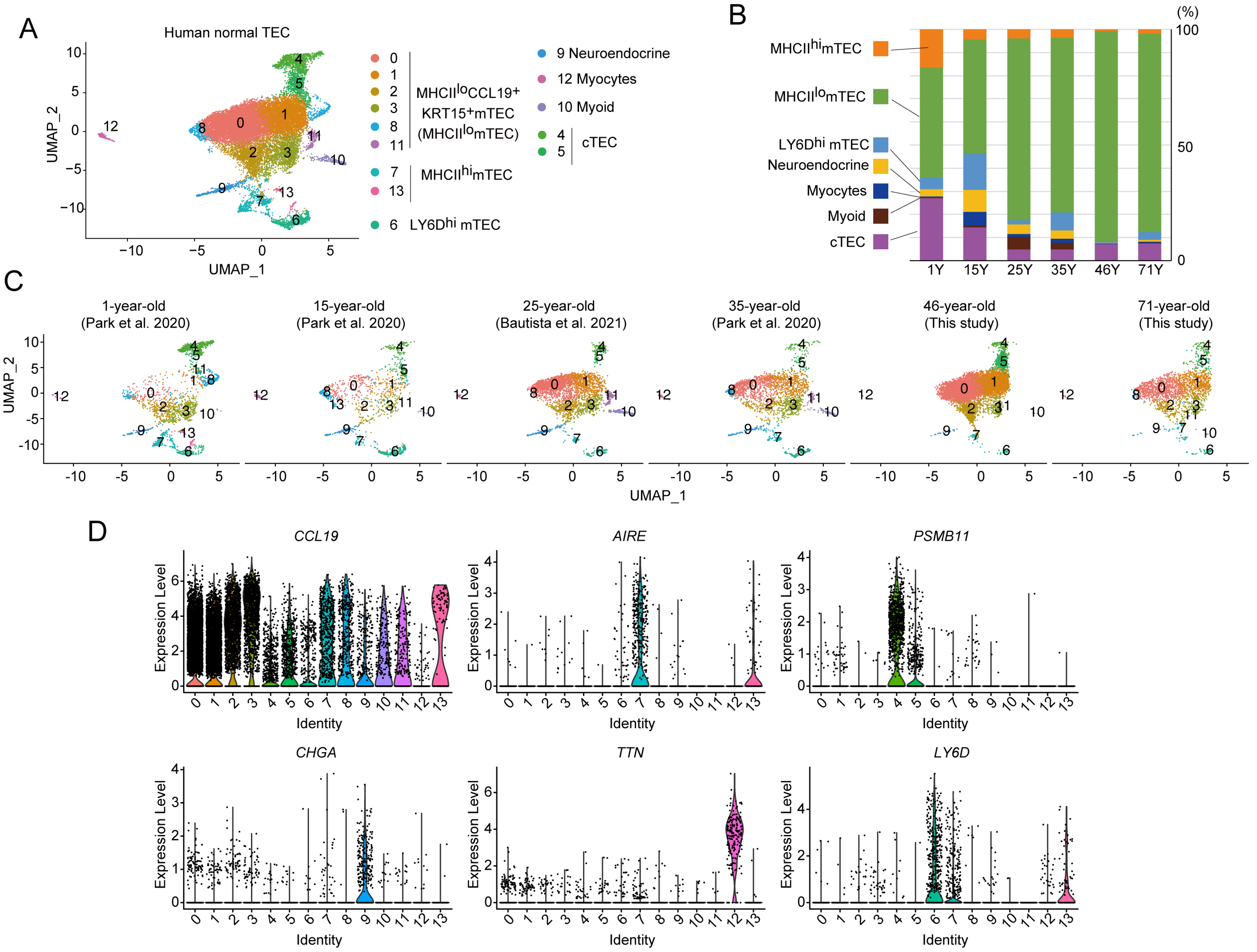


**Fig. S4. Integration of normal human TECs**

A. UMAP projection of scRNA-seq data from normal human TECs. Data from normal TECs of individuals aged 46 and 71 years in this study were integrated with previous datasets from younger human TECs (1, 15, 25, and 35 years old). Each cluster was assigned to indicated subsets according to expression levels of marker genes.

B. The panel represents the percentage of individual subsets in total TECs for each sample.

C. Individual UMAP projections of normal human TECs from each sample.

D. Typical violin plots for expression of genes used for assignment of human TEC subsets.

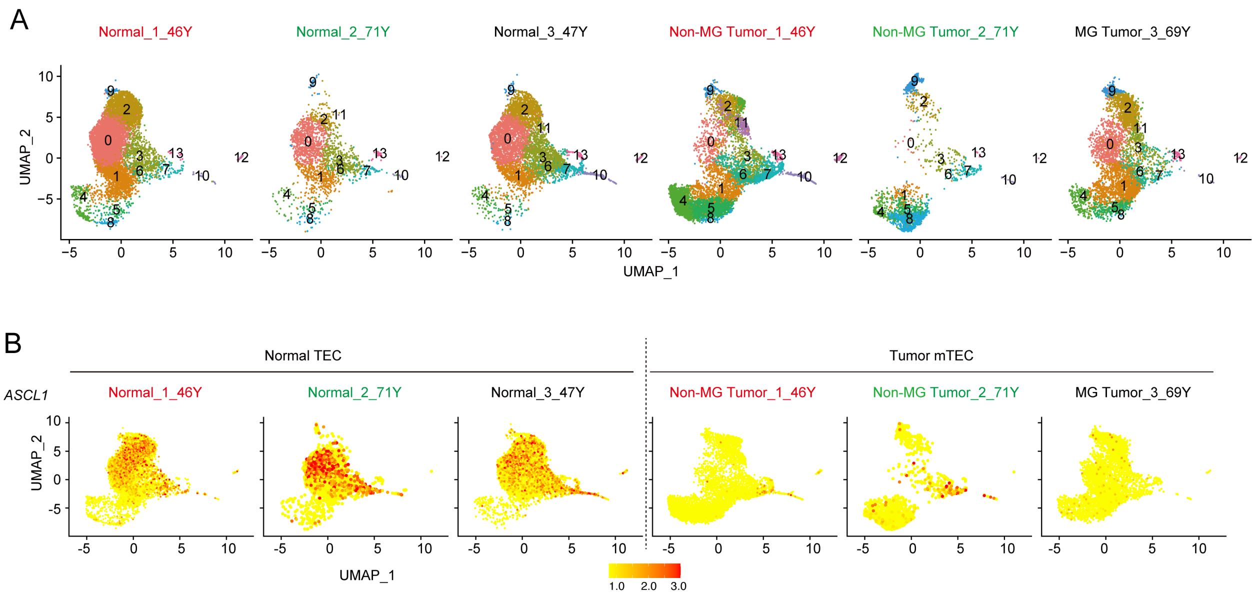


**Fig. S5. UMAP projection for scRNA-seq data of TECs**

A. UMAP projections for TEC clusters of individual patients are exhibited.

B. UMAP visualizations of *ASCL1* expression of individual patients are exhibited.

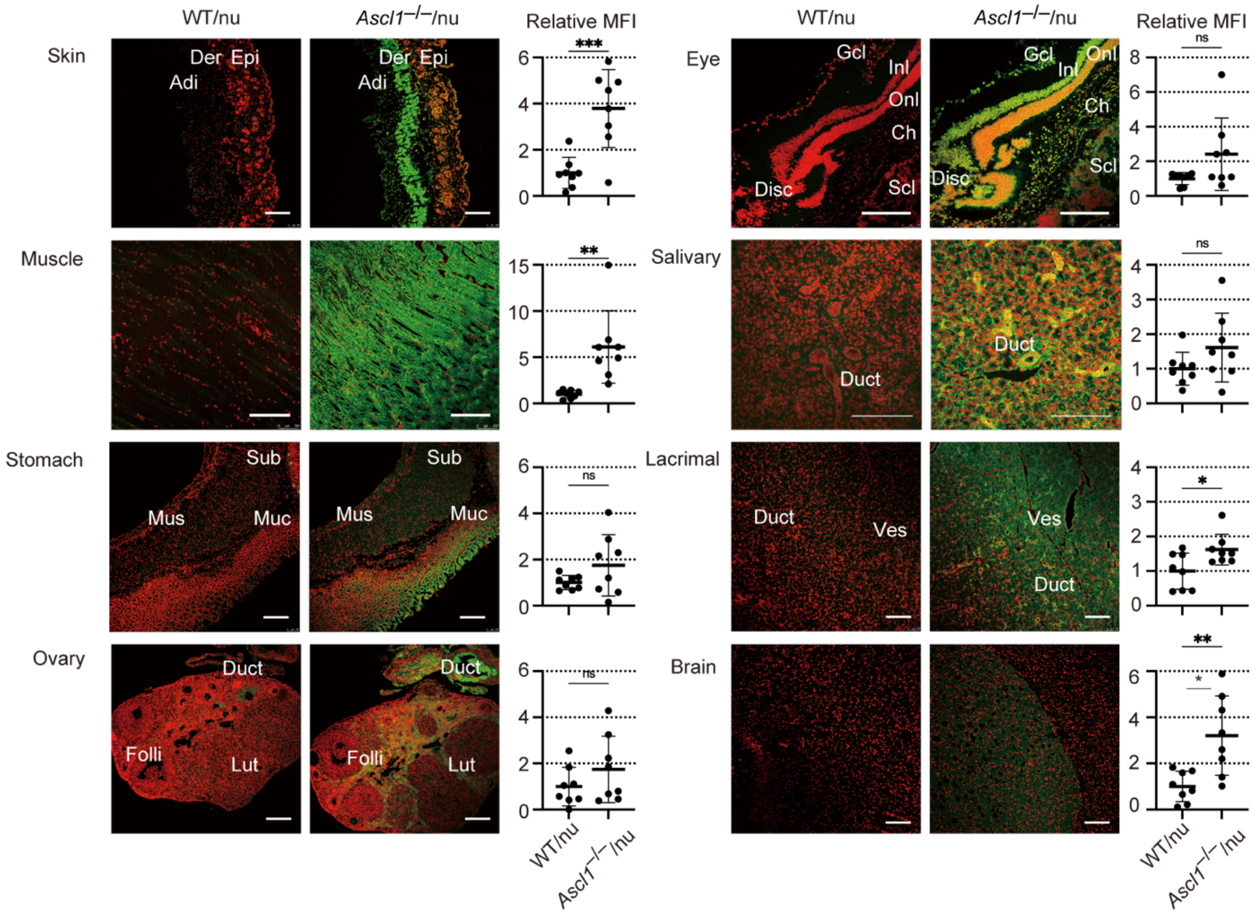


**Fig. S6. Detection of autoantibodies in thymus-grafted nude mouse serum.**
Organs from *Rag1*$^{-/-}$ mice were stained with sera (green) from nude mice grafted with control and *Ascl1*$^{-/-}$ thymic stroma and propidium iodide (red); Skin, Epi: epidermis, Der: dermis, Adi: adipose tissue. Stomach, Mus: muscle layer, Sub: submucosa, Muc: mucosal layer. Ovary, Folli: follicle, Lut: luteum, Duct: oviduct. Eye, Gcl: ganglion cell layer, Inl: inner nuclear layer, Onl: outer nuclear layer, Ch: choroid, Scl: sclera. Lacrimal, Ves: blood vessels. Scale bars, 200 µm. Right plots, relative MFI/pixel of autoantibodies (N=8 each, WT/nu and KO/nu); center line, median; box limits, upper and lower quartiles; whiskers, minimum and maximum range; points, individual animals.

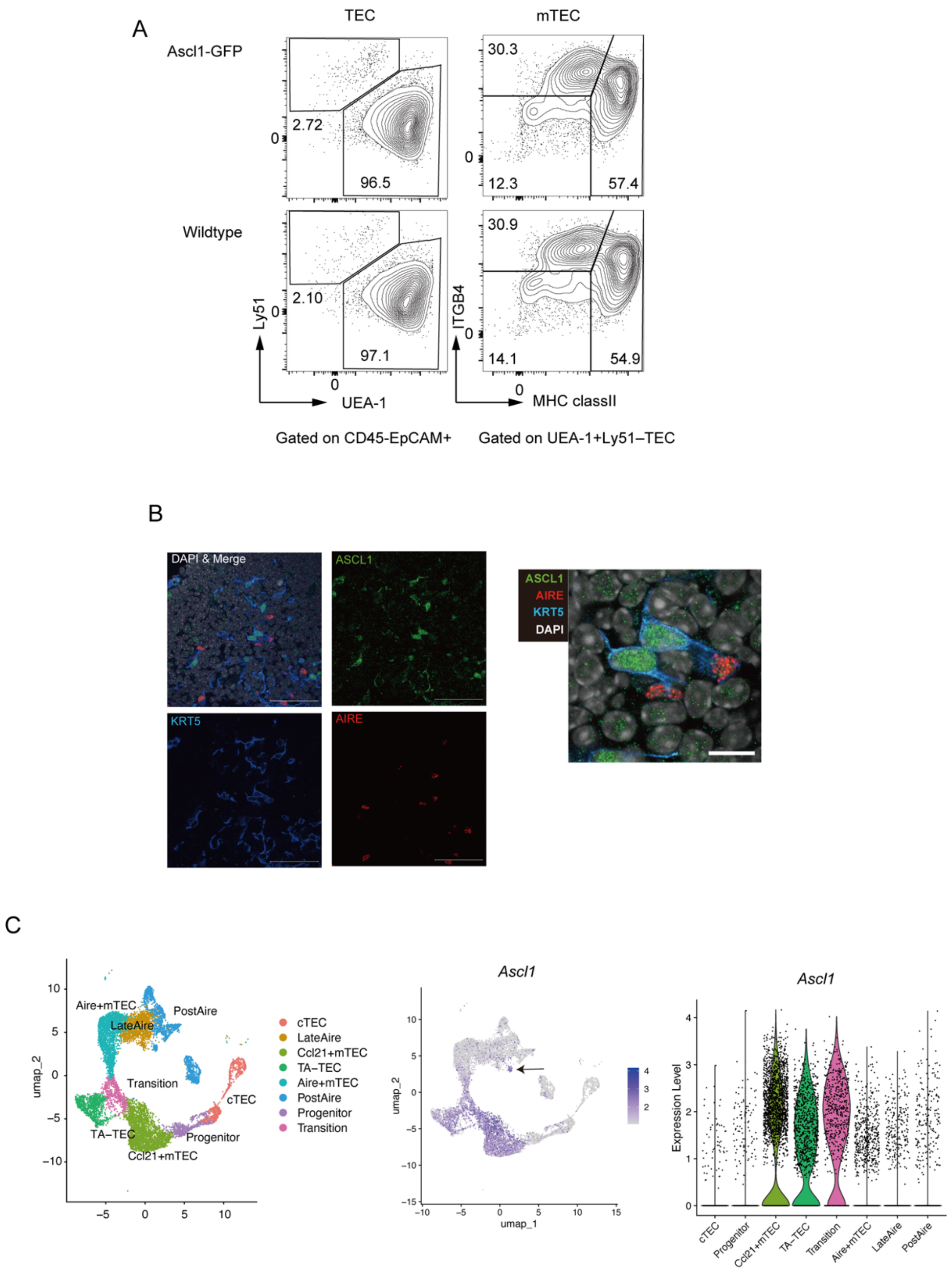


**Fig. S7. Expression of ASCL1 in TECs.**
A. Gating strategy of TECs in ASCL1-GFP mice.
B. Immunostaining of ASCL1 in the murine thymus. Scale Bar, left; 50μm, right; 10 μm.
C. Expression of *Ascl1* in murine TECs (4-week-old) in the previous scRNA-seq data set[11]. Assignment of TEC clusters, feature plots for *Ascl1* expression in TECs, and volcano plot for *Ascl1* expression in assigned mTEC clusters were shown. Arrow shows a population in PostAire mTECs expressing Ascl1, most likely Endo-TECs.

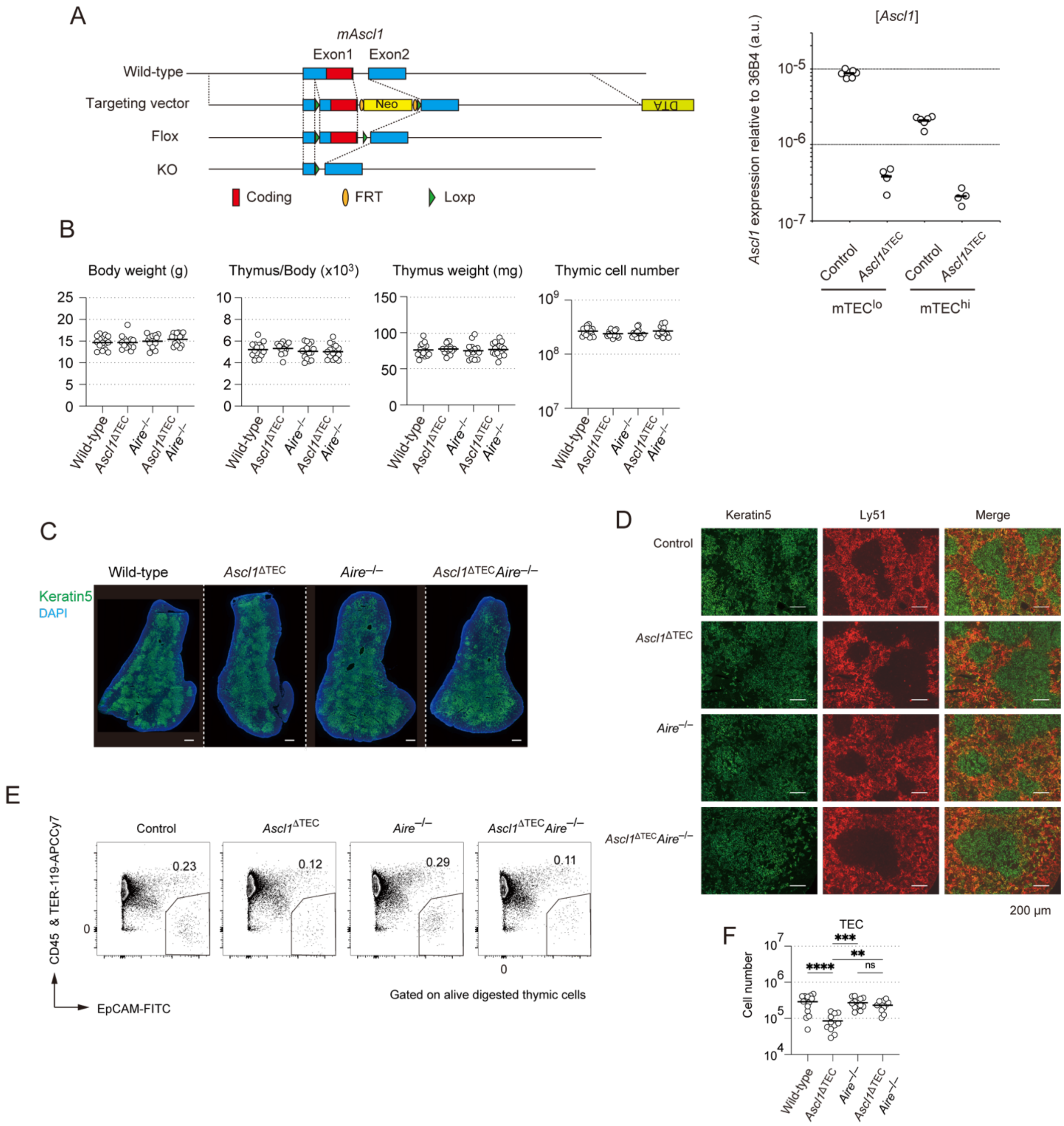

**Fig. S8. Establishment of TEC-specific ASCL1 deficient mice and immunohistochemical analyses of their thymic structures**

A. The genomic structure of the conditional allele of *Ascl1*. A gene-targeting vector contains two loxP sites (green triangle) in the non-coding region of exon 1 and the first intron. The right panel displays qPCR analysis of *Ascl1* gene expression in each mTEC subset of both TEC-specific ASCL1 deficient (*Ascl1*$^{\Delta TEC}$) and control mice to investigate the efficiency of Ascl1 reduction. Gating strategy is shown in Fig. 3B. mTEC$^{hi}$; CD45$^{-}$TER119$^{-}$EpCAM$^{+}$Ly51$^{-}$UEA-1$^{+}$CD80$^{hi}$, mTEC$^{lo}$; CD45$^{-}$TER119$^{-}$EpCAM$^{+}$Ly51$^{-}$UEA-1$^{+}$CD80$^{lo}$. Control (*Ascl1*$^{flox/flox}$); N=6, *Ascl1*$^{\Delta TEC}$ (Foxn1-Cre::*Ascl1* $^{flox/-}$); N=4.

B. Body weight, thymic mass and total thymic cell numbers of control (*Ascl1*$^{flox/flox}$), *Ascl1*$^{\Delta TEC}$ (Foxn1-Cre::*Ascl1* $^{flox/–}$), *Aire*$^{–/–}$, and *Ascl1*$^{\Delta TEC}$*Aire*$^{–/–}$ (Foxn1-Cre::*Ascl1* $^{flox/–}$ *Aire*$^{–/–}$) mice (N=15 per group).
C. Immunohistochemical staining of thymic sections from control, *Ascl1*$^{\Delta TEC}$, *Aire*$^{–/–}$, and *Ascl1*$^{\Delta TEC}$*Aire*$^{–/–}$ mice with a combination of anti-Keratin-5 (green) and DAPI staining (blue). scale bar: 500 μm.
D. Immunohistochemical staining of thymic sections from control, *Ascl1*$^{\Delta TEC}$, *Aire*$^{–/–}$, and *Ascl1*$^{\Delta TEC}$*Aire*$^{–/–}$ mice with a combination of anti-Keratin-5 (green) and Ly51 (red) antibodies. scale bar: 200 μm.
E. Gating strategy for TECs in alive digested thymic cells from 4-week-old control, *Ascl1*$^{\Delta TEC}$, *Aire*$^{–/–}$ and *Ascl1*$^{\Delta TEC}$*Aire*$^{–/–}$mice.
F. Cell number of total TECs of 4-week-old control, *Ascl1*$^{\Delta TEC}$, *Aire*$^{–/–}$ and *Ascl1*$^{\Delta TEC}$*Aire*$^{–/–}$ mice. Control (N=14), *Ascl1*$^{\Delta TEC}$ (N=11), *Aire*$^{–/–}$ (N=12) and *Ascl1*$^{\Delta TEC}$*Aire*$^{–/–}$ (N=11). *P<0.05, ***P<0.001; Tukey-Kramer test.

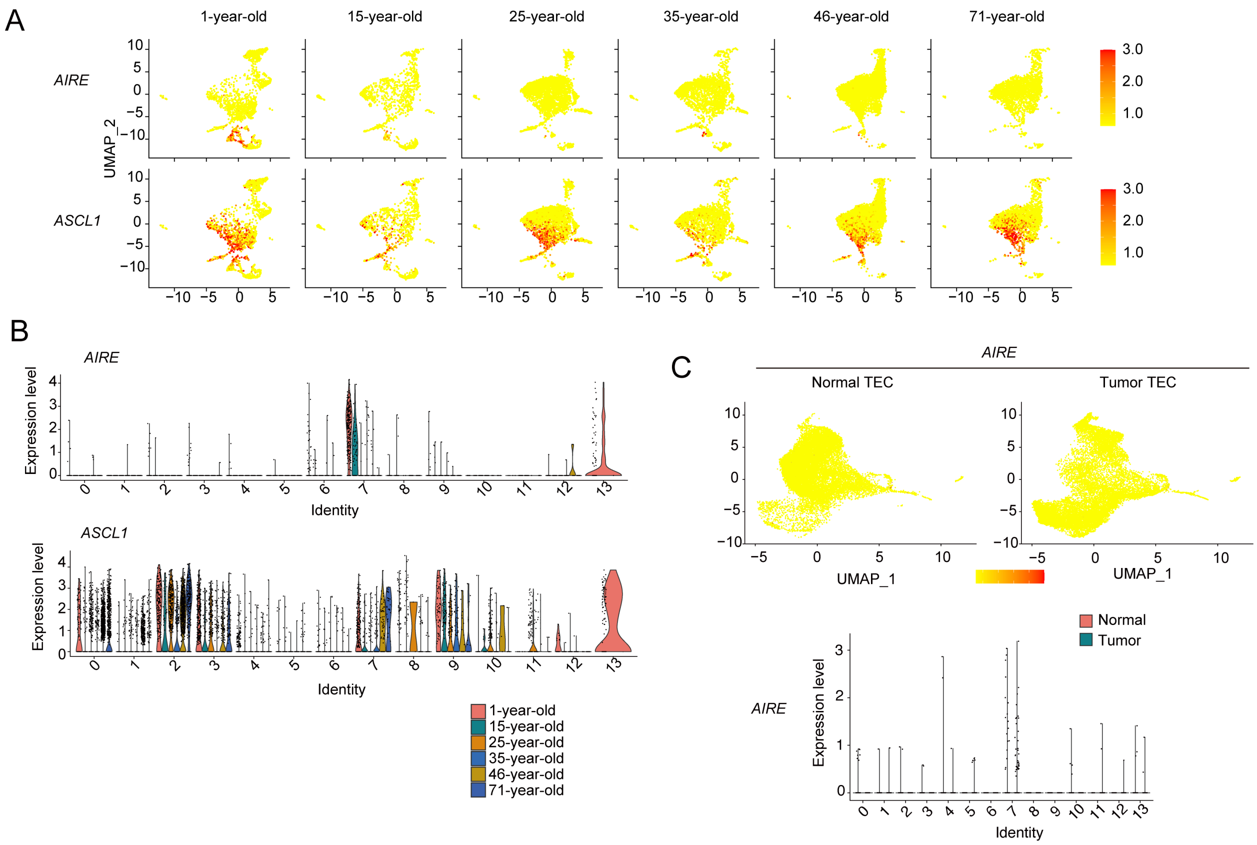


**Fig. S9. Aging-dependent down-regulation of AIRE gene expression, but not ASCL1, in normal human TECs**

A. Feature plots illustrate the expression levels of *AIRE* and *ASCL1* at indicated ages.

B. Violin plots illustrate the expression levels of *AIRE* and *ASCL1* at indicated ages.

C. Feature plots and a violin plot illustrate the expression levels of *AIRE* in normal TECs and tumor TECs

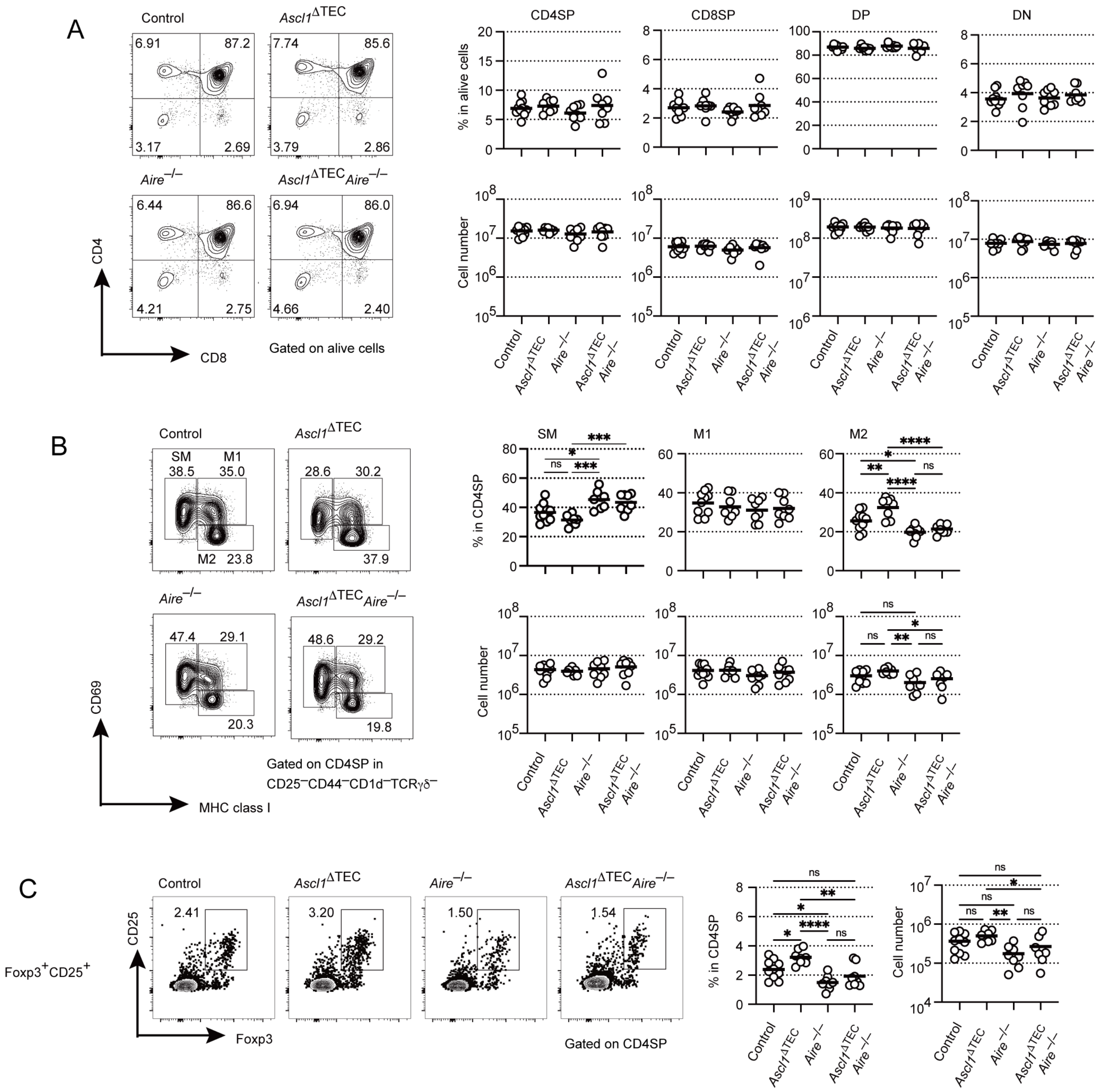


**Fig. S10. Flow-cytometric analysis of thymocytes.**

A. Representative FACS profiles for CD4 and CD8 expression from four-week-old mice. Cell percentages and cell numbers of $CD4^{+}CD8^{-}$ (CD4SP), $CD4^{-}CD8^{+}$ (CD8SP), $CD4^{+}CD8^{+}$(DP), and $CD4^{-}CD8^{-}$ (DN) in thymocytes. Control (N=10), *Ascl1*$^{\Delta TEC}$ (N=8), *Aire*$^{-/-}$(N=8), and *Ascl1*$^{\Delta TEC}$*Aire*$^{-/-}$ (N=8) mice at 4-week-old. Data are means (bars) and individual animals (circles). No significant difference was found by Tukey-Kramer test.

B. Flow cytometric analysis of thymocytes for $CD69^{+}MHCI^{-}$ (Semi Mature; SM), $CD69^{+}MHCI^{+}$ (Mature1; M1) or $CD69^{-}MHCI^{+}$ (Mature2; M2) cells in CD4SP with exclusion of $CD1d^{+}$NKT cells, $CD25^{+}$T reg cells, $CD44^{hi}$ recirculating memory cells and $GL3^{+}$ γδ T cells. Quantification of SM, M1 and M2 CD4SP in control (N=10), *Ascl1*$^{\Delta TEC}$ (N=8), *Aire*$^{-/-}$ (N=8), and

*Ascl1*$^{\Delta TEC}$*Aire*$^{-/-}$ (N=8) mice. Data are means (bars) and individual animals (circles). *P<0.05, **P<0.01, ***P<0.001; Tukey-Kramer test.
C. Flow cytometric analysis of Foxp3$^{+}$CD25$^{+}$ regulatory T cells from control, *Ascl1*$^{\Delta TEC}$, *Aire*$^{-/-}$, and *Ascl1*$^{\Delta TEC}$*Aire*$^{-/-}$ mice. Left: representative data of control, *Ascl1*$^{\Delta TEC}$, *Aire*$^{-/-}$, and *Ascl1*$^{\Delta TEC}$*Aire*$^{-/-}$. Right: percentages of regulatory T (Foxp3$^{+}$CD25$^{+}$) cells in CD4SP and cell numbers. Control (N=10), *Ascl1*$^{\Delta TEC}$ (N=8), *Aire*$^{-/-}$(N=8), and *Ascl1*$^{\Delta TEC}$*Aire*$^{-/-}$ (N=8). Data are means (bars) and individual animals (circles). *P<0.05, **P<0.01, ***P<0.001; Tukey-Kramer test.

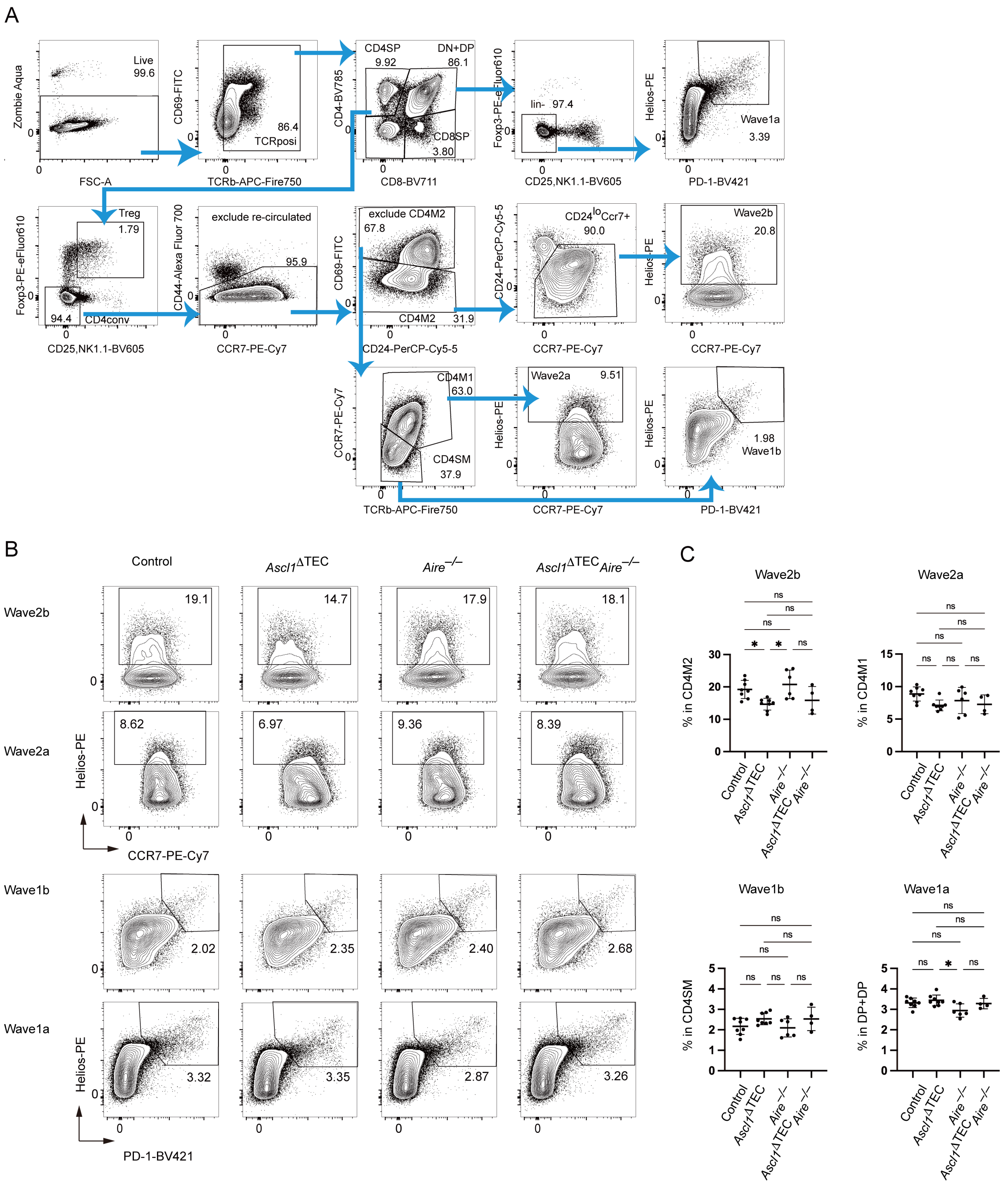


**Fig. S11. Flow cytometric analysis of TCR-dependent negative selection during thymocytes maturation from control and mutant mice.**

A. Flow cytometry gating strategy to separate different T-cell subtypes.

B. Representative profile of flow cytometric analysis of thymocytes from control, *Ascl1*$^{\Delta TEC}$, *Aire*$^{-/-}$, and *Ascl1*$^{\Delta TEC}$*Aire*$^{-/-}$ adult mice (age matched from 8 to14 week-old) .

C. Percentages of live CD4-single-positive thymocytes at stages of negative selection are shown. Data are means (bars) and individual animals (dots). Control (N=8), *Ascl1*$^{\Delta TEC}$ (N=8), *Aire*$^{-/-}$ (N=6), and *Ascl1*$^{\Delta TEC}$*Aire*$^{-/-}$ (N=4). *P<0.05, **P<0.01, ***P<0.001; Tukey-Kramer test.

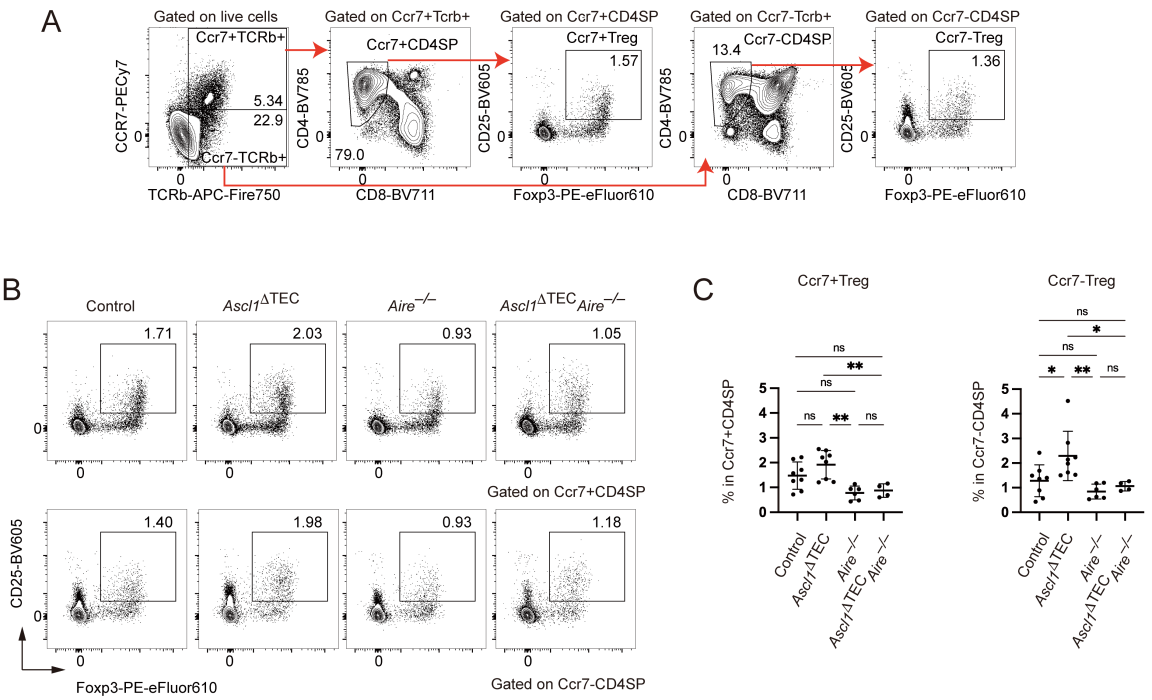


**Fig. S12. Flow cytometric analysis of Ccr7+ and Ccr7– regulatory T cells from control and mutant adult mice.**

A. Flow cytometry gating scheme showing Ccr7$^{+}$ and Ccr7$^{-}$ regulatory T cells in thymocytes.

B. Representative data of control (N=8), *Ascl1*$^{\Delta TEC}$ (N=8), *Aire*$^{-/-}$ (N=6), and *Ascl1*$^{\Delta TEC}$*Aire*$^{-/-}$ (N=4) adult mice (age matched from 8 to14 week-old).

C. Percentages of Ccr7$^{+}$ and Ccr7$^{-}$ regulatory T (Foxp3+CD25+) cells in CD4SP. Data are means (bars) and individual animals (dots). *P<0.05, **P<0.01, ***P<0.001; Tukey-Kramer test.

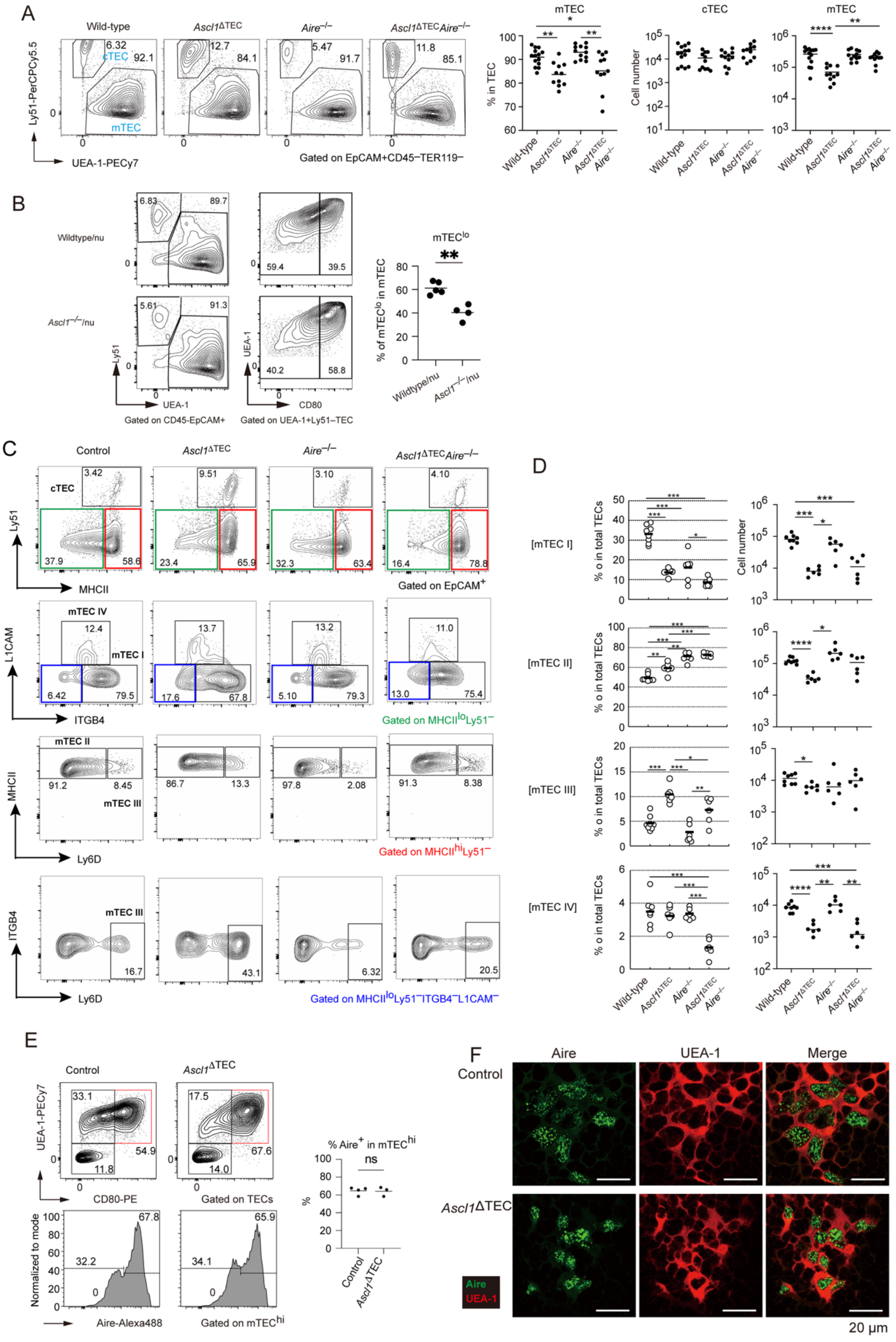
A
Wild-type
Ascl1ΔTEC
Aire−/−
Ascl1ΔTECAire−/−
cTEC
mTEC
Ly51-PerCPCy5.5
UEA-1-PECy7
Gated on EpCAM+CD45−TER119−
% in TEC
Cell number
B
Wildtype/nu
Ascl1−/−/nu
mTEClo
% of mTEClo in mTEC
Gated on CD45-EpCAM+
Gated on UEA-1+Ly51−TEC
UEA-1
CD80
Ly51
C
Control
MHCII
Gated on EpCAM+
mTEC IV
mTEC I
L1CAM
ITGB4
Gated on MHCIIloLy51−
mTEC II
mTEC III
Ly6D
Gated on MHCIIhiLy51−
Gated on MHCIIloLy51−ITGB4−L1CAM−
D
[mTEC I]
[mTEC II]
[mTEC III]
[mTEC IV]
% o in total TECs
E
UEA-1-PECy7
CD80-PE
Gated on TECs
Normalized to mode
Aire-Alexa488
Gated on mTEChi
% Aire+ in mTEChi
ns
F
Aire
UEA-1
Merge
Ascl1ΔTEC
20 µm

**Fig. S13. Flow cytometric analysis and immunostaining analysis of mTEC subsets.**
A. Flow-cytometric analysis of TECs from 4-week-old control, *Ascl1*$^{\Delta TEC}$, *Aire*$^{-/-}$ and *Ascl1*$^{\Delta TEC}$*Aire*$^{-/-}$mice. TEC were separated into mTEC and cTEC by expression of UEA-1 ligand and Ly51. Percentage of mTECs and the absolute number of mTEC and cTEC were summarized in right graphs.
B. Flow-cytometric analysis of TECs from control and *Ascl1*$^{-/-}$ thymi grafted into nude mice. TEC were separated into mTEC and cTEC by expression of UEA-1 ligand and Ly51. Percentage of mTEC$^{lo}$ in mTEC were summarized in right graphs.
C. Flow-cytometric analysis of mTECs from 4-week-old control, *Ascl1*$^{\Delta TEC}$, *Aire*$^{-/-}$ and *Ascl1*$^{\Delta TEC}$*Aire*$^{-/-}$mice. mTECs were separated into mTEC I, mTEC II, mTEC III, and mTEC IV by expression of MHCII, Ly51, ITGB4, L1CAM and Ly6D. mTEC I includes immature mTECs expressing CCL21, mTEC II mainly includes mature Aire$^{+}$mTECs. mTEC III and IV are regarded as post-Aire mTEC, the latter is tuft-like mTECs.
D. Ratio and absolute number of mTEC I, mTEC II, mTEC III, and mTEC IV in total TECs of control (N=8), *Ascl1*$^{\Delta TEC}$ (N=6), *Aire*$^{-/-}$(N=6), and *Ascl1*$^{\Delta TEC}$*Aire*$^{-/-}$ (N=6). Data are means (bars) and individual animals (circles). *P<0.05, **P<0.01, ***P<0.001; Tukey-Kramer test of each TEC ratio and Brown-Forsythe and Welch ANOVA test of cell numbers.
E. Flow-cytometric analysis of Aire-expressing TECs in the thymus of 4-week-old control and *Ascl1*$^{\Delta TEC}$ mice. Panel shows percentages of Aire$^{+}$ cells in mTEC$^{hi}$ (CD45$^{-}$TER119$^{-}$ EpCAM$^{+}$Ly51$^{-}$UEA-1$^{+}$CD80$^{hi}$). Control; N=4, *Ascl1*$^{\Delta TEC}$; N=3.
F. Immunohistochemical staining of thymic sections from control and *Ascl1*$^{\Delta TEC}$mice with a combination of anti-AIRE (green) and UEA-1-lectin (red). Scale bar: 20 μm.

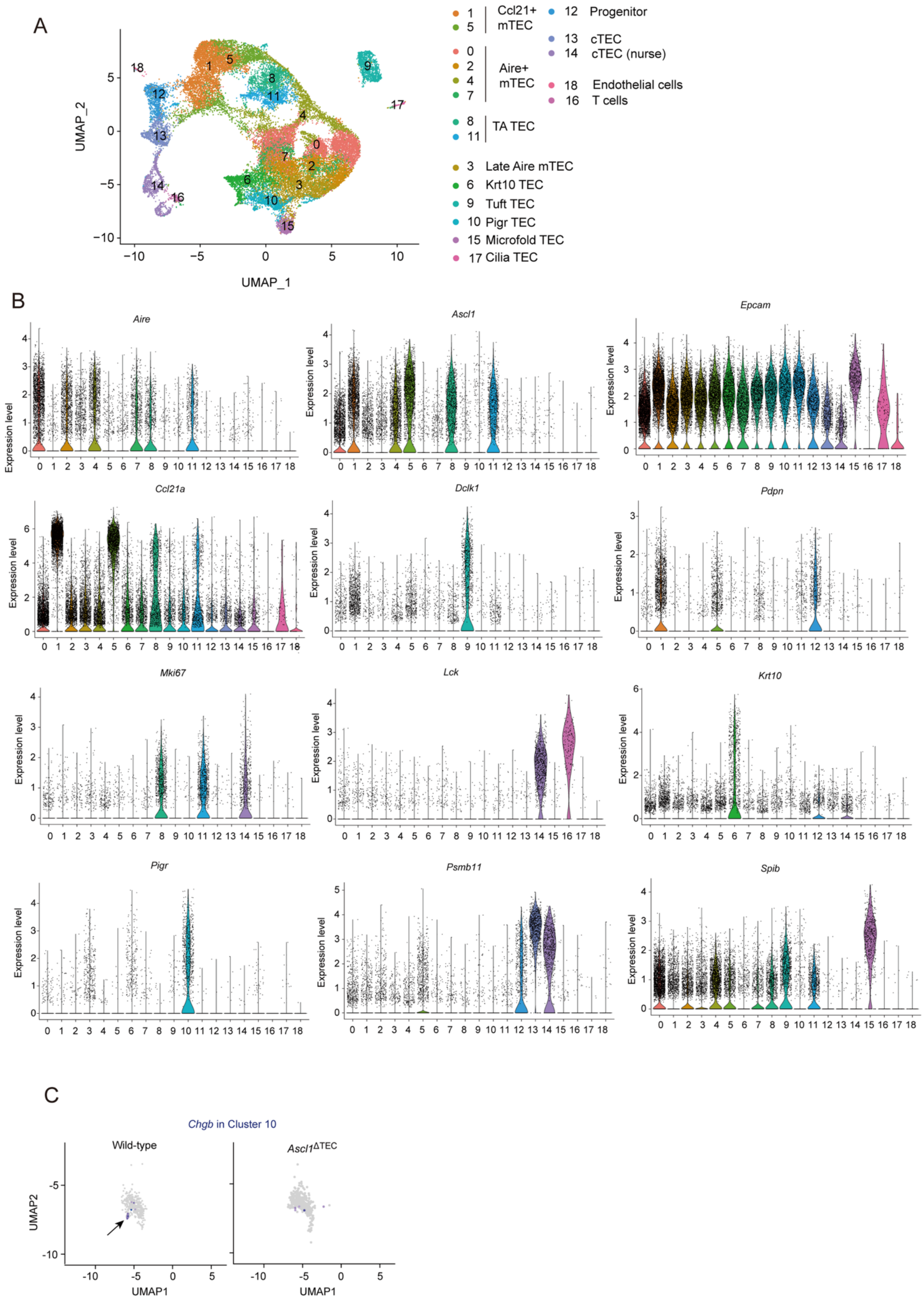
A
UMAP_1
UMAP_2
1 5 Ccl21+ mTEC
0 2 4 7 Aire+ mTEC
8 11 TA TEC
3 Late Aire mTEC
6 Krt10 TEC
9 Tuft TEC
10 Pigr TEC
15 Microfold TEC
17 Cilia TEC
12 Progenitor
13 cTEC
14 cTEC (nurse)
18 Endothelial cells
16 T cells
B
Aire
Ascl1
Epcam
Ccl21a
Dclk1
Pdpn
Mki67
Lck
Krt10
Pigr
Psmb11
Spib
Expression level
C
Chgb in Cluster 10
Wild-type
Ascl1^ΔTEC
UMAP1
UMAP2

**Fig. S14. Assignment of scRNA-seq clusters to mTEC subset in the mouse thymus.**
A. UMAP visualization of integrated scRNA-seq data of TECs from 4-week-old wild-type, *Ascl1*$^{\Delta\text{TEC}}$, *Aire*$^{-/-}$, and *Ascl1*$^{\Delta\text{TEC}}$*Aire*$^{-/-}$ mice. Clusters were assigned to each subset according to the marker gene expression.
B. Violin plots show the expression of marker genes in each cluster of the control dataset
C. Endo TEC marker *Chgb*-expressing cells in post-Aire mTEC cluster 10. Arrow indicates the *Chgb*-expressing cluster.

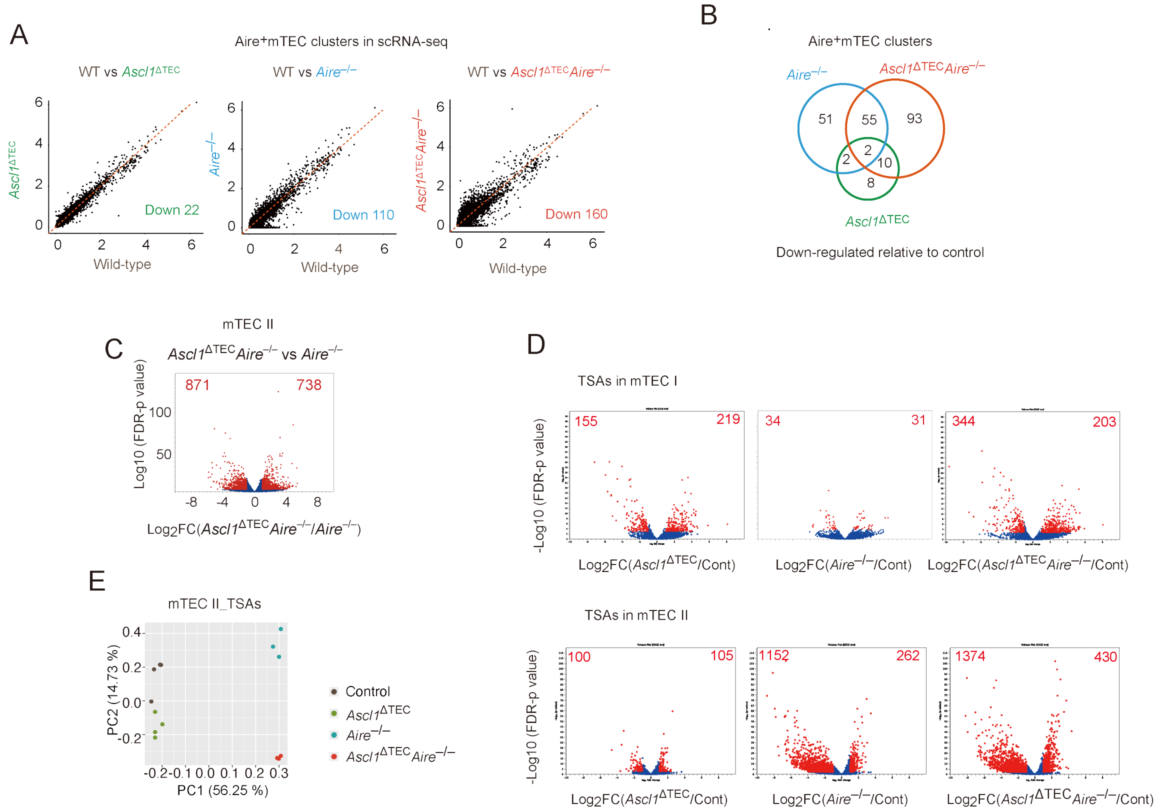


**Fig. S15. Gene expression analysis of mTECs.**

A. Scatterplot comparing gene expression levels in Aire$^{+}$ mTEC cluster between wild-type and *Ascl1*$^{\Delta TEC}$, *Aire*$^{-/-}$, or *Ascl1*$^{\Delta TEC}$*Aire*$^{-/-}$ mice in scRNA-seq. The numbers of down-regulated genes (log2FC < -0.25 and P < 0.05) are indicated in the panels.

B. Venn diagram compare the number of downregulated genes ($\log_2$FC < –1, P < 0.05) in Aire$^{+}$ mTECs between wild-type mice and *Ascl1*$^{\Delta TEC}$, *Aire*$^{-/-}$, or *Ascl1*$^{\Delta TEC}$*Aire*$^{-/-}$ mice in scRNA-seq analysis.

C. The volcano plot to illustrate differential gene expression from bulk-RNA-seq analysis in mTEC II of *Ascl1*$^{\Delta TEC}$*Aire*$^{-/-}$mice compared to *Aire*$^{-/-}$mice is shown. Red dots indicate significantly changed genes (FDR P<0.05, Fold-change >2 or < -2).

D. Volcano plots to illustrate differential TSA gene expression in mTEC I (upper) and mTEC II (lower) of *Ascl1*$^{\Delta TEC}$, *Aire*$^{-/-}$, and *Ascl1*$^{\Delta TEC}$*Aire*$^{-/-}$mice compared to control mice. For mTEC I, N=3 for all samples. For mTEC II, N=4 for control and *Ascl1*$^{\Delta TEC}$, and N=3 for *Aire*$^{-/-}$ and *Ascl1*$^{\Delta TEC}$*Aire*$^{-/-}$. Red points indicate significantly changed TSA genes (FDR P<0.05, Fold-change >2).

E. PCA analysis of TSA gene expression in mTEC II of control (N=4), *Ascl1*$^{\Delta TEC}$ (N=4), *Aire*$^{-/-}$ (N=3), and *Ascl1*$^{\Delta TEC}$*Aire*$^{-/-}$ mice (N=3) is shown.

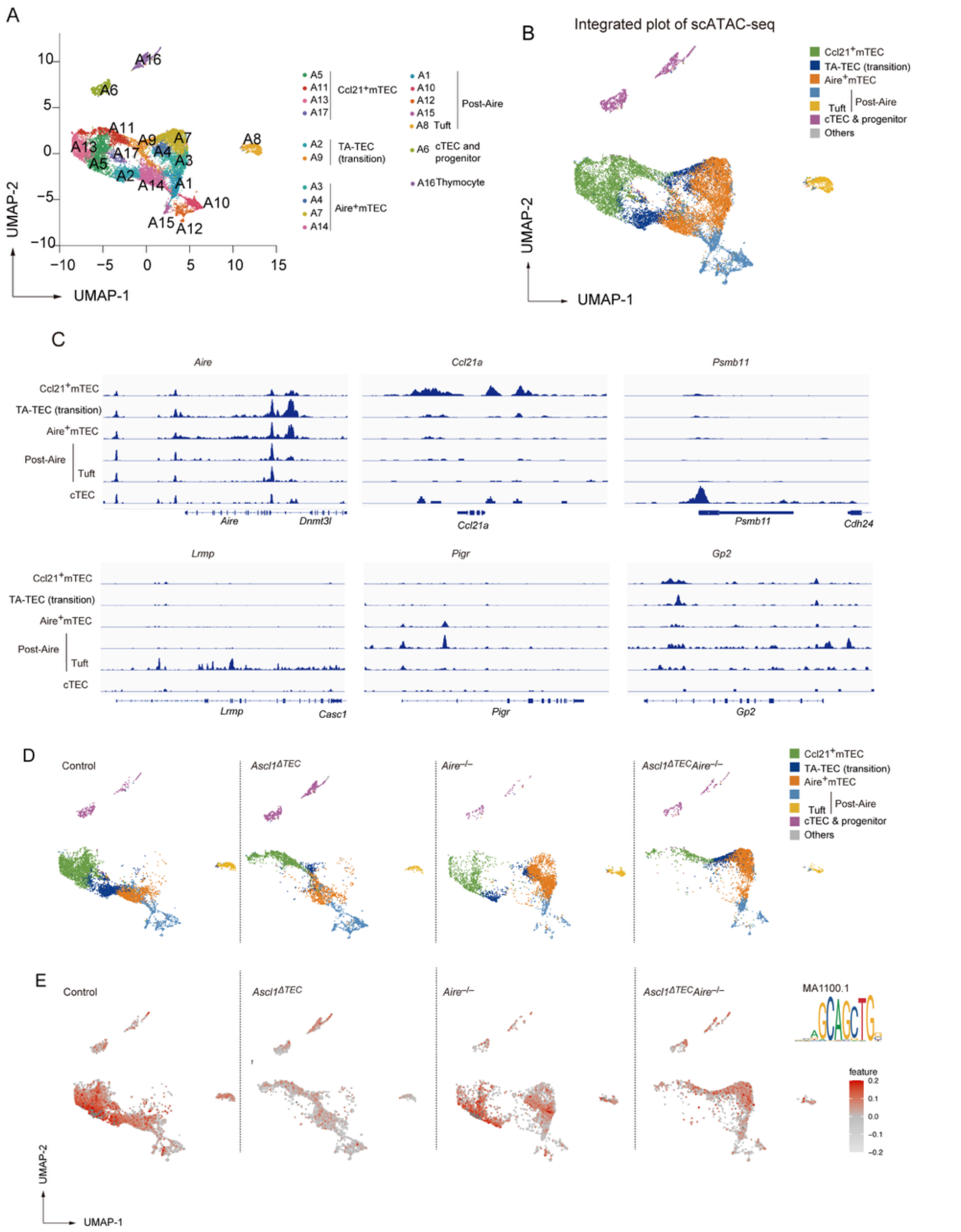

**Fig. S16. scATAC analysis of TECs.**

A. UMAP visualization of integrated scATAC-seq data of TECs from 4-week-old wild-type, *Ascl1*$^{\Delta TEC}$, *Aire*$^{-/-}$, and *Ascl1*$^{\Delta TEC}$*Aire*$^{-/-}$ mice. Clusters were assigned to each subset by the integrative analysis with scRNA-seq data of TECs.

B. Individual cells in the UMAP plot of scATAC-data (left) were assigned and transferred to the UMAP plot of scRNA-seq cluster according to the predicted gene expression profile from scATAC-seq data.

C. Tracks of ATAC peak for typical marker genes of mTEC subsets in scATAC-seq data.

D. The split UMAP plots based on their genotypes.

E. ASCL1 motif activity in each genotype.

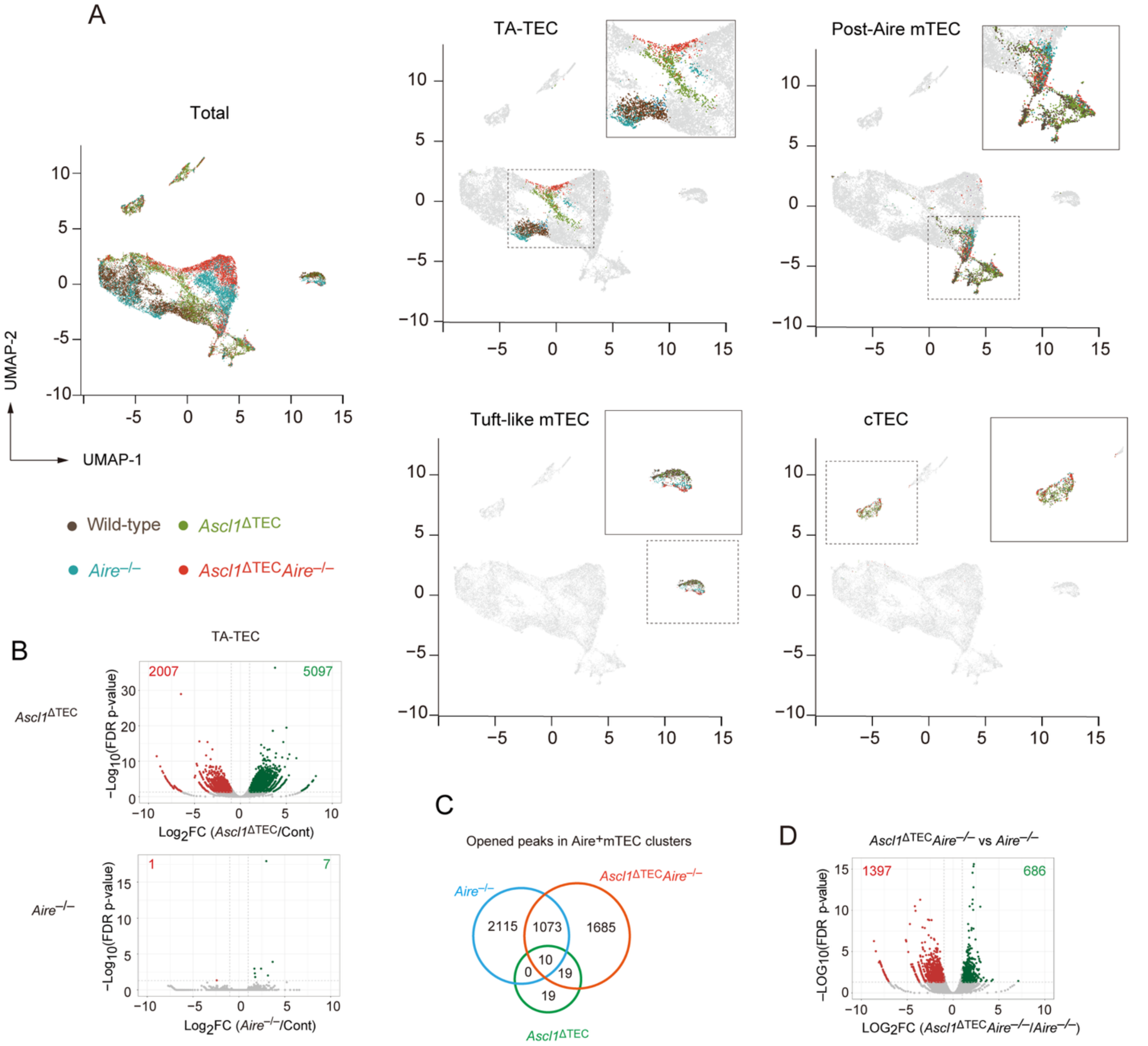


**Fig. S17. scATAC analysis of mTECs from the mutant mice.**

A. The UMAP plot for TEC scATAC data is color-coded according to the genotypes and further separated by TEC subsets (Post-Aire mTEC, Tuft-like mTEC, and cTEC). Enlarged inset views of individual TEC subsets are shown in the upper right corner of the UMAP plots.

B. Volcano plots to illustrate differentially opened (green) and closed (red) genomic regions in TA-TEC of *Ascl1*$^{\Delta TEC}$ (left panels) and *Aire*$^{-/-}$ (right panels) mice compared to control mice.

C. Venn diagram for ATAC peaks opened (FDR P < 0.05 and Fold change < -2) in Aire$^{+}$mTEC of *Ascl1*$^{\Delta TEC}$, *Aire*$^{-/-}$, and *Ascl1*$^{\Delta TEC}$*Aire*$^{-/-}$mice as compared to that of control mice.

D. Volcano plots to illustrate differentially opened (green) and closed (red) genomic region of TECs from *Aire*$^{-/-}$ *Ascl1*$^{\Delta TEC}$ compared to and *Aire*$^{-/-}$mice.

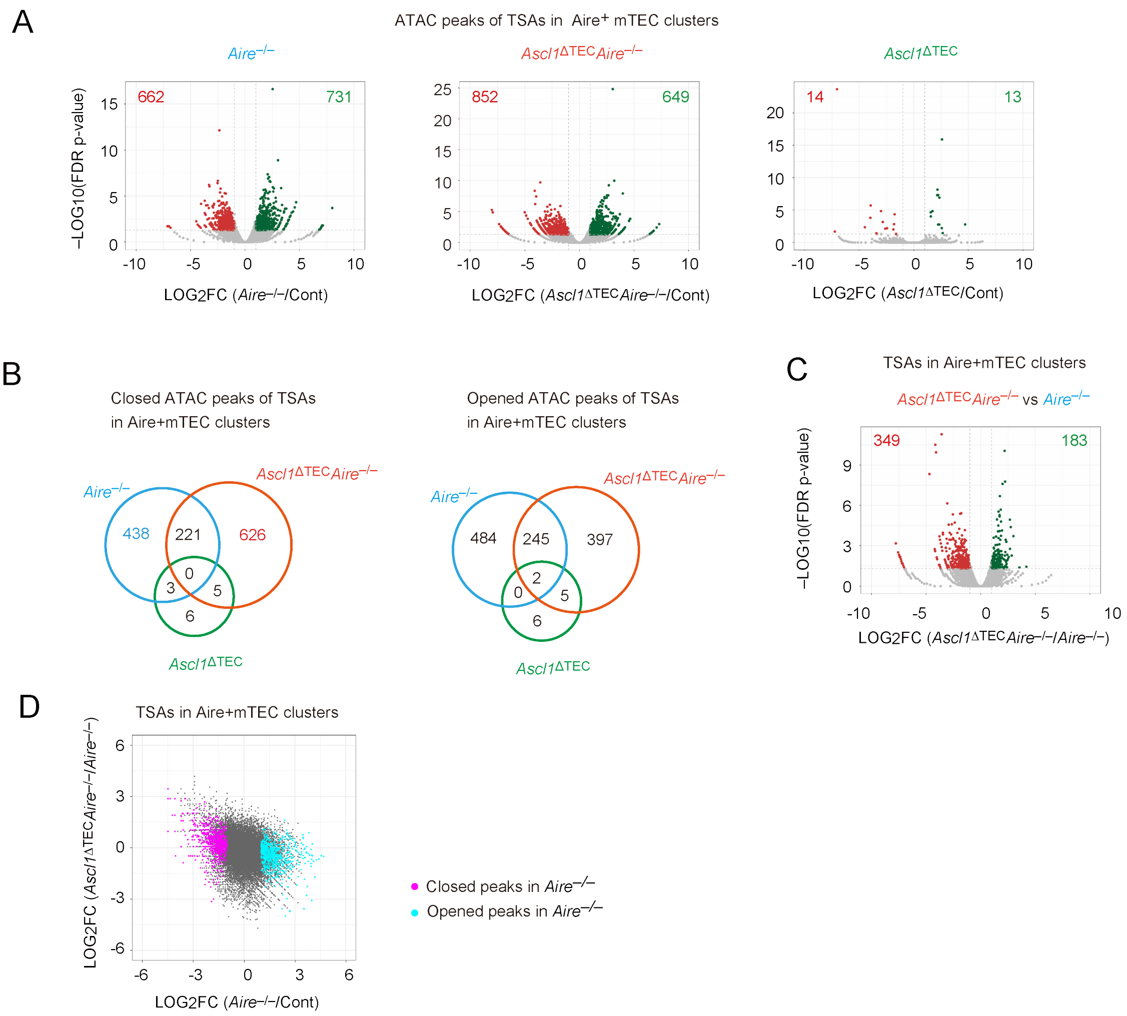


**Fig. S18. Influences of *Ascl1* and *Aire* deletion in chromatin accessibility of TSA genes of TECs.**

A. Volcano plots illustrate differential open (green) and closed (red) genomic regions of TSAs in Ccl21$^{+}$ mTEC and Aire$^{+}$ mTEC of *Aire*$^{-/-}$, *Ascl1*$^{\Delta TEC}$*Aire*$^{-/-}$, and *Ascl1*$^{\Delta TEC}$ mice compared to control mice.

B. Venn diagrams for ATAC peaks closed (FDR P < 0.05 and Fold change > 2) and open (FDR P < 0.05 and Fold change < -2) in Aire$^{+}$ mTEC of *Ascl1*$^{\Delta TEC}$, *Aire*$^{-/-}$, and *Ascl1*$^{\Delta TEC}$*Aire*$^{-/-}$mice as compared to that of control mice.

C. Volcano plots to illustrate differentially opened (green) and closed (red) genomic region of TSAs in Aire$^{+}$ mTECs from *Aire*$^{-/-}$ *Ascl1*$^{\Delta TEC}$ compared to and *Aire*$^{-/-}$mice.

D. A scatter plot comparing log2 fold changes of ATAC peaks of TSA regions in Aire$^{+}$mTEC cluster between *Aire*$^{-/-}$ versus control and *Ascl1*$^{\Delta TEC}$*Aire*$^{-/-}$ versus *Aire*$^{-/-}$. Pink dots indicate peaks significantly closed in Aire$^{+}$mTEC of *Aire*$^{-/-}$. Blue dots indicate peaks significantly opened in Aire$^{+}$mTEC of *Aire*$^{-/-}$ mice.

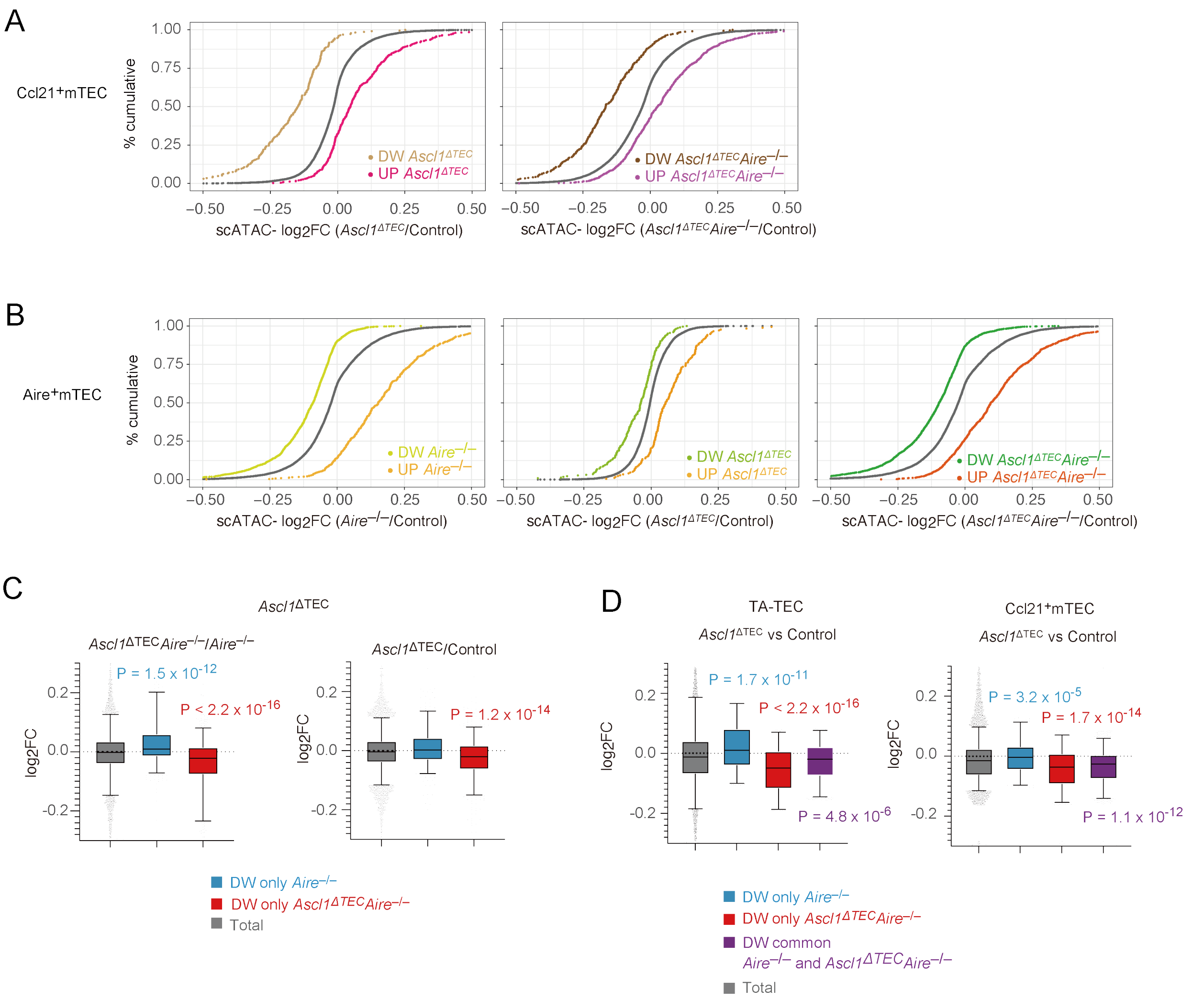


**Fig. S19. Cumulative distribution function plots in ATAC gene activity.**

A. The plots in ATAC activity in Ccl21$^{+}$mTEC for *Ascl1*$^{\Delta TEC}$ vs. control and *Ascl1*$^{\Delta TEC}$*Aire*$^{-/-}$ vs. control.

B. The plots in ATAC activity in Aire$^{+}$mTEC for *Aire*$^{-/-}$ vs. control, *Ascl1*$^{\Delta TEC}$ vs. control and *Ascl1*$^{\Delta TEC}$*Aire*$^{-/-}$ vs. control.

C. The box plots of cumulative distribution function plot for log2 fold change in 'ATAC gene activity' in Aire$^{+}$mTEC for *Ascl1*$^{\Delta TEC}$*Aire*$^{-/-}$ vs. *Aire*$^{-/-}$ (left) and *Ascl1*$^{\Delta TEC}$ vs. control (right) to investigate the correlation between scATAC-seq data and bulk RNA-seq data (Fig. 4D, Fig 5F). The ATAC gene activity is computed by adding the ATAC peaks within both the gene body and the 2kbp upstream region of the transcriptional start site. The ATAC gene activity of genes down-regulated only in *Aire*$^{-/-}$ mTEC II and that of genes down-regulated only in *Ascl1*$^{\Delta TEC}$*Aire*$^{-/-}$ mTEC II are plotted as blue and red dots, respectively. Gray dots are the total detected genes. P; Mann-Whitney U test.

D. The box plots of cumulative distribution function plot for log2 fold change in 'ATAC gene activity' in TA-TEC (left) or Ccl21$^{+}$ mTEC (right) for *Ascl1*$^{\Delta TEC}$ vs. control (Fig 5H). The ATAC gene activity of genes down-regulated only in *Aire*$^{-/-}$ mTEC II (blue), that of genes down-regulated only in *Ascl1*$^{\Delta TEC}$*Aire*$^{-/-}$ mTEC II (red) and that of genes commonly down-regulated both *Aire*$^{-/-}$ and *Ascl1*$^{\Delta TEC}$*Aire*$^{-/-}$ mTEC II (purple). Gray dots are the total detected genes. P; Mann-Whitney U test.

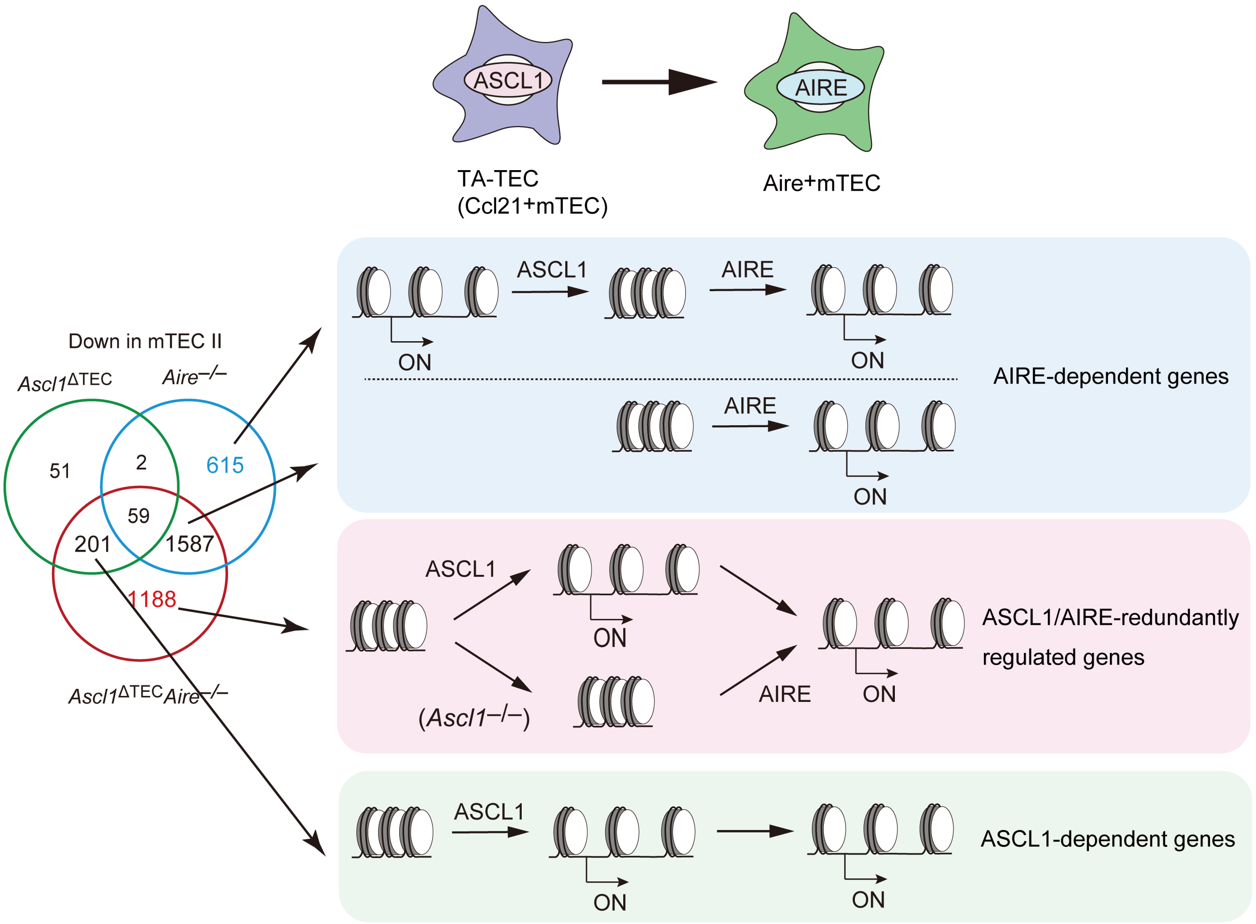


**Fig. S20. Proposed mechanism of the interplay between ASCL1 and AIRE in the regulation of gene expression in Aire$^{+}$mTECs.**
ASCL1 initially modulates chromatin structure in precursor mTECs. Some regions opened by ASCL1 remain accessible in mature mTECs and support gene expression. In Ascl1$^{-/-}$ mice, a subset of these regions is instead opened by AIRE, resulting in genes that become redundantly regulated by both factors. Conversely, chromatin regions that ASCL1 keeps closed—or never opens—are later opened by AIRE in mature mTECs, corresponding to AIRE-dependent gene regulation.

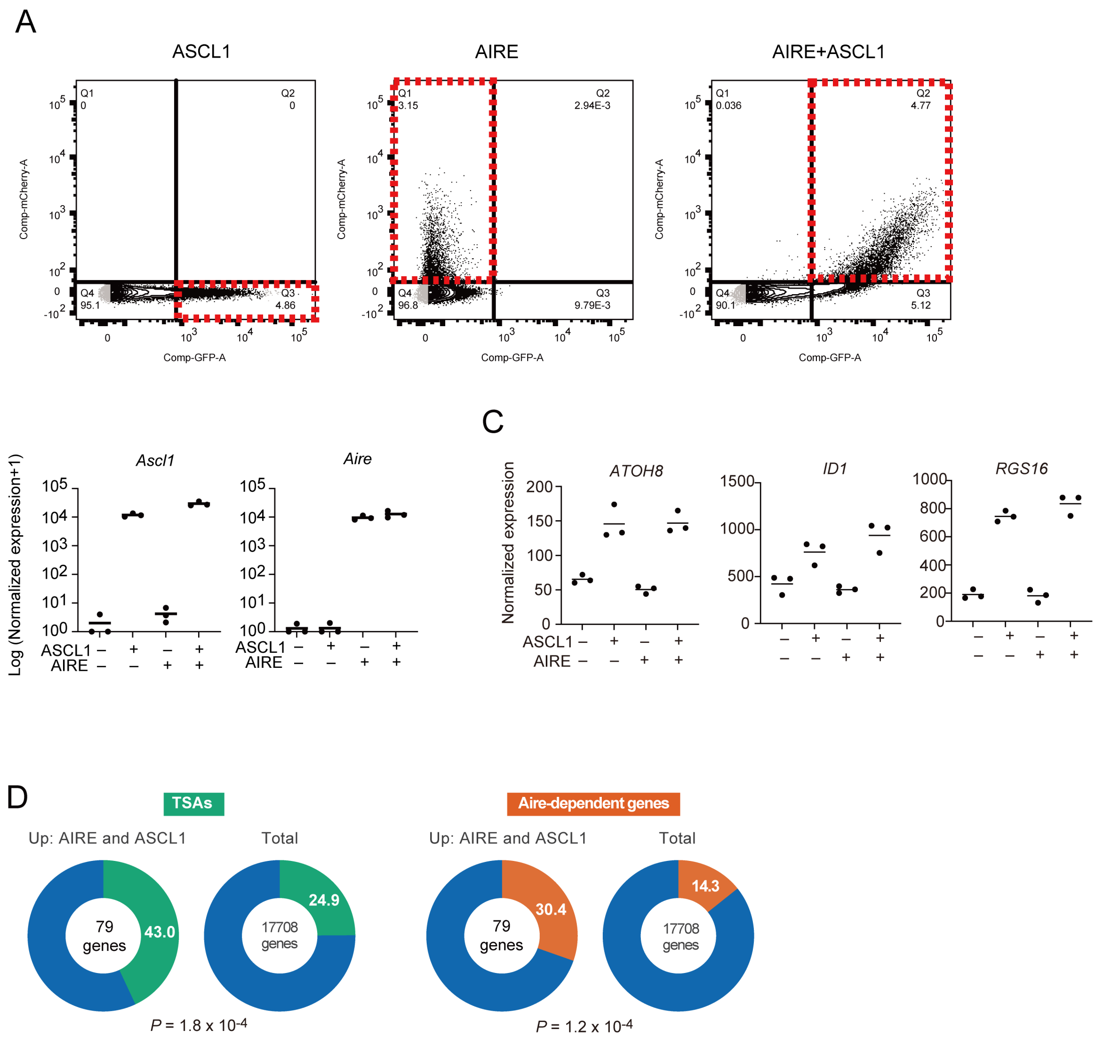


**Fig. S21. Gene expression changes in HEK293 cells following transfection with AIRE and ASCL1**

A. Gating strategy of sorting cells transfected with expression vector of ASCL1, AIRE, and its combination. IRES-GFP and IRES mCherry was used to detect the transfected cells.

B. Expression of Ascl1 and Aire in transfected cells.

C. Normalized expression levels of representative genes in HEK293 cells transfected with ASCL1, AIRE, or both (N=3 each).

D. Donut charts showing the proportion of Aire-dependent gene sets (human orthologs of mouse Aire-dependent genes) and tissue-specific antigens (TSAs; human orthologs of mouse TSAs) among genes upregulated by AIRE and ASCL1 transfection, or among all expressed genes in HEK293 cells. P values indicate the statistical significance of enrichment for Aire-dependent genes or TSAs.

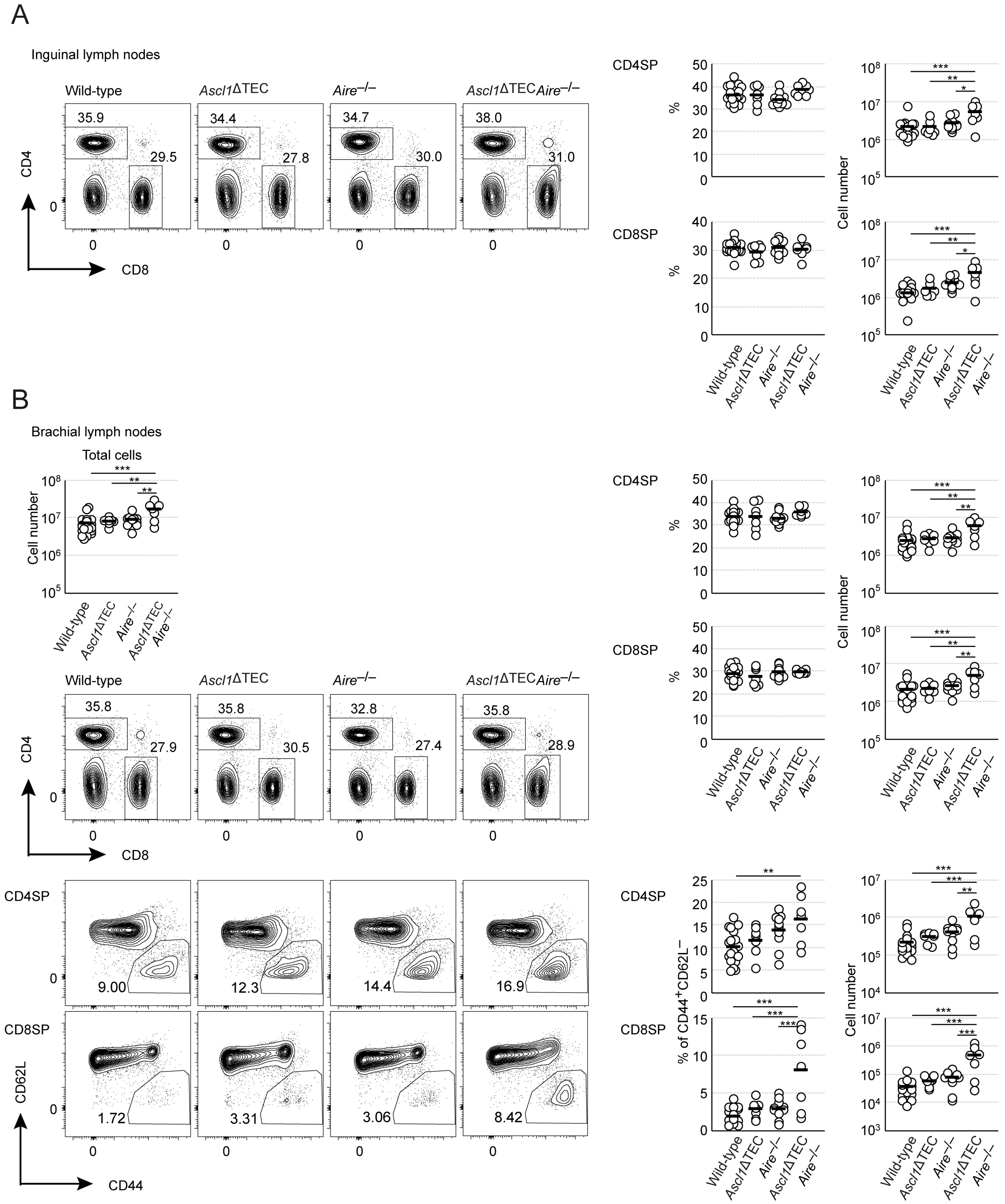


**Fig. S22. Flow cytometric analysis of peripheral organs from control and mutant mice.** A. Representative data of flow cytometric analysis of inguinal lymph nodes from 20-week-old control, *Ascl1*$^{\Delta TEC}$, *Aire*$^{-/-}$, and *Ascl1*$^{\Delta TEC}$*Aire*$^{-/-}$ mice. Distributions of CD4- and CD8-positive cells in lymphocytes are shown on the left. Control (n = 19), *Ascl1*$^{\Delta TEC}$ (N=7), *Aire*$^{-/-}$ (N=10), and *Ascl1*$^{\Delta TEC}$*Aire*$^{-/-}$ (N=7). Percentages and cell numbers of CD4-, or CD8-single-positive T

cells in inguinal lymph nodes are shown on the right. Data are means (bars) and individual animals (circles). *P<0.05, **P<0.01, ***P<0.001; Tukey-Kramer test.
B. Representative data of flow cytometric analysis of brachial lymph nodes from 20-week-old control, *Ascl1*$^{\Delta TEC}$, *Aire*$^{-/-}$, and *Ascl1*$^{\Delta TEC}$*Aire*$^{-/-}$ mice. Distributions of CD4- and CD8-positive cells, profiles of memory T cells in brachial lymphocytes are shown on the left. Control (N=19), *Ascl1*$^{\Delta TEC}$ (N=7), *Aire*$^{-/-}$ (N=10), and *Ascl1*$^{\Delta TEC}$*Aire*$^{-/-}$ (N=7). Percentages and cell numbers of CD4- and CD8-positive cells, memory T cells in brachial lymph nodes are shown on the right. Data are means (bars) and individual animals (circles). **P<0.01, ***P<0.001; Tukey-Kramer test.

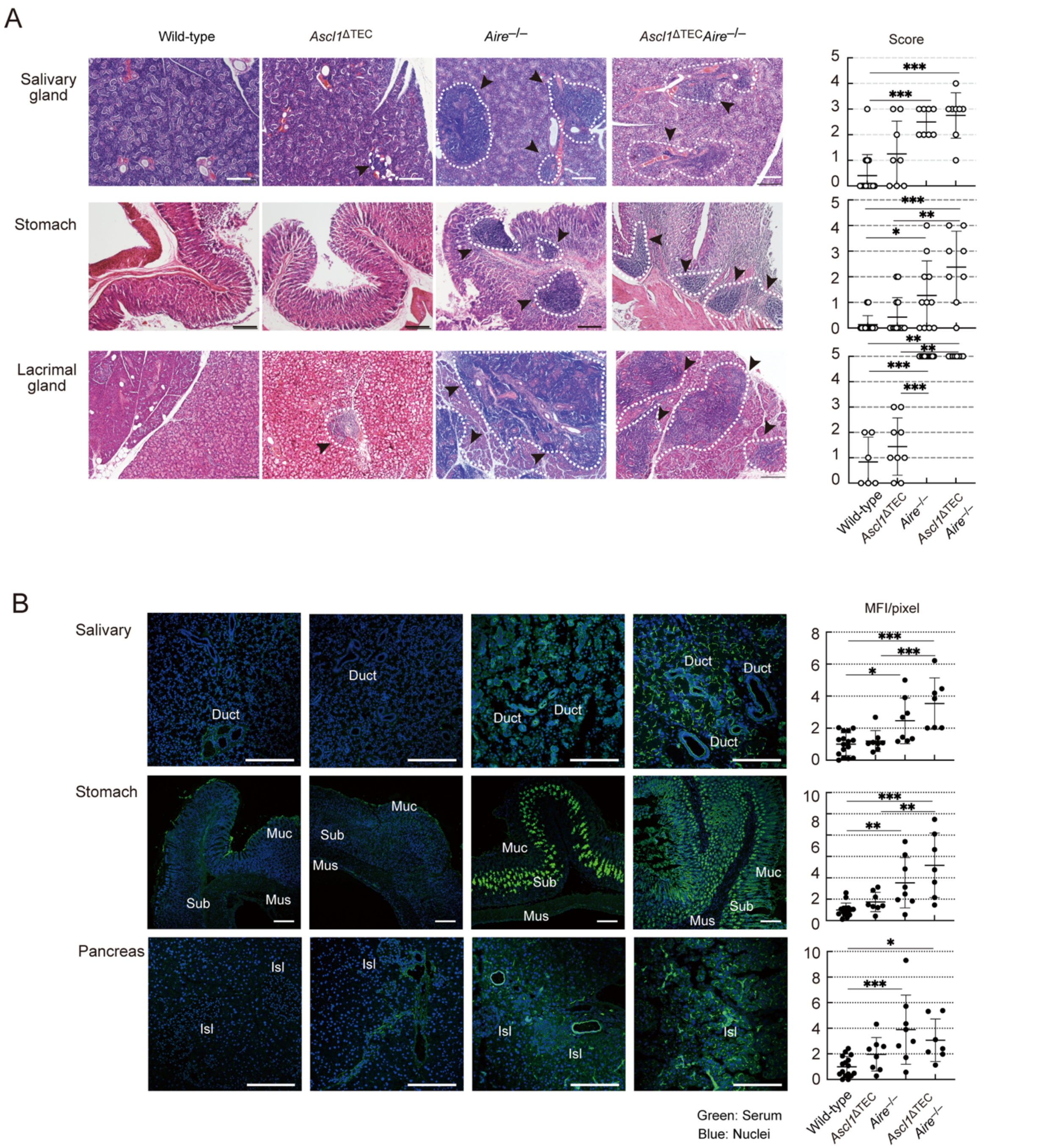


**Fig. S23. Autoimmunity caused by Ascl1 deficiency, Aire deficiency, or the combined deficiency of both.**

A. Histological analysis of some organs from control and mutant mice at 20 weeks, H&E staining for infiltrating lymphocytes. White dotted lines and arrows: inflammatory cell infiltration. Left: representative sample data. For salivary gland; Control (N = 15), *Ascl1*$^{\Delta TEC}$ (N=8), *Aire*$^{-/-}$ (N=8), and *Ascl1*$^{\Delta TEC}$*Aire*$^{-/-}$ (N=8), stomach; Control (N = 15), *Ascl1*$^{\Delta TEC}$ (N=14), *Aire*$^{-/-}$ (N=12), and *Ascl1*$^{\Delta TEC}$*Aire*$^{-/-}$ (N=8), lachrymal gland; Control (N = 6), *Ascl1*$^{\Delta TEC}$ (N=9), *Aire*$^{-/-}$ (N=12), and *Ascl1*$^{\Delta TEC}$*Aire*$^{-/-}$ (N=8).Bars, 200 µm. Right: infiltration scores. *P<0.05, **P<0.01, ***P<0.001; Steel-Dwass test.

B. Detection of autoantibodies. Organs from *Rag1*$^{-/-}$ mice were stained with sera (green) from control (n = 15), *Ascl1*$^{\Delta TEC}$ (N=8), *Aire*$^{-/-}$ (N=8), and *Ascl1*$^{\Delta TEC}$*Aire*$^{-/-}$ (N=7) mice at 20 weeks and DAPI (blue); Stomach, Mus: muscle layer, Sub: submucosa, Muc: mucosal layer. Pancreas, Isl; pancreatic islets. Scale bars, 200 µm. Right plots, relative MFI/pixel of autoantibodies, points, individual animals. *P<0.05, **P<0.01, ***P<0.001; Tukey-Kramer test.

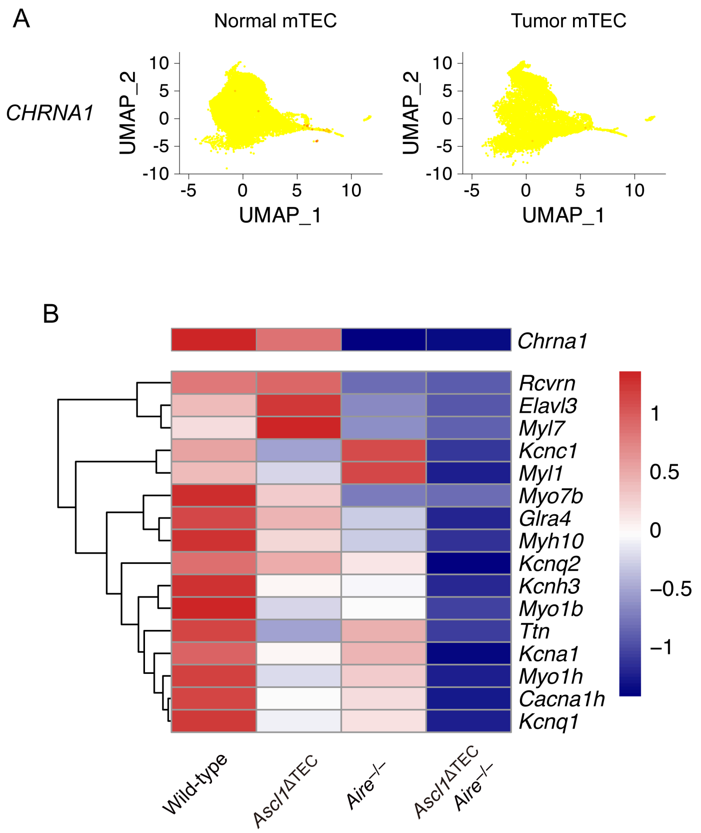


**Fig. S24. Expression of Chrna1 and possible MG antigen genes in thymoma and mouse models.**

A. Feature plots illustrate the expression levels of *CHRNA1,* a gene encoding AchR, in normal TECs and tumor TECs from patient thymus.

B. Heatmap of mouse orthologues of potential MG autoantigen genes that are down-regulated in mutant versus control mice.

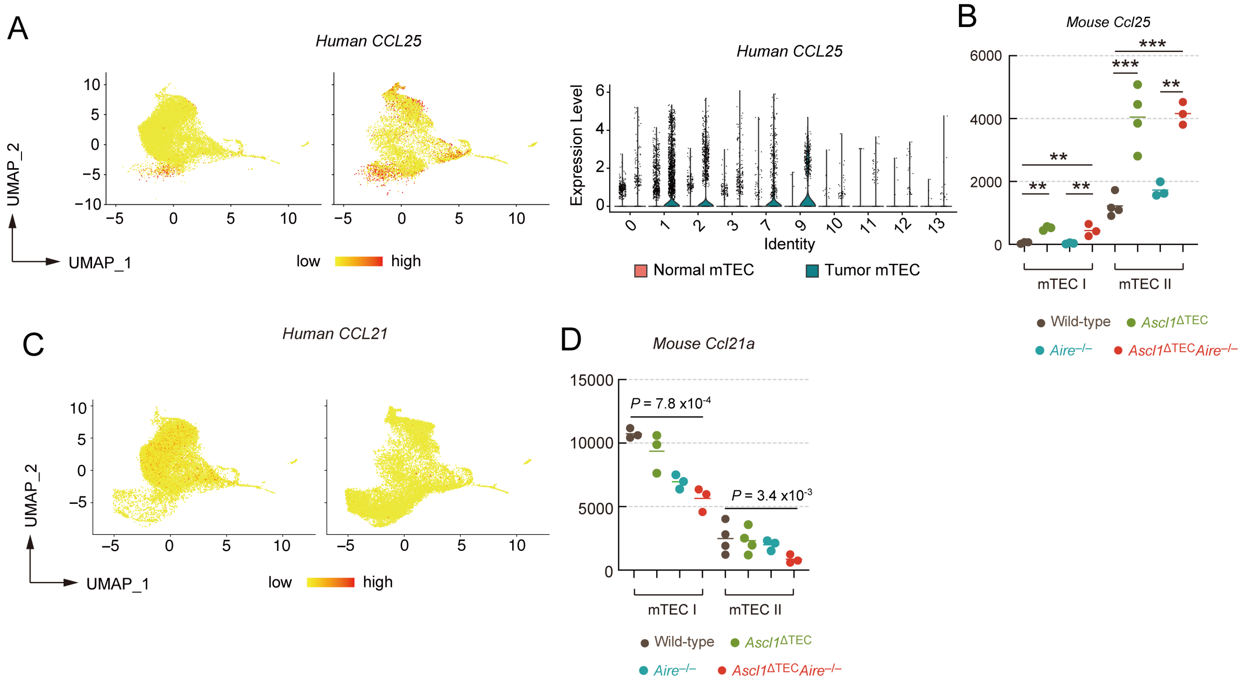


**Fig. S25. Increment of cTEC-associated genes and reduction of mTEC chemokines in human tumor TEC and TEC deficient in ASCL1 and AIRE**

A. UMAP projection highlighting mTECs expressing human CCL25 in normal and tumor mTECs (left). Violin plots show expression levels of CCL25 in normal versus tumor mTEC clusters (Figure 1A) from human patients (right).

B. Normalized expression values of mouse Ccl25 in mTEC I and mTEC II of control, *Ascl1*$^{\Delta\mathrm{TEC}}$, *Aire*$^{-/-}$, and *Ascl1*$^{\Delta\mathrm{TEC}}$*Aire*$^{-/-}$ mice in bulk RNA-seq analysis (Fig. 4B).

C. UMAP projection highlighting mTECs expressing human CCL21 in normal and tumor mTECs from scRNA-seq (Fig. 1).

F. Normalized expression values of mouse Ccl21a in mTEC I and mTEC II of control, *Ascl1*$^{\Delta\mathrm{TEC}}$, *Aire*$^{-/-}$, and *Ascl1*$^{\Delta\mathrm{TEC}}$*Aire*$^{-/-}$ mice in bulk RNA-seq (Fig. 4B)
